# A panel of near-isogenic lines derived from locally adapted populations of a wild plant: A powerful tool for dissecting additive and non-additive effects on ecologically important traits

**DOI:** 10.64898/2025.12.03.692110

**Authors:** Samuel J. Mantel, Gwonjin Lee, Juan Diego Rojas-Gutierrez, Brian J. Sanderson, M. Inam Jameel, Patrick Woods, Brian P. Dilkes, John K. McKay, Jon Ågren, Christopher G. Oakley

## Abstract

Identifying and estimating the effects of loci contributing to natural variation in ecologically important traits can be hampered by quantitative inheritance, dominance, epistasis, and environmentally dependent trait expression. Here we announce the availability of germplasm and sequence data for a reciprocal panel of near-isogenic lines (NILs) derived from locally adapted natural populations of *Arabidopsis thaliana,* for investigating the genetic basis of ecologically important traits. We created a panel of 54 NILs and performed whole genome sequencing to precisely locate introgression segments(s) in each NIL. Deep sequencing largely confirmed prior knowledge of NIL genotypes but also identified multiple novel small introgressions and regions of residual heterozygosity. To illustrate the utility of this panel, we identified genomic regions underlying ecotypic differences in flowering time in a laboratory common garden experiment. We detected strong additive effects on flowering time in multiple NILs with segments at the top of chromosome 5 implicating the floral regulator *FLC*, as expected based on previous quantitative trait locus studies. We also detected novel and complex contributions to ecotypic differences in flowering time, only visible in the NILs, suggesting the possibility of epistasis. Our results highlight the utility of this panel for dissecting the genetic architecture of ecologically important traits, including the future potential for fine mapping of additive effects and testing for epistasis and linkage using NILs derived from the panel. This panel can be used by any member of the research community to investigate any of a broad suite of traits for which the parents differ.

## Introduction

Local adaptation of natural populations is commonly observed (Kawecki & Ebert, 2004; Leimu & Fischer, 2008; Hereford, 2009), and is likely the result of selection imposed by heterogenous environmental conditions. However, there are few examples where the causal variants underlying ecotypic divergence in phenotypes that contribute to local adaptation have been identified (Dittmar *et al*., 2016; VanWallendael *et al*., 2019; Wadgymar *et al*., 2022). Ecologically important traits are often quantitatively inherited (Orr, 1998; Ungerer, 2005; Stinchcombe & Hoekstra, 2008; Mackay & Anholt, 2024), making it difficult to detect and estimate the additive effects of individual loci. Genetic interactions including dominance (Wardyn *et al*., 2007; Bouvet *et al*., 2009; Leinonen *et al*., 2013), epistasis (Phillips, 2008; Li *et al*., 2014b; Mackay & Anholt, 2024), and genotype-by-environment interactions (Anderson *et al*., 2011; Des Marais *et al*., 2013; Josephs, 2018) present additional challenges to detection and accurate effect estimation. Even when quantitative trait loci (QTL) are identified, candidate genes are seldom functionally validated (Stinchcombe & Hoekstra, 2008; Bomblies & Peichel, 2022, but see Lee *et al*., 2024). This is in part because of the effort inherent in constructing materials capable of testing the effects of a single-locus, and the large number of genes typically contained in a QTL region. Thus, genetic resources that enable efficient fine mapping of causal variants and allow estimation of both additive and non-additive genetic effects across spatially and temporally replicated experiments are extremely valuable.

Near-isogenic lines (NILs, also known by other names, see Jeuken & Lindhout, 2004; Lippman *et al*., 2007; Zhao *et al*., 2009; Shuang *et al*., 2023) represent one such genetic resource. As homozygous lines, NILs are “immortal” in the sense that they are easily maintained through selfing. They need only be genotyped once and can be replicated readily within and between experiments. Other popular mapping populations, such as F_2_ populations (Fishman *et al*., 2014; Marshall *et al*., 2020; Schluter *et al*., 2021), are easier to produce but do not allow replication of individual genotypes and require genotyping the individuals for each experiment. Recombinant inbred line populations (RILs; Wang *et al*., 2022; Oakley *et al*., 2023; Mantel & Sweigart, 2024) are also frequently used and “immortal”, allowing for simple replication of genotypes, as with NILs. Nevertheless, in both F_2_ and RIL populations, hundreds of lines are needed for the statistical power to detect additive effects, QTL intervals are still often large and contain hundreds or thousands of genes, and there is limited power to detect epistatic interactions (Mackay, 2014). In comparison, several advantages are offered by NILs, which each contain a small homozygous introgressed segment of an alternate genotype in an isogenic background, allowing for the direct comparison of genotypes differing in targeted genomic regions (i.e. broad scale allele swaps). This greatly simplifies the estimation of additive effect sizes, eliminates the averaging of additive effects across the genome (as in heterogenous mapping populations like RILs and F_2_), and aids in the detection of epistatic interactions between loci. Furthermore, NILs can be readily deployed to fine map causal loci in as few as 2 generations by backcrossing a NIL to the recurrent parent followed by selfing and genotyping. As such, NILs can be used for effect size quantification, fine mapping QTL, and/or breaking up QTLs linked in repulsion (Bentsink *et al*., 2003; Alonso-Blanco *et al*., 2003; Edwards *et al*., 2005; Chen & Yuan, 2024).

In addition to targeting single regions of the genome, NILs can be used to construct “introgression libraries” with most of the genome included in at least one introgressed segment of a NIL. This reduced number of lines representing a minimum tiling path across the genome results in maximum efficiency for experimental effect size estimation. One notable example of this approach used a set of NILs to identify many additive and one large-effect overdominant locus in tomato (Eshed & Zamir, 1994, 1995). NIL libraries or panels have been produced in *Arabidopsis thaliana* (Keurentjes *et al*., 2007; Törjék *et al*., 2008; Fletcher *et al*., 2013; Wijnen *et al*., 2024) and a variety of plant systems including cereal crops (Liu *et al*., 2006; Szalma *et al*., 2007; Ali *et al*., 2010), tomato (Monforte & Tanksley, 2000; Lippman *et al*., 2007; Chitwood *et al*., 2013), and a variety of other crop species (Ramsay *et al*., 1996; Eduardo *et al*., 2005; Shuang *et al*., 2023).

Near-isogenic lines can also be a powerful tool for detecting and quantifying non-additive genetic effects. The genetic simplicity and homozygosity of NILs can be leveraged with additional crossing (to produce heterozygous introgressions for example) and phenotyping to map dominance and epistatic effects (Melchinger *et al*., 2007; Reif *et al*., 2009; Guerrero *et al*., 2017; Khangura *et al*., 2019, 2020; Kaur *et al*., 2025; Rojas-Gutierrez *et al*., 2026). Intercrossing NILs can create double near-isogenic lines (Muir & Moyle, 2009; Yang *et al*., 2018) to test whether the phenotypic effects of the alleles in the double NIL is equal to the sum of the effects of both “single” NILs, as expected in the absence of epistasis.

Here we describe a new, densely genotyped, reciprocal panel of NILs developed from a cross between a pair of locally adapted ecotypes of *A. thaliana* (IT and SW), from central Italy and northern Sweden (Ågren & Schemske, 2012; Ågren *et al*., 2013; Oakley *et al*., 2023). To our knowledge, this is the only such NIL panel with direct evidence for local adaptation between the parental lines. This pair of accessions have additionally been used in extensive prior work mapping QTL affecting a wide range of ecologically important traits in RILs, including water use (Mojica *et al*., 2016), cold acclimation and freezing tolerance (Oakley *et al*., 2014; Sanderson *et al*., 2020; Lee *et al*., 2024), seed dormancy (Postma & Ågren, 2015), germination time (Postma & Ågren, 2018), flowering time (Dittmar *et al*., 2014; Ågren *et al*., 2017), photosynthetic traits (Oakley *et al*., 2018), and fitness components in the field (Ågren *et al*., 2013; Postma & Ågren, 2016; Ellis *et al*., 2021; Oakley *et al*., 2023). As such, these NILs represent an invaluable resource for the further dissection of the genetic basis of these and other traits. This densely genotyped reciprocal NIL panel from two natural (i.e., from intact native communities) populations provides a unique opportunity to fine map previously identified QTL, estimate additive effect sizes of genomic regions (for any trait where the ecotypes differ) without averaging over heterogeneous background effects, and with some simple additional crosses, to test for the contribution of dominance and epistasis to phenotypic differentiation.

Here, we fully describe genotype information from whole genome sequencing of the NIL panel and announce the availability of both the germplasm and associated sequence data of this unique genetic resource to the broader research community. To illustrate the utility of the NIL panel, we additionally investigate the genetic basis of time to first flowering (AKA flowering time) in a laboratory common garden experiment. Flowering time is well known to be an ecologically important trait (Lacey, 1988; Kalisz & Wardle, 1994; Olsson & Ågren, 2002), and its genetic basis is well studied (Simpson & Dean, 2002; Sung & Amasino, 2004; Amasino, 2010). Previous QTL mapped for flowering time in our mapping populations in both laboratory experiments (Grillo *et al*., 2013; Dittmar *et al*., 2014) and at the native field sites (Ågren *et al*., 2017) provide an opportunity to compare NIL results to QTL results from RIL and F_2_ mapping populations.

## Materials and Methods

### Study System

*Arabidopsis thaliana* is a small, predominantly selfing, annual plant with a large native range across much of Eurasia and with scattered occurrences across Africa (Koornneef *et al*., 1991; Beck *et al*., 2008; Durvasula *et al*., 2017; Lopez *et al*., 2025). Many natural populations, including our focal ecotypes, exhibit a winter annual life history. Our focal ecotypes include two lines collected from natural populations, one native to central Italy and one native to northern Sweden, near the northern range limit of *A. thaliana.* The Italian ecotype (IT, ABRC stock number: CS98761), collected from Castelnuovo di Porto, Italy (42°07′N, 12°29′E), germinates in October-November in the field and experiences cold but largely nonfreezing temperatures throughout the winter, before flowering in February-April (Ågren & Schemske, 2012; Ågren *et al*., 2017; Lee *et al*., 2024). The Swedish ecotype (SW, ABRC stock number: CS98762), collected from Rödåsen, Sweden (62°48′N, 18°12′E), germinates in August-September and experiences low autumn temperatures and winters with variable severity of freezing, and flowers in May-June (Ågren & Schemske, 2012; Ågren *et al*., 2017; Lee *et al*., 2024). Despite extensive investigation of the genetic basis of local adaptation using a RIL population derived from a cross between these ecotypes (ABRC stock number: CS98760), uncovering many QTL underlying long-term fitness (Oakley *et al*., 2023) and flowering time (Ågren *et al*., 2017) at the native sites, many of the causal variants remain unknown (but see Lee *et al*., 2024) because of the typically large QTL intervals identified.

### Construction of NIL Panel

Using this well-established RIL population as starting material, we created a genome-wide NIL panel in both ecotypic genetic backgrounds. We describe the whole-genome sequencing of these lines in detail in the next section but first give a brief overview of the process of constructing the panel. We selected ∼50 crosses between RILs and parental ecotypes for approximately complete coverage of alternate introgression segments in both backgrounds. See Figure S1 for a pedigree of descendants of one example of these ecotype-by-RIL combinations. For each combination, we performed hand pollinations with ecotypes as maternal parents and RILs as paternal parents.

We genotyped a single individual from each of two fruits from each of the combinations using multiple cleaved amplified polymorphic sequence (CAPS) markers (Table S1) to confirm heterozygosity where the RIL and parental genotypes differed. We then vernalized these individuals and backcrossed them to their recurrent ecotype, collecting seeds from 1-6 fruits for each BC_1_ combination. We grew approximately 16 individuals from each fruit and genotyped each individual and the parental ecotypes in a highly multiplexed fashion using a Type-II restriction site-associated DNA approach (2b-RAD; Wang *et al*., 2012), and called genotypes with Stacks (Catchen *et al*., 2013). This yielded approximately 200-400 SNP genotypes per line. We collected autogamously selfed seeds (BC_1_S_1_) from these individuals. We prioritized specific regions of the genome underlying QTL for lifetime fitness at both field sites over 3 years identified by Ågren et al. (2013) by multiple rounds of either backcrossing or selfing and genotyping using CAPS or TaqMan markers (Tables S1 & S2) followed by another round of 2b-RAD genotyping. This process resulted in hundreds of lines with varying degrees of heterozygosity and homozygous introgression segment sizes. For the subset of the overall panel presented here, we selected a total of 54 NILs (23 in the IT background and 31 in the SW background) for advancement and whole genome sequencing. These lines had homozygous introgression segments spanning as much of the genome as possible (covering all previously reported fitness QTL) in otherwise isogenic ecotypic backgrounds.

### NIL Genotyping by WGS

#### Identification of Diagnostic SNPs in Parental Ecotypes

To establish a set of high-confidence SNP markers for this pair of ecotypes, we harvested leaf tissue from each parental ecotype and extracted DNA using the Qiagen DNeasy Plant Mini Kit (Valencia CA, USA). WGS libraries were prepared at the University of Colorado Boulder sequencing core and sequenced (150bp paired-end reads) over 2 lanes of an Illumina HiSeq sequencer at the University of Colorado Anschutz Medical Campus. We removed sequencing adaptors from the raw reads using Trimmomatic (v0.39; Bolger *et al*., 2014), confirmed adaptors had been removed with FastQC (v0.12.1; https://www.bioinformatics.babraham.ac.uk/projects/fastqc/), and merged files originating from independent sequencing lanes with the Unix ‘cat’ command. We then aligned reads to the Araport11 *A. thaliana* reference genome (https://phytozome-next.jgi.doe.gov/) using BWA-MEM (v0.7.17; Li, 2013; Li and Durbin, 2009) using default parameters. Using SAMtools ‘view’ (v1.7; Li *et al*., 2009), we removed reads with alignment qualities <Q29, and sorted the subsequent BAM files with SAMtools ‘sort’. We used Picard (v2.26.10; broadinstitute.github.io/picard) to identify and fix incorrectly mated reads, added read groups, and removed PCR duplicates using ‘FixMateInformation’, ‘AddOrReplaceReadGroups’, and ‘MarkDuplicates’ respectively. Finally, we used SAMtools ‘view’ flag 524 to ensure reads were properly paired.

Using the resultant filtered BAM files, we used BCFtools ‘mpileup’ and ‘call -m’ (v1.17; Danecek *et al*., 2021) to call SNPs. We used GATK ‘VariantFiltration’ (v3.8.1; McKenna *et al*., 2010) to filter heterozygous calls, and calls with depth less than 6x, or greater than 2 standard deviations above the genome wide mean (calculated for each line with Qualimap2 v2.2.1; Okonechnikov *et al*., 2016, IT: 111x, SW: 158x). Additionally, we identified a subset of SNPs with a minimum depth of 12x to be used for high-confidence genotyping near putative introgression break points. We used BCFtools ‘merge’ (v1.17; Danecek *et al*., 2021) to create a multi-sample VCF containing both parental ecotypes and used GATK ‘VariantFiltration’ (v3.8.1; McKenna *et al*., 2010) to filter sites with a mapping quality of less than 40, followed by ‘SelectVariants’ to retain only biallelic sites, remove filtered sites, and remove calls from filtered genotypes. We then used VCFtools (v0.1.16; Danecek *et al*., 2011) to retain sites polymorphic between the ecotypes that were also identified as high quality variant sites by the 1001 Genomes Consortium (Alonso-Blanco *et al*., 2016). This produced datasets of 361,867 SNPs that distinguish the haplotypes of the parental genomes (∼1 SNP per 330bp) in the 6x dataset, to be used for genotyping the NILs across the genome, and 162,076 SNPs (∼1 SNP per 735bp) in the 12x dataset, to confirm genotypes near introgression breakpoints.

#### NIL DNA Extraction and Whole Genome Sequencing

We collected leaf tissue from a single plant of each NIL genotype and stored it at −80°C before freeze drying the tissue and grinding it with stainless steel ball bearings in a tissue homogenizer (Tallboys, Troemner LLC, Thorofare, NJ). We extracted DNA with a modified SDS extraction method (Edwards *et al*., 1991) in a 96-well format and resuspended it at 5-20 ng/uL in de-ionized (DI) water. We submitted all samples to the Novogene Corporation Inc. UC Davis Sequencing Center for preparation of DNA-seq libraries and sequencing (150-bp paired-end reads) on an Illumina NovaSeq X Plus sequencer, generating approximately 3Gb of data per sample.

#### Sequence Processing and Genotyping

We processed reads from each of the 54 NILs using the same bioinformatic tools and parameters described above. We called SNPs and filtered them as above but retained heterozygous calls to identify residual heterozygosity in the NILs. We also filtered and removed genotypes with depths less than 3x, or greater than 2 standard deviations above the genome wide mean for that line (Table S3, 136x – 368x). We then created two multi-sample VCF files, each containing both parental ecotypes, and all NILs from one of the two genetic backgrounds. Finally, we filtered these VCFs, retaining only sites contained in the list of diagnostic sites identified above (6x dataset).

#### Determination of Introgression Location

We used our identified diagnostic sites (6x dataset) to determine the genotype of each NIL across the genome. We visualized these data using three complementary methods. For each NIL we calculated genotype proportions in 25kb (1) and 100 SNP (2) non-overlapping windows across the genome. We then identified haplotypes in each NIL (3), requiring at least three consecutive SNPs of the same genotype (IT, SW, or Heterozygous) covering at least 5kb for positive identification. Consistent agreement among these methods allowed for identification of genotypes across the genome in each NIL. We checked each block of introgression not identified by previous genotyping approaches by visual inspection of reads in the Integrative Genomics Viewer (IGV; Robinson *et al*., 2011). We additionally identified and removed rare false genotype calls caused by poor mapping and low read coverage (frequently in and around centromeres), when they appeared across multiple unrelated NILs. To ensure accuracy, we used genotypes at our identified high-quality sites (12x dataset) to confirm genotypes near each identified breakpoint.

### Plant Growth and Phenotyping

As a small-scale example of the utility of this NIL panel we conducted a common garden experiment with all 54 NILs and the parental ecotypes in common growth chamber conditions to estimate the effects of individual genomic regions on flowering time. For each genotype, we surface sterilized ∼15 seeds by submerging them in a 20% v/v bleach and 0.04% v/v Tween 20 solution for 10 min. After repeated rinsing in sterile DI water, we suspended seeds in 0.1% w/v Phytoblend agar (Caisson Laboratories, Smithfield, UT, USA). Seeds were sown on media consisting of autoclaved Gamborg’s B-5 basal salts (with sucrose) and 0.64% w/v Phytoblend agar poured into sterilized petri dishes. We cold stratified plated seeds in the dark at 4°C for 5 days to break dormancy and synchronize germination before moving plates to a growth chamber at 22°C, 16-h day (16L:8D), for 15 days. We then transplanted 8 seedlings of each genotype (448 total plants) into 72-cell flat inserts (12.25 cm^2^/cell) containing moist Berger BM2 germination and propagation mix, and grew them on a light rack at room temperature (12L:12D) for 8 days before being vernalized in a refrigerator fitted with lights at 4°C (10L:14D) for 8 weeks. During vernalization we bottom watered flats with DI water approximately once every two weeks.

Following vernalization, we transplanted each plant into a 36 cm^2^ pot containing Berger BM2 germination and propagation mix in a fully randomized design and moved them to a growth chamber at 22°C (14L:10D). We bottom watered plants twice a week with ½ strength Hoagland’s solution and recorded day of bolting and day of first flowering (n=8 replicate individuals for each parental ecotype and NIL) relative to “Day 0”, the day plants were removed from vernalization. Day of bolting was defined as the day on which the inflorescence first visibly extended beyond the rosette, and day of first flowering was defined as the day on which the corolla was first visible in an opening bud. As plants matured, we added bases and Aracons (Arasystem, Ghent, Belgium) to each plant to collect all seeds produced. When plants began to senesce and were no longer producing new flowers, we allowed them to dry-down from saturation on a light rack at room temperature until all fruits were dehiscent.

Total seed production per plant was recorded to provide an estimate of seed production that can be expected by “bulking” up seed under standard laboratory conditions, and also to ensure the absence of genetic incompatibilities grossly affecting overall vigor in some NILs. We collected all seeds produced by each plant (8 of each parental ecotype, and 3 of each NIL genotype) and recorded total seed mass per plant. Because total seed mass per plant can be influenced by seed size (Gnan *et al*., 2014), seed number, or both, we also used Fiji (Schindelin *et al*., 2012) to estimate seed size as the average area covered by each seed in an image of a pool of >30 seeds taken with a flatbed scanner (Herridge *et al*., 2011; Van Daele *et al*., 2012). We found no correlation between individual total seed mass and individual mean estimated seed size (Kendall’s τ = 0.05, p = 0.577) and therefore use total seed mass as a proxy for total seed number per plant.

### Testing for Phenotypic Effects of Introgressed Regions

To test for the effect of NIL introgression segment on flowering time we fit non-parametric linear models (on ranked values because residuals indicated departures from normality and homogeneity of variances) with the lm command in the ‘lme4’ package (v1.1-34, Bates *et al*., 2015) in R (v4.3.1). Models for both days to flower and days to bolt tested the effects of genotype (IT and SW ecotypes plus 23 IT background and 31 SW background NILs), as well as distance from the center of the flat during vernalization to account for spatial variation in light intensity. Both terms were treated as fixed effects. To test hypotheses about the influence of specific genomic regions on flowering time we used the emmeans command in the ‘emmeans’ package (v1.8.8, Lenth, 2025) to calculate genotype least squares means (LSMs) from each model (with the same procedure for seed mass per plant and average seed size). Least squares mean comparisons of each NIL to its background ecotype were performed with post hoc Dunnett’s tests (contrast command in the ‘emmeans’ package). For these comparisons, we adjusted p-values for the number of tests performed (23 in the IT background, 31 in the SW background).

## Results

### WGS Based Genotyping

Overall, WGS derived genotypes and introgression breakpoints agreed with reduced representation and PCR based marker genotyping but provided much greater confidence and higher resolution due to increased marker density. Whole genome sequencing resulted in 361,867 SNP markers (6x dataset; ∼1 SNP per 330bp), improving the resolution of recombination breakpoints nearly 150-fold, and practically eliminating the likelihood of unidentified small introgression blocks in the NILs. In total, introgression segments in this panel cover 68.1% and 79.1% of the *A. thaliana* reference sequence in the IT and SW backgrounds respectively (Fig. 1).

**Figure 1.**
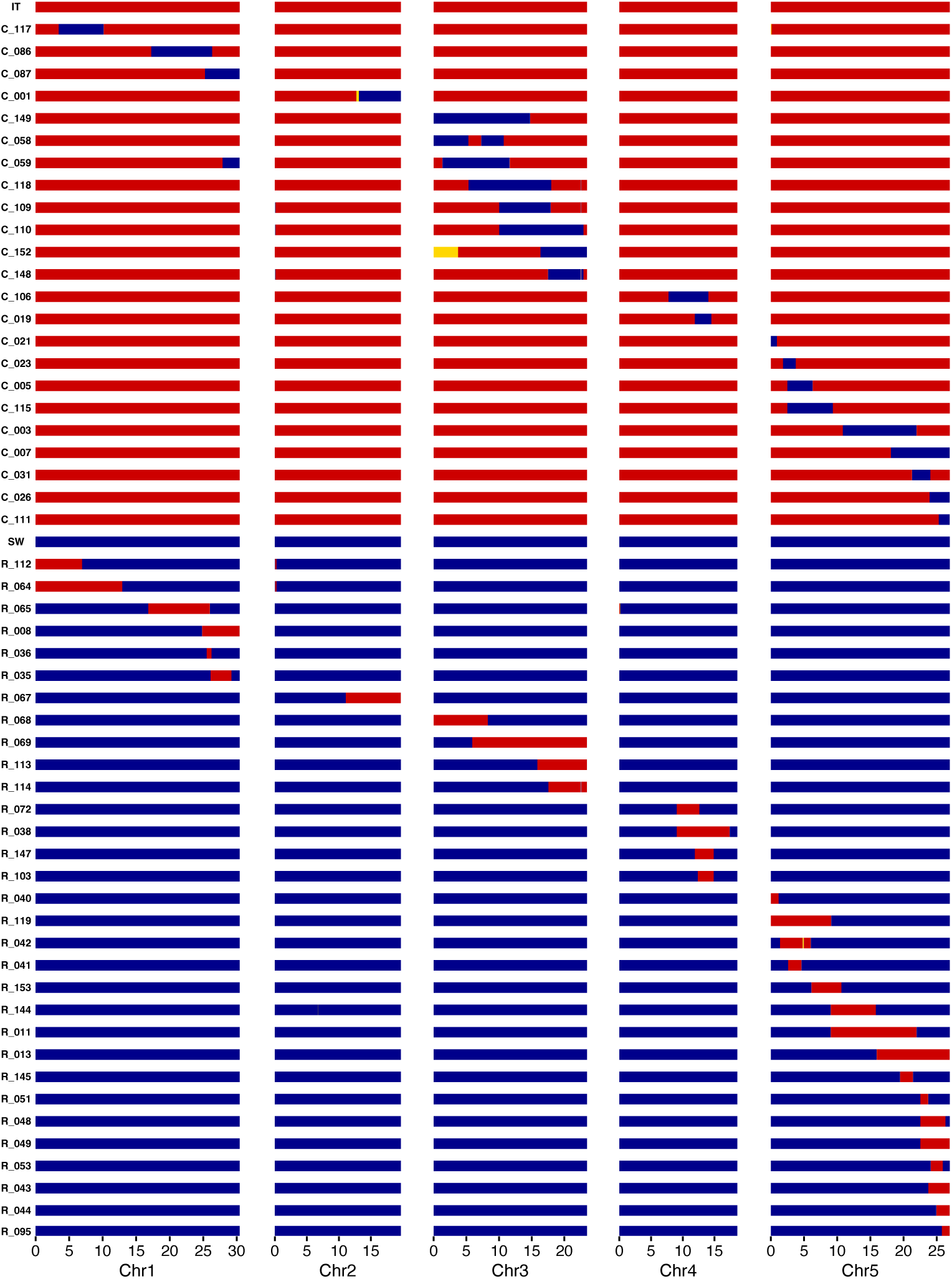
Genotypes of the full NIL panel and parental ecotypes. Horizontal bars represent the 5 chromosomes of the *A. thaliana* genome for each of the 54 NILs and the parental ecotypes. NILs in the IT background (ID starts with C) are predominantly red and NILs in the SW background (ID starts with R) are predominantly blue. NIL IDs are shown along the y-axis and the genomic location (Mb) of IT (red) and SW (blue) genotypes is shown on the x-axis. Regions of retained heterozygosity (gold) and crossover regions between SNP markers (grey) are also shown. NILs are ordered by the starting position of their main introgressed region. Finer scale genotype information surrounding introgression breakpoints is shown in Figs. 2 and S2.

With WGS based genotyping we identified highly accurate positional information for each introgressed segment in all NILs (Figs. 1, 2, & S2; Table S3). The panel contains an average SW introgression size of 6.94 Mb (0.93 – 14.76 Mb) in the IT background, containing an average of 1,720 genes (range = 281 – 3,439), and an average IT introgression size of 5.66 Mb (0.70 – 17.53 Mb) in the SW background, containing an average of 1,560 genes (range = 176 – 3,665; Table S3). In addition to higher resolution of known introgression segments, WGS data for the whole NIL panel identified 12 small introgression segments (range = 0.006 - 0.18 Mb, mean = 0.06 Mb) and one larger (2.53 Mb) segment not detected with previous genotyping methods. The WGS data also revealed 5 small (0.01 - 0.37 Mb, mean = 0.13 Mb) and one larger (3.74 Mb) regions of retained heterozygosity not visible by other methods. Newly identified homozygous segments larger than 150 kb, and the largest heterozygous segment (i.e. those regions large enough to contain multiple usable PCR based markers) were directly confirmed by genotyping with 2 or more additional CAPS markers in at least two individuals (Table S1). Newly identified introgression blocks and regions of retained heterozygosity were overwhelmingly located in areas of the genome with relatively few genotyped 2b-RAD/CAPS/TaqMan markers – i.e. regions of the genome with the lowest resolution in our previous genotyping scheme.

**Figure 2.**
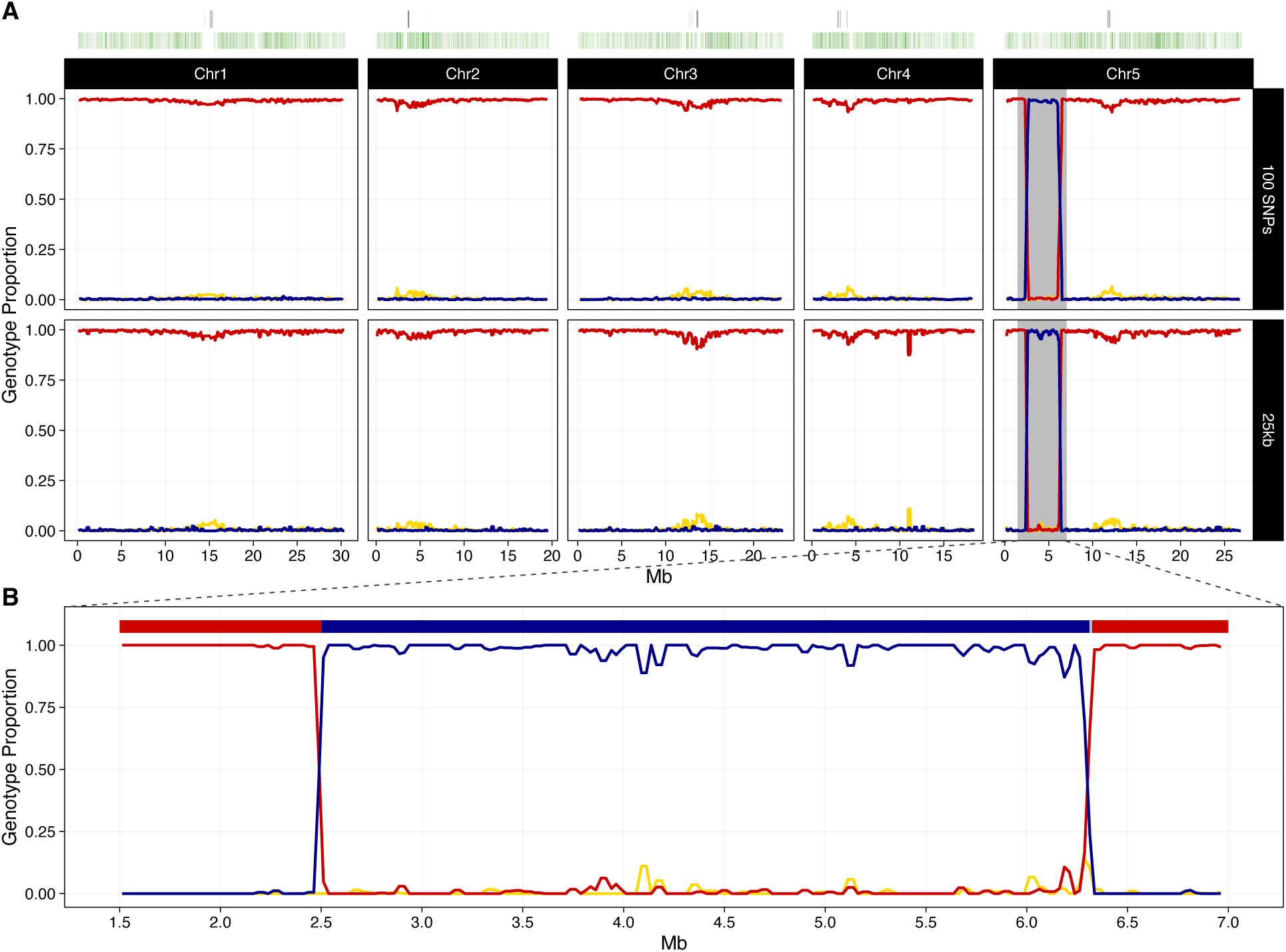
Detailed genotyping of a representative example NIL (C_005). (A) Rolling average (period = 10) of genotype proportion (IT: red, SW: blue, heterozygous: gold) in non-overlapping 100 SNP windows and non-overlapping 25kb windows across the 5 chromosomes of the *A. thaliana* genome. Heatmaps above show the density of centromeric repeats (grey) and density of diagnostic SNP markers (green) across the genome. (B) Zoomed in view of the introgressed region of interest at the beginning of chromosome 5, rolling average (period = 2) of genotype proportion (IT: red, SW: blue, heterozygous: gold) in non-overlapping 25kb windows, with a summary haplotype map above. Crossover regions between SNP markers (grey) are also shown. Detailed genotyping information for the remaining NILs of the panel is shown in Fig. S2.

### Genomic Regions Contributing to Flowering Time

On average, the IT ecotype flowered 18 days earlier than the SW ecotype (18 vs. 36 days after the end of vernalization; Fig. 3; Table S4). As a whole, the IT background NILs flowered more than 15 days earlier on average than the SW background NILs (Fig. 3), as expected based on ecotypic differences in flowering time. Within each background, we observed extensive variation in flowering time among the NILs, including some transgressive segregation (Figs. 3 & S3; Table S4). In the IT background, the only statistically significant effects were delays in flowering in 2 NILs (Figs. 4A & S3A; Table S4). These two lines flowered ∼6 (C_005) and 9 days (C_115) later than IT on average. Both lines carry partially overlapping introgression segments near the top of chromosome 5, that contain the well-studied floral regulator *FLOWERING LOCUS C* (*FLC*; Fig. 1; Table S3). There were additionally several NILs that did not meet the significance threshold, but which produced large average effects on flowering time (more than 4 days). These three IT background NILs all carry introgression segments on chromosome 3 and flowered between ∼5 and 6 days faster than the IT ecotype (Fig. S3A; Table S4). The direction of these effects was unexpected because the SW genotype delayed flowering at every previously detected flowering time QTL in this population (Grillo *et al*., 2013; Dittmar *et al*., 2014; Ågren *et al*., 2017).

**Figure 3.**
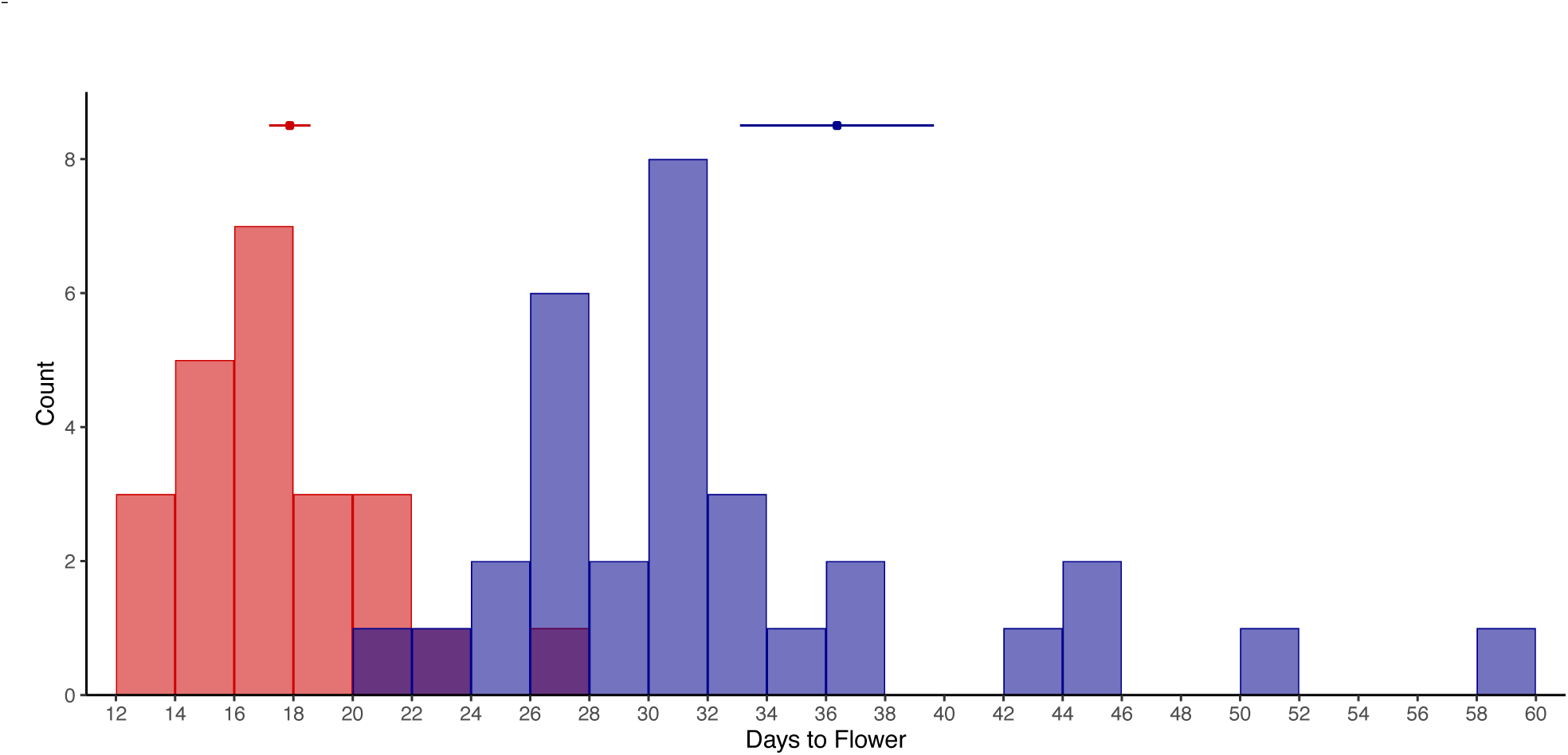
Distribution of NIL mean days to first flowering. IT background (red), SW background (blue). Means and +/- SE for the parental ecotypes, IT (red) and SW (blue) are shown above.

**Figure 4.**
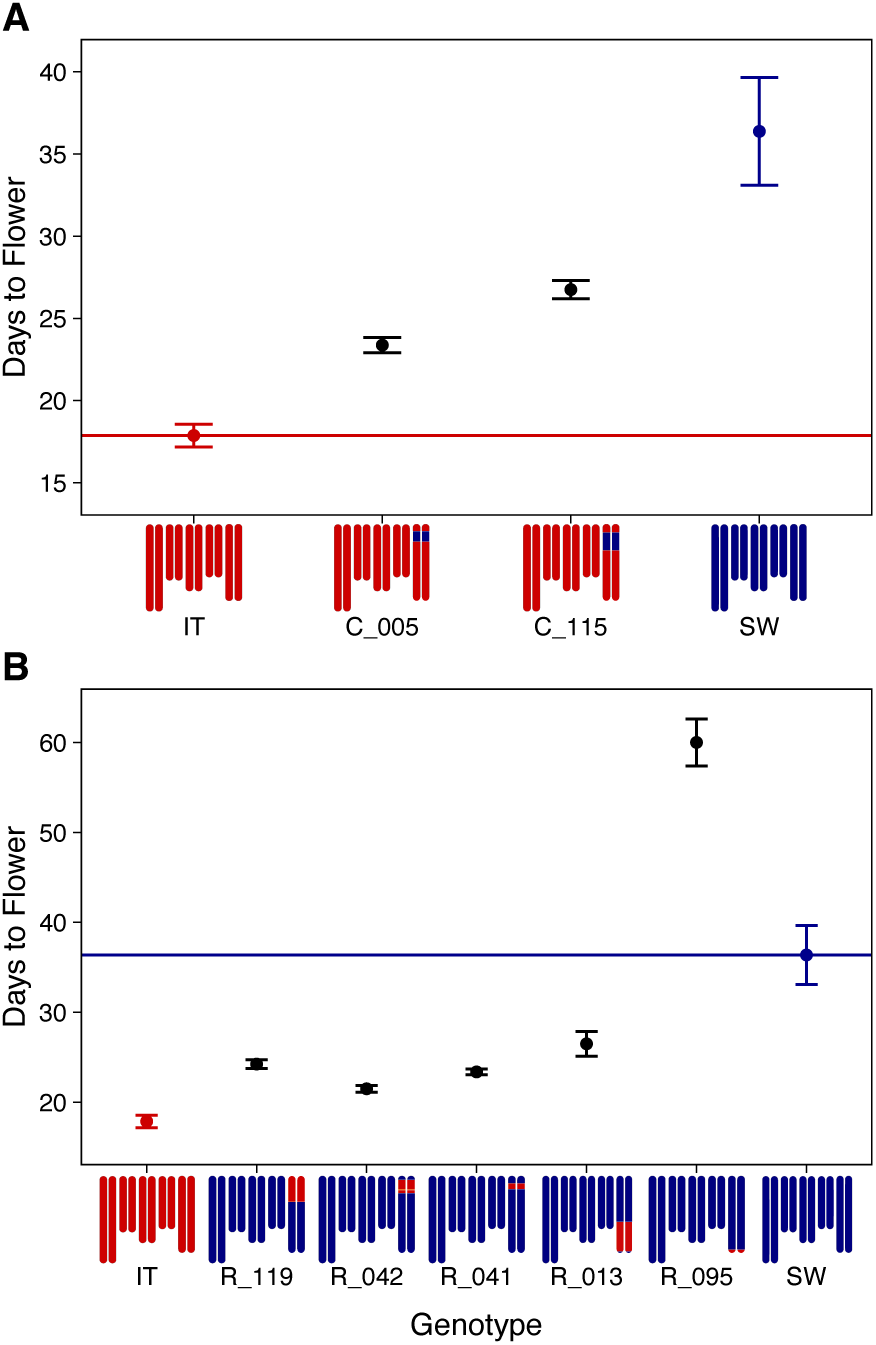
Genotype mean days to first flowering of NILs with significant differences from their recurrent ecotype. p<0.05, corrected for multiple testing by a post hoc Dunnett’s test. (A) Mean days to first flowering +/- SE of ecotypes, IT (red), SW (blue), and NILs in the IT background (black). (B) Ecotypes and NILs in the SW background. All NILs are shown in Fig. S3. Note that panels (A) and (B) have different y-axis scales.

In the SW background NILs, flowering time was significantly accelerated in 4 NILs and significantly delayed in 1 NIL when compared to the SW ecotype (Figs. 4 & S3B; Table S4). Three of the NILs with significantly accelerated flowering (R_041, R_042, and R_119) flowered between ∼12 and 15 days faster than the SW ecotype (Fig. 1; Table S3), and all share partially overlapping introgression segments near the top of chromosome 5 containing *FLC*. The remaining SW background NIL with significantly accelerated flowering (R_013) flowered ∼10 days earlier than the SW ecotype (Fig. 4B) and contains an introgression segment at the bottom of chromosome 5. Surprisingly, 1 SW background NIL containing a small (∼1Mb) introgression encompassed within the introgression in R_013 at the bottom of chromosome 5 had severely delayed flowering (R_095, ∼24 days slower than SW on average; Figs. 4B & S4). None of the other SW NILs with introgression segments in this region had delayed flowering (Fig. S4). Many SW background NILs also had large effects on flowering time which did not reach our significance threshold. Fifteen SW NILs flowered between ∼4.5 and 9 days faster than the SW ecotype (Fig. S3; Table S4) with introgression segments across the five chromosomes, and 4 SW NILs unexpectedly flowered between ∼6.5 and 14 days slower than the SW ecotype (Fig. S3; Table S4) with introgression segments on chromosomes 1, 3, and 4.

Results for bolting time generally follow similar patterns to flowering time (Figs. S5 & S6), and bolting time and flowering time were strongly correlated (Kendall’s τ = 0.89, p < 0.001). However, there was 1 NIL in each background with significant effects on bolting time but not flowering time (Figs. S5 & S6; Table S4). C_007 bolted ∼2 days later than IT and R_011 bolted ∼4 days earlier than SW. Both contain an introgression segment overlapping the one in R_013 associated with flowering time. Additionally, the interval between bolting and flowering was significantly affected in one NIL (R_036; Table S4) which matured and opened its buds an average of ∼6 days faster than SW.

### Total Seed Mass

On average the parental ecotypes had similar total seed mass per plant (IT: 0.23g, SW: 0.20g). As with flowering time, there was considerable variation among NILs for mean seed mass per plant (Figs. S7 & S8; Table S4). The only 2 NILs with significant effects (both reduced compared to the background ecotype) on total seed mass per plant (C_115 and R_095) were those with the largest observed delays in flowering time in each background (Figs. S5 & S6; Table S4). However, seed mass and flowering time varied independently such that other NILs with significant effects on flowering time had no consistent effects on total seed mass (Figs. S7 & S8; Table S4), suggesting that the reduced seed production of these 2 lines in this environment is unlikely to be solely the result of differences in flowering time.

## Discussion

Here we announce the availability of germplasm and high-resolution genotype information for a reciprocal NIL panel derived from a cross between natural populations of *Arabidopsis thaliana*. To our knowledge, this is the only such panel with direct evidence for local adaptation between the parental lines. Extensive fitness, trait, QTL, and climate data is available from long term field experiments on this pair of ecotypes (Ågren & Schemske, 2012; Postma & Ågren, 2016, 2018; Ågren *et al*., 2017; Ellis *et al*., 2021; Oakley *et al*., 2023; Lee *et al*., 2024). Ecotypic differences in a broad suite of traits have also been documented and mapped in controlled environment experiments (Dittmar *et al*., 2014; Oakley *et al*., 2014, 2018; Mojica *et al*., 2016; Sanderson *et al*., 2020). There is also documented evidence for a role of both dominance and epistatic effects affecting fitness in these populations (Ågren *et al*., 2013; Oakley *et al*., 2015, 2019, 2023; Ellis *et al*., 2021). Despite extensive research, the causal variants underlying many QTL for these traits remain undiscovered. This new NIL panel will enable fine mapping of candidate genes as well as estimation of additive effects contributing to phenotypic differences between the ecotypes. In addition, simple crosses of these NILs to their recurrent parents permit direct testing of dominance and epistatic effects (Rojas-Gutierrez *et al*., 2026). When combined with environmental conditions relevant to the native sites, these NILs can also be used to investigate the genetic basis of adaptive plasticity and local adaptation (Sanderson *et al*., 2020; Lee *et al*., 2024). While other panels of NILs are available in *A. thaliana* (Keurentjes *et al*., 2007; Törjék *et al*., 2008; Fletcher *et al*., 2013; Wijnen *et al*., 2024), the foundation of ecological and evolutionary context in this pair of ecotypes is unique.

An additional value of this panel is the high density of SNP markers from whole genome sequencing. This resolved introgression breakpoints to an average of 165 bp windows. This high-resolution genotyping also uncovered small introgression blocks and regions of retained heterozygosity (Fig. S2; Table S3). The better performance of whole genome sequencing relative to RAD seq and PCR markers suggest that heterozygosity and small introgressions are likely present, but undetected, in many NIL and RIL panels. The genomic coverage of introgression segments in our panel is nearly 70% of the genome in the IT background, and nearly 80% in the SW background (Fig. 1; Table S3), comparable to many NIL panels (for example: Eduardo *et al*., 2005; Schmalenbach *et al*., 2011; Shuang *et al*., 2023) but less than some panels in *A. thaliana* and tomato (for example: Eshed & Zamir, 1995; Törjék *et al*., 2008; Fletcher *et al*., 2013). Even though the present NIL panel has incomplete genomic coverage, it still represents the best available NIL panel for populations that have been demonstrated to be locally adapted. Furthermore, the introgression regions in the current panel span all QTL for lifetime fitness detected at the two parental sites over 3 years (Ågren *et al*., 2013), suggesting that most of the genomic regions governing ecologically important differences between these ecotypes is represented in this panel. That said, we have additional NILs under construction to fill the remaining genomic gaps (Fig. 1; top of chromosomes 2 and 4 for example) and will make the germplasm available at ABRC, along with their genotype data, as they are completed.

### Proof of concept: Effects of genomic regions on flowering time

The IT ecotype flowered earlier than the SW ecotype in our controlled environment (Fig. 3) as expected from previous results in growth chambers and at both native field sites (Grillo *et al*., 2013; Dittmar *et al*., 2014; Ågren *et al*., 2017). The magnitude of the difference (∼18 days) was less than that observed in a previous growth chamber experiment (∼27 days; Grillo *et al*., 2013), and in field experiments at the Italian site (∼42 days averaged over 3 years) but much greater than in field experiments at the Swedish site (∼5 days averaged over 3 years; Ågren *et al*., 2017).

All of the 7 NILs with significant effects on flowering time contained introgression segments on either end of chromosome 5 (Fig. 4). This is in agreement with the detection of flowering time QTL at both ends of chromosome 5 in IT x SW mapping populations in all five previously studied environments (Grillo *et al*., 2013; Dittmar *et al*., 2014; Ågren *et al*., 2017). Five NILs with altered flowering time had introgression segments at the top of chromosome 5 spanning the well-studied flowering time regulator *FLC* (Fig. 4), known to modulate flowering behavior mainly through the vernalization pathway (Yan *et al*., 2010; Whittaker & Dean, 2017). The SW allele in this region delayed flowering in the IT background by as much as 9 days, while introgression of the IT allele accelerated flowering in the SW background by up to 15 days (Fig. 4; Table S4). Similarly, flowering time QTL spanning *FLC* with large additive effects were previously found both in an F_2_ population grown in a growth chamber environment similar to the present study (Grillo *et al*., 2013), and in RILs grown in growth chambers simulating temperature and photoperiod at the native sites (Dittmar *et al*., 2014). In field experiments using RILs however, QTL overlapping *FLC* have been shown to consistently influence flowering time at the Italian field site, but not the Swedish site (Ågren *et al*., 2017). It may be that the extended period of temperatures colder than 4°C of the Swedish field conditions minimize differences between IT and SW genotypes at the *FLC* locus by “saturating” vernalization requirements, while the constant 4°C vernalization period in the present study or the warmer vernalization conditions at the Italian field site reveals differences at *FLC* important for initiation of flowering. This might also contribute to minimal ecotypic differences in flowering time in Sweden compared to Italy (Ågren *et al*., 2017).

The coding sequences of *FLC* in IT and SW differ by only a single synonymous SNP (Grillo *et al*., 2013) suggesting that divergent regulatory elements controlling *FLC* are likely responsible for its effect on flowering time and fitness. This is consistent with studies demonstrating the importance of cis-regulatory/promoter haplotype variation in the regulation of *FLC* expression, vernalization response, and seasonal flowering time variation across the range of *A. thaliana* (Shindo *et al*., 2006; Li *et al*., 2014a; Hepworth *et al*., 2020; Zhu *et al*., 2023). The gene *FRIGIDA* (*FRI*) has received considerable attention as a major determinant of flowering time upstream of *FLC* (Simpson & Dean, 2002; Le Corre *et al*., 2002). However, both SW and IT ecotypes have functional *FRI* alleles (Grillo *et al*., 2013) explaining the lack of flowering time QTL detected near *FRI* (Grillo *et al*., 2013; Dittmar *et al*., 2014; Ågren *et al*., 2017). The lack of introgression segments at the top of chromosome 4 in our NIL panel is therefore unlikely to influence our flowering time results.

The two other NILs (both in the SW background) with significant effects on flowering time each had introgression segments at the bottom of chromosome 5, but with effects in opposite directions (Fig. 4; Table S4). The large IT introgression region contained in R_013 significantly accelerated flowering time (∼10 days faster) as expected based on flowering time QTL previously mapped to the bottom of chromosome 5 (Grillo *et al*., 2013; Dittmar *et al*., 2014; Ågren *et al*., 2017). However, a much smaller nested subset of that introgression segment in R_095 significantly delayed flowering (nearly 24 days slower) relative to the SW ecotype (Figs. 4 & S4). The strongly transgressive flowering time of this NIL is in the opposite direction expected based on flowering time QTL detected previously in this system in three different studies and five different environments, where the IT genotype always accelerated flowering (Grillo *et al*., 2013; Dittmar *et al*., 2014; Ågren *et al*., 2017). This novel result is further noteworthy because such a strong delay in flowering of the IT genotype at this locus in Italy would be strongly selected against (Ågren *et al*., 2017). No other NILs with introgression segments in this region had significant effects of flowering time, and the NIL with the most similar introgression segment in the IT background had no effect on flowering time (Fig. S4).

Taken together our results lead us to hypothesize that there is an epistatic interaction between the IT genotype at the very bottom of chromosome 5 (near FlrT 5:5 detected at the Italian Site, Ågren *et al*., 2017; Fig. S4), and the SW genetic background. Such an interaction would suppress the strong delay in flowering in other SW background NILs with introgressions spanning these regions (R_043, R044, and R_049) and could explain the lack of an effect on flowering time in C_111 (Fig. S4). Though it is formally possible that there is a novel locus that strongly accelerates flowering time closely linked to the region contained in R_095 between ∼25-26 Mb (Fig. S4), we have not previously observed flowering time QTL in that region and we did not observe a significant or even strong decrease in flowering time for NILs R_053 and R_048 (Fig. S4). The lack of prior evidence for epistatic interactions for flowering time in this region of chromosome 5 (Grillo *et al*., 2013; Dittmar *et al*., 2014; Ågren *et al*., 2017) could indicate environmental dependence of the epistatic effects, but also may result from limited power to detect epistatic interactions between tightly linked loci and higher order epistasis using QTL mapping approaches in F_2_ and RIL populations. In this case, the hypothesized interaction would only be discoverable using a NIL panel such as ours. Using this NIL panel as a starting point, simple backcrosses could be done using NILs in Fig. S4 to test if the effects observed here are due to strong antagonistic effects linked in repulsion. If not, further crosses to produce combinatorial NILs could be used to directly test for epistatic effects. Either way, these NILs are useful for investigating the novel genetic effect discovered here and have utility in understanding the genetic basis of complex traits, as well as questions about the genetic basis of transgressive segregation more generally (Rieseberg *et al*., 1999, 2003; Dittrich-Reed & Fitzpatrick, 2013; Pabuayon *et al*., 2021).

One important caveat to our results on flowering time is that we have limited statistical power because of our small-scale experiment. Our purpose here was not to provide definitive answers to the genetic basis of flowering time, but to simply illustrate the utility of the panel for addressing these questions. That we were able to detect these effects even in the present experiment with limited replication speaks to the power of the NIL population. Many of the non-significant but large effects (4+ days) detected here would likely be statistically significant even in a modestly larger experiment targeting specific regions of the genome. Additionally, subsets of NILs have already been used to estimate effects of genomic regions for freezing tolerance (Sanderson *et al*., 2020), local adaptation (Lee *et al*., 2024), and heterosis (Rojas-Gutierrez *et al*., 2026), demonstrating their utility for examining the genetic basis of a variety of different traits.

### Summary

We announce the availability of a new genetic resource: germplasm and associated sequence data for a unique reciprocal NIL panel. All lines have been genotyped with high density SNP markers, ensuring accurate genotyping information and all but eliminating the probability of unidentified introgression blocks. Such panels are rare and thus represent a valuable resource for investigating the genetic basis of complex traits. Illustrating the utility of the panel to estimate effects of genomic regions affecting flowering time in a small-scale experiment, we both confirmed results for previously identified QTL containing *FLC* and uncovered a novel locus (or epistatic interaction) that has not been reported previously. In addition to indicating the complex genetic architecture of this trait in this environment, the NIL panel provides a mechanism for testing for epistatic effects in the short-term future using NILs with multiple introgression segments produced by a single round of additional crosses. More generally, this panel is derived from a pair of well-studied, locally adapted natural populations of *A. thaliana*, with a wealth of prior environmental, fitness, and trait data, and thus is a valuable resource for the study of the genetic basis of any trait for which the ecotypes differ. Replicating experiments with these lines across environments can also provide insight into the genetic basis of genotype-by-environment interactions, and paired with environmental conditions relevant to the native field sites, can provide unique insight into the genetic basis of adaptive plasticity and local adaptation (Sanderson *et al*., 2020; Lee *et al*., 2024).

## Acknowledgements

We thank M. Johnson, J. Miller, A. Hostrawser, S. Andrewlavage, and S. Shen, for assistance with plant care, phenotyping, and seed collection. S. Ahmed, P. Chidambaram, R. Deater, J. French, C. Gallick, J. Kraft, K. Palacio-Lopez, S. Park, A. Ralston, I. Turner, P. Caldwell, H. Chou, and C. Leclercq assisted with NIL development. We also thank the Purdue University Plant Growth Facility staff including M. Woodard and B. Kim. For valuable comments on earlier drafts of this manuscript, we thank S. Mills, E. Lancaster, and J. Miller. Special thanks to D.W. Schemske for the inspiration and early support to make the panel. This work was supported by National Science Foundation grants DEB-1743273 to C.G.O. and J.K.M., IOS-2246545 to C.G.O. and B.P.D., and DEB-2325338 to C.G.O.

## Data Accessibility Statement

Raw whole genome sequence data is archived on the NCBI Sequence Read Archive (BioProject: PRJNA1372624). Phenotypic data and VCF files containing NIL SNP calls are archived at the Dryad Digital Repository (https://doi.org/10.5061/dryad.hhmgqnkw5). Germplasm is deposited at the ABRC (Stock Number: CS99678).

## Author Contributions

Research conceived and designed by C.G.O., J.K.M., J. Å., and S.J.M. NILs developed by C.G.O., J.D.R.G., G.L., B.J.S., and M.I.J. Sequencing of ecotypes by P.W. and J.K.M. NIL sequencing, analysis, and genotyping by S.J.M. Phenotypic data collected and analyzed by S.J.M. Manuscript written by S.J.M., J.D.R.G., and C.G.O. Supervision by J.K.M., J.Å., B.P.D., and C.G.O. All authors contributed to revising the manuscript.

## Supplemental Materials

**Table S1. PCR based markers used for NIL genotyping.** Cleaved amplified polymorphic sequence (CAPS) markers used for initial genotyping during NIL construction. Additional CAPS markers used to confirm previously unidentified introgressed regions and regions of retained heterozygosity discovered with WGS based genotyping are also shown.

Table S2. PCR **based markers used for NIL genotyping.** Taqman markers used for initial genotyping during NIL construction. FAM: 5-(and 6-) carboxy-fluorescein, VIC: 2′-chloro-7′phenyl-1,4-dichloro-6-carboxy-fluorescein.

**Table S3. Summary of WGS and NIL genotypes.** Genome wide mean and standard deviation of sequencing depth for each NIL genotype separated by background genotype. Chromosome (Chr) containing an introgressed region, the start and end of that region, the total size of the region (Mb), and the number of genes contained within, is shown for each NIL. Regions of the background genotype are only included for a NIL if they are entirely contained within an introgressed region of the alternative background (i.e. representative of small double crossover regions).

**Table S4. Full phenotypic summary.** Mean days to bolt, days to first flowering, difference between days to bolt and days to first flowering, total seed mass per plant, and seed size and standard error for each ecotype and NIL genotype. The proportional change (effect) of each introgressed genotype relative to its recurrent ecotype and P-value (corrected for multiple testing with a post hoc Dunnett’s test, and uncorrected) from comparing each NIL to its recurrent parent are also included. NIL genotypes significantly different than their recurrent ecotype by Dunnett’s test are shown in bold.

**Figure S1.**
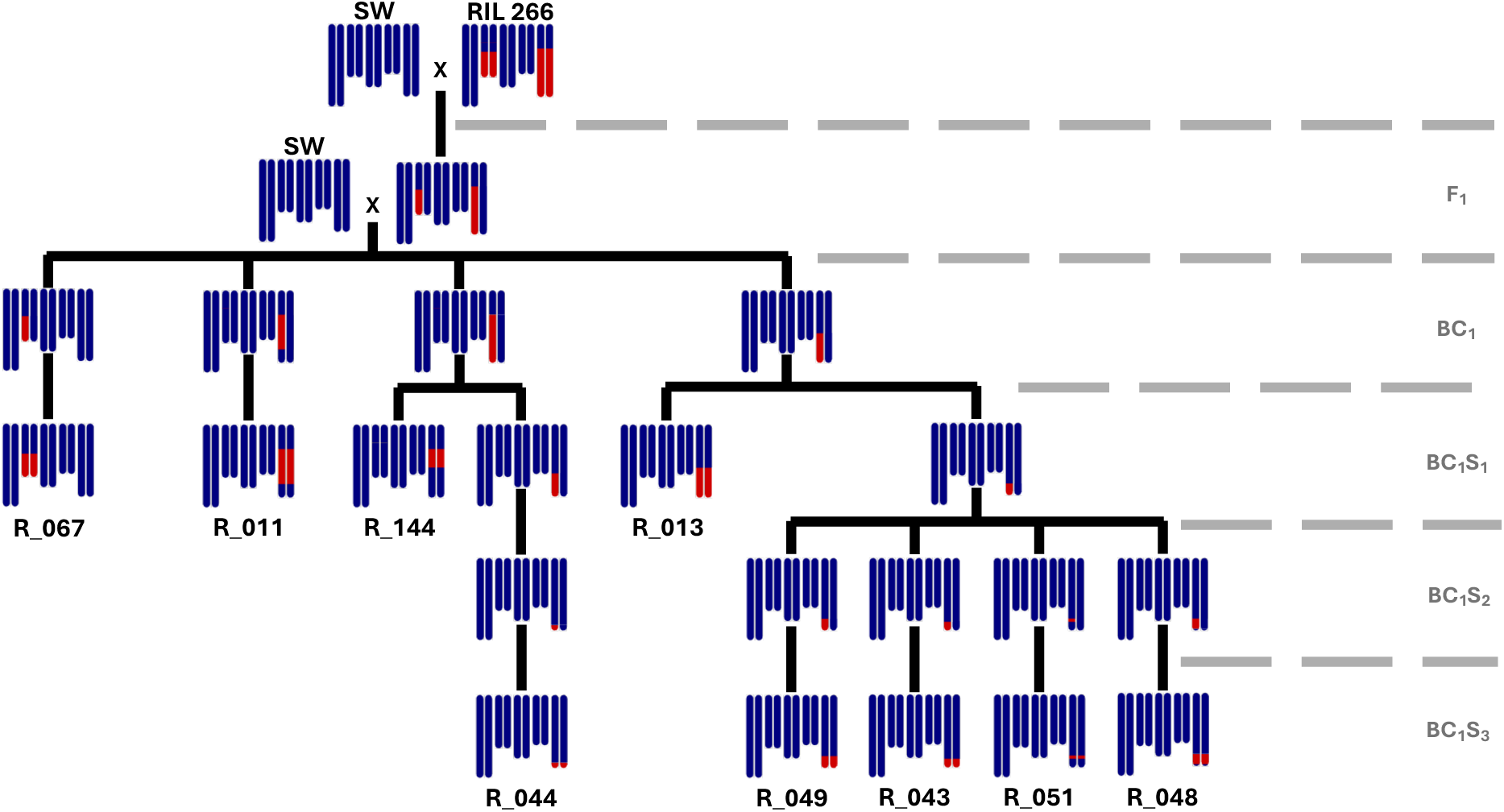
Pedigree illustrating the NIL making process. Genotypes of all NIL descendants included in the manuscript, and some intermediate descendants of RIL 266 are shown. IT genotypes are red; SW genotypes are blue. NIL genotypes included in this paper and genotyped based on WGS data are labeled below their image. Dotted grey lines separate distinct generations during NIL production. In this example, RIL 266 was crossed to SW, to form a RIL F_1_, which was backcrossed to SW, producing the BC_1_ generation. Multiple BC_1_ individuals were then self-pollinated and genotyped over multiple generations (BC_1_S_1_ – BC_1_S_3_) to produce subsequent NILs.

**Figure S2.** Detailed genotyping of all NILs. **(A)** Rolling average of genotype proportion (IT: red, SW: blue, heterozygous: gold) in non-overlapping 100 SNP windows and non-overlapping 25kb windows across the 5 chromosomes of the *A. thaliana* genome. Heatmaps above show the density of centromeric repeats (grey) and density of diagnostic SNP markers (green) across the genome. **(B)** Zoomed in view of the introgressed region of interest, rolling average of genotype proportion (IT: red, SW: blue, heterozygous: gold) in non-overlapping 25kb windows, with a summary haplotype map(s) above. Crossover regions between SNP markers (grey) are also shown.

**Figure S3.**
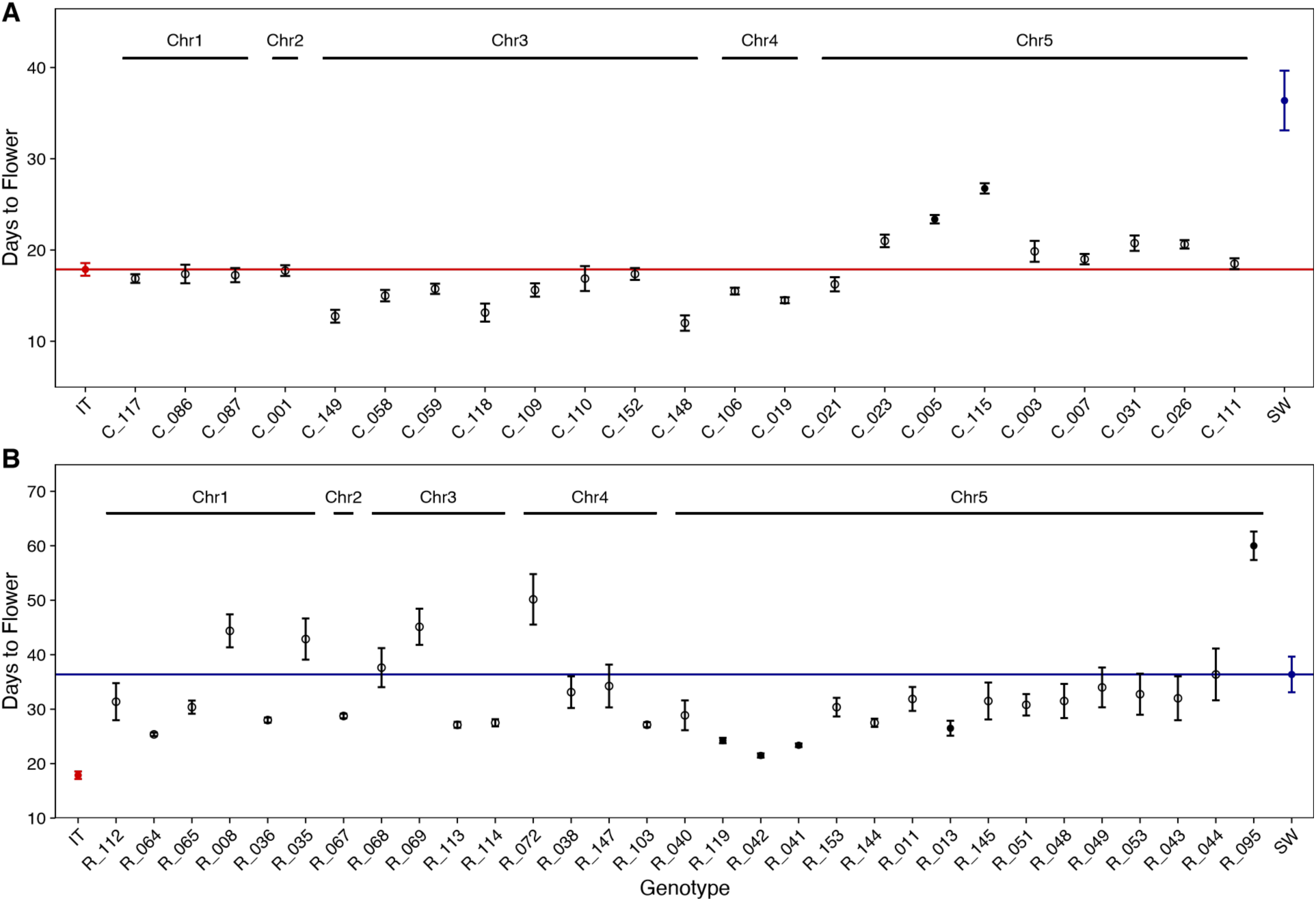
Genotype mean days to first flowering of ecotypes and NILs. **(A)** Mean days to flower +/-SE of ecotypes, IT (red), SW (blue), and NILs in the IT background (black). **(B)** Ecotypes and NILs in the SW background. Solid dots indicate a significant difference from its recurrent ecotype (p<0.05, corrected for multiple testing by a post hoc Dunnett’s test). Note that panels **(A)** and **(B)** have different y-axis scales.

**Figure S4.**
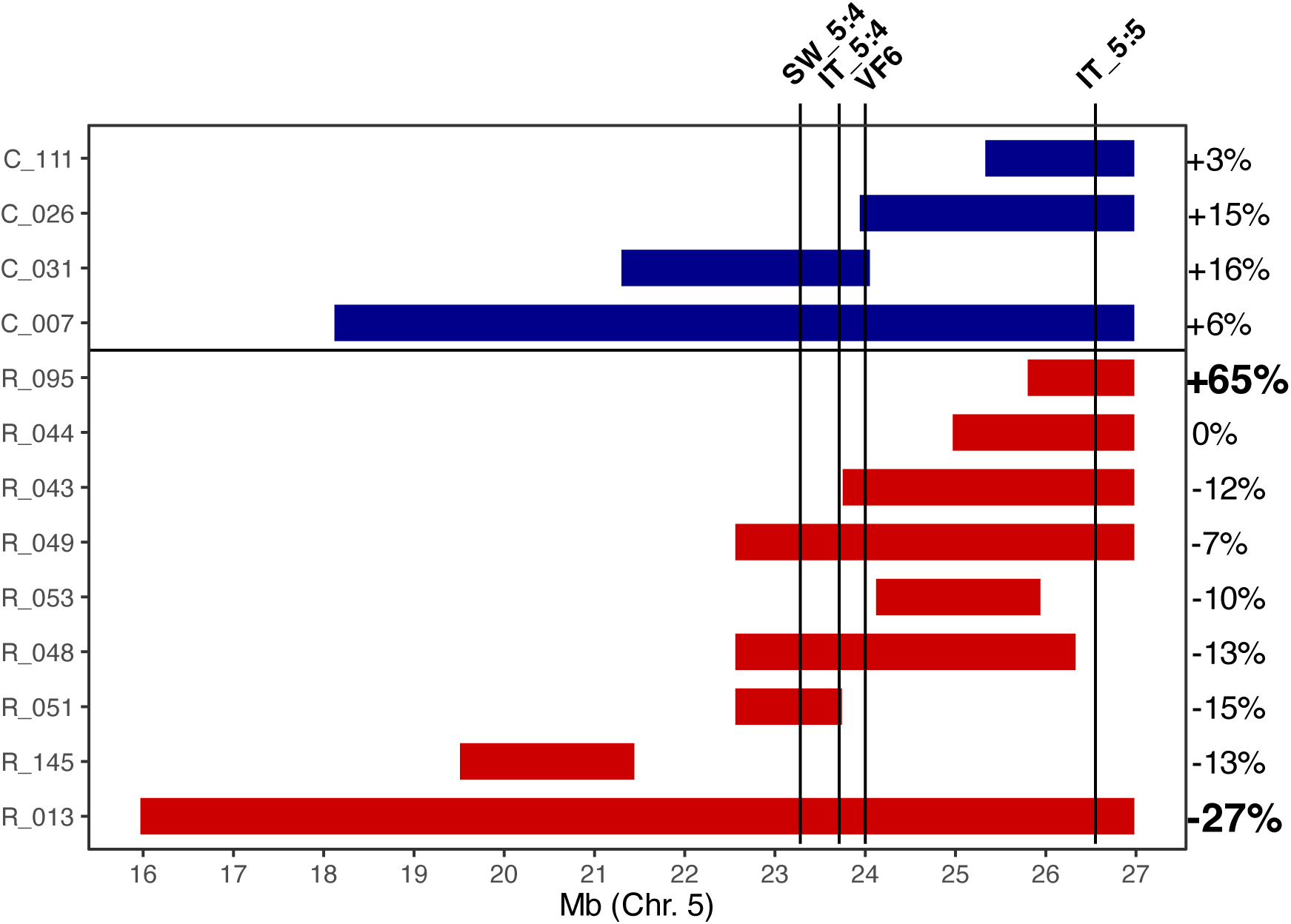
NILs at the bottom of chromosome 5 and previously identified flowering time QTL. Introgressed regions at the end of Chr. 5 are shown for four IT background NILs (blue introgression blocks), and nine SW background NILs (red introgression blocks). The positions of flowering time QTL identified in Grillo *et al*., 2013 (VF6) and Ågren *et al*., 2017 (SW_5:4, IT_5:4, and IT_5:5) are shown. Note that the linkage map used in Grillo *et al*., 2013 was not published, so the physical position of QTL VF6 is estimated from the information provided in the paper. Additionally, the linkage map used in Grillo *et al*., 2013 is slightly truncated, with its last marker upstream of the position of QTL IT_5:5 identified in Ågren *et al*., 2017, and not including the last ∼1Mb of Chr5 (a large portion of the region of interest, introgressed in R_095). At the far right the proportional effect of each introgressed region on flowering time is shown. Significant effects (p<0.05, corrected for multiple testing from its recurrent parent by Dunnett’s test) are shown in bold.

**Figure S5.**
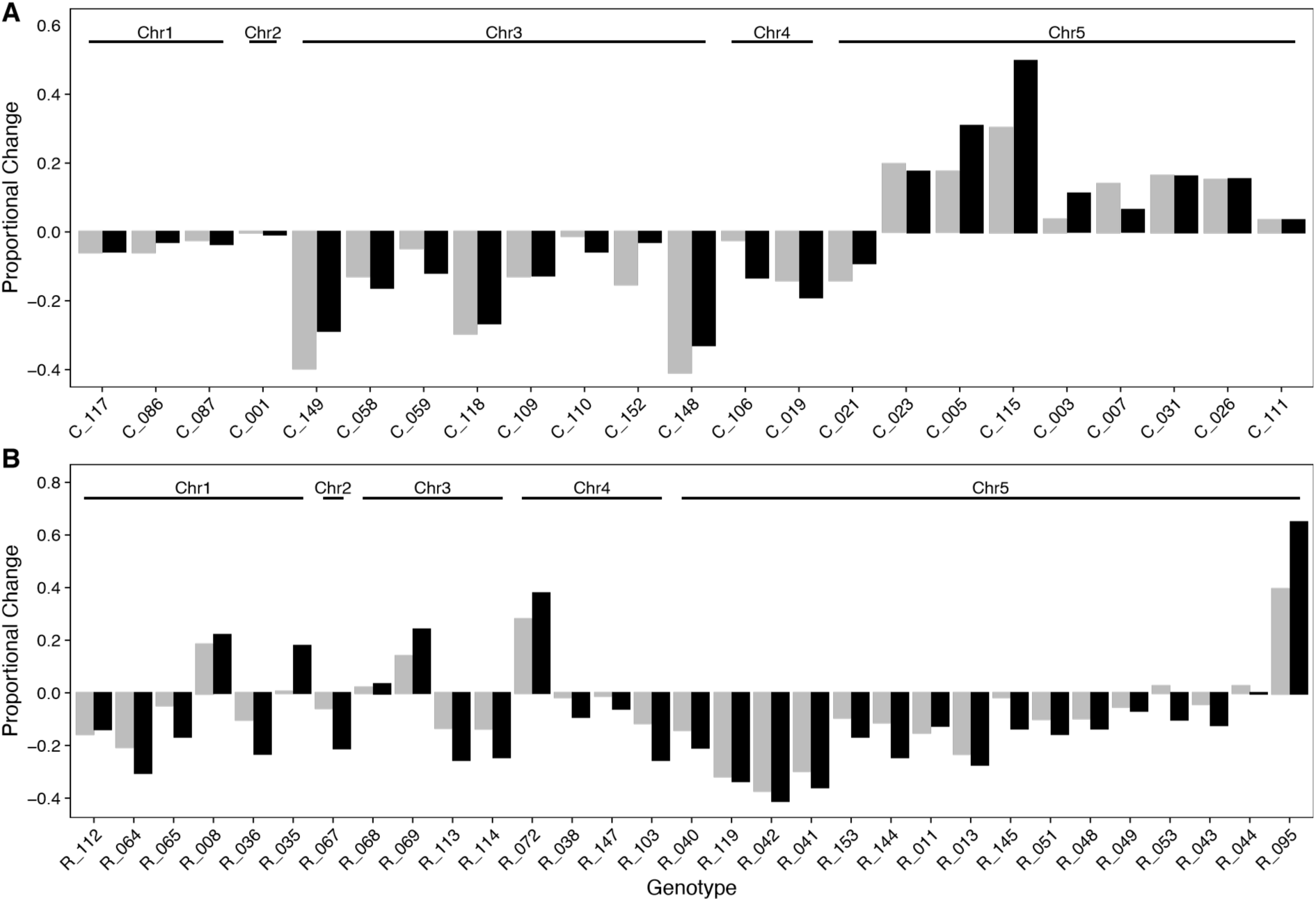
Bolting time and flowering time effect sizes. **(A)** Proportional effect size (NIL compared to IT) of bolting time (grey) and flowering time (black) for each NIL in the IT background. **(B)** Proportional effect size for each NIL in the SW background. Positive values indicate slower bolting or flowering than the recurrent parent, negative values indicate faster bolting or flowering.

**Figure S6.**
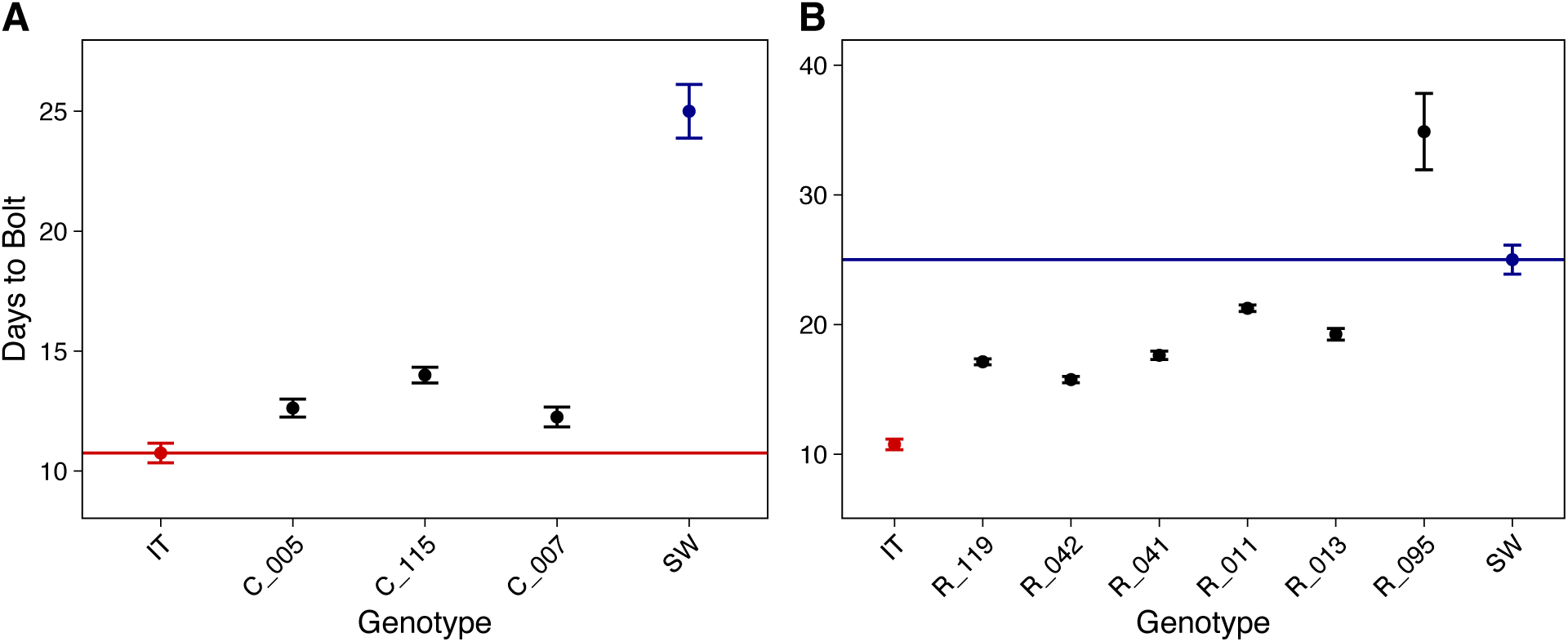
Genotype mean days to bolt of NILs significantly different than their recurrent ecotype. p<0.05, corrected for multiple testing by a post hoc Dunnett’s test. **(A)** Mean days to bolt +/- SE of ecotypes, IT (red), SW (blue), and NILs in the IT background (black). **(B)** Ecotypes and NILs in the SW background. Note that panels **(A)** and **(B)** have different y-axis scales.

**Figure S7.**
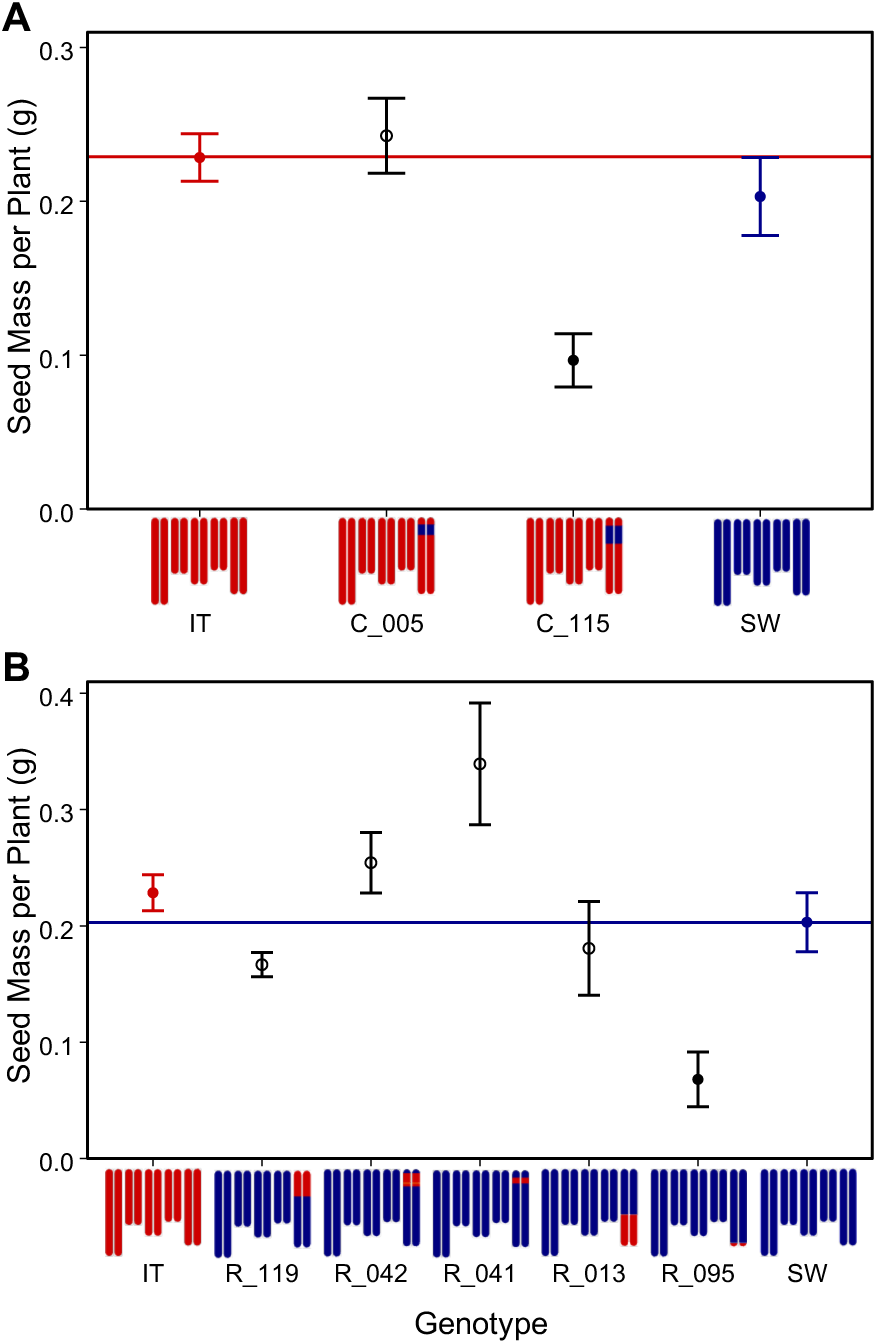
Genotype mean total seed mass per plant for NILs with significantly different first flowering time than their recurrent ecotype (see Figure 4). NILs with significantly different seed mass per plant than their recurrent ecotype (p<0.05, corrected for multiple testing) by a post hoc Dunnett’s test are shown as filled dots. **(A)** Mean seed mass per plant +/- SE of ecotypes, IT (red), SW (blue), and NILs in the IT background (black). **(B)** Ecotypes and NILs in the SW background. All NILs are shown in Fig. S8. Note that panels **(A)** and **(B)** have different y-axis scales.

**Figure S8.**
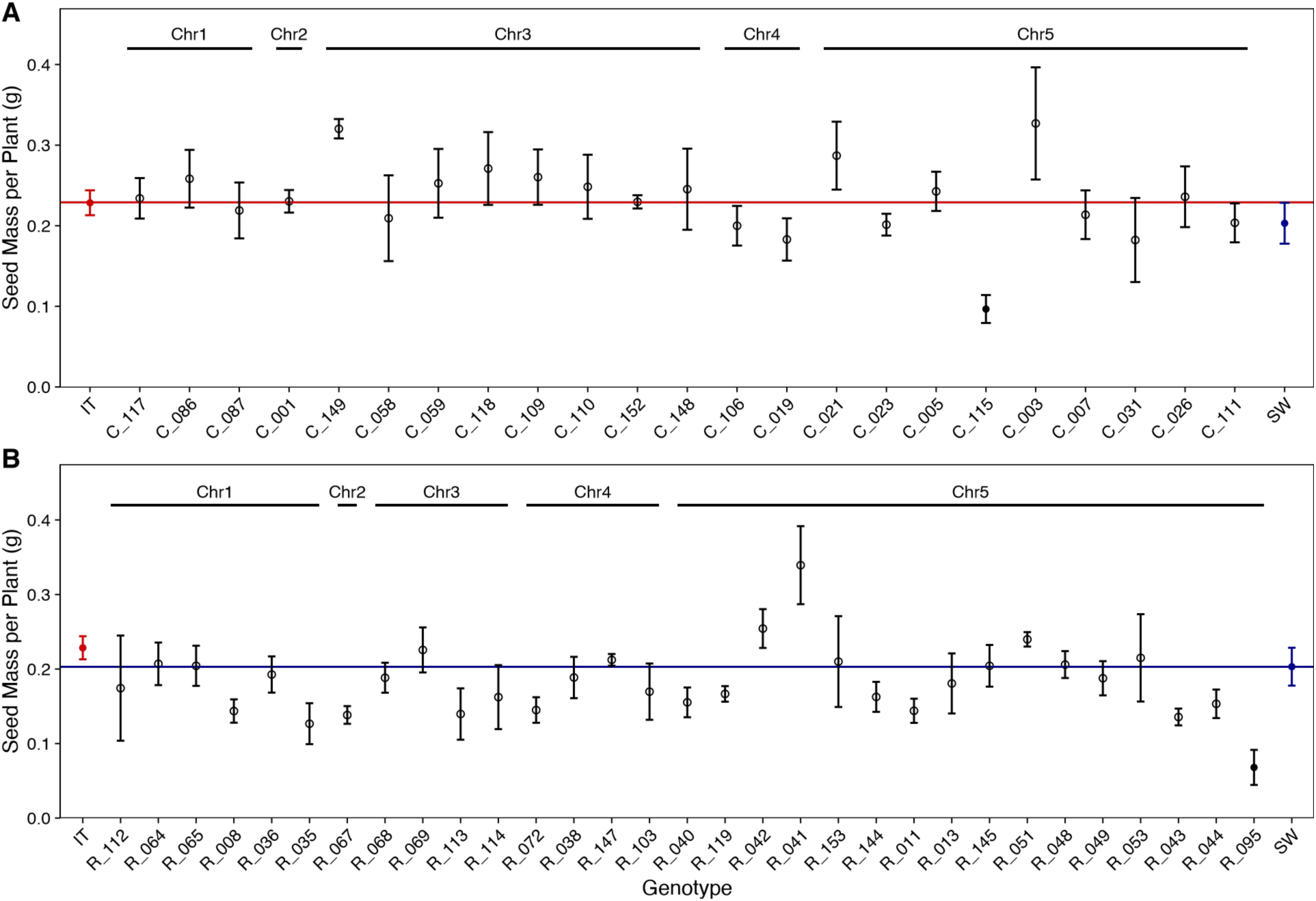
Genotype mean total seed mass per plant of NILs. **(A)** Mean total seed mass +/- SE of ecotypes, IT (red), SW (blue), and NILs in the IT background (black). **(B)** Ecotypes and NILs in the SW background. Solid dots indicate a significant difference (p<0.05, corrected for multiple testing) from its recurrent ecotype by a post hoc Dunnett’s test.

## References

Ågren J, Oakley CG, Lundemo S, Schemske DW. 2017. Adaptive divergence in flowering time among natural populations of *Arabidopsis thaliana*: Estimates of selection and QTL mapping. Evolution 71: 550–564.

Ågren J, Oakley CG, McKay JK, Lovell JT, Schemske DW. 2013. Genetic mapping of adaptation reveals fitness tradeoffs in *Arabidopsis thaliana*. Proceedings of the National Academy of Sciences 110: 21077–21082.

Ågren J, Schemske DW. 2012. Reciprocal transplants demonstrate strong adaptive differentiation of the model organism *Arabidopsis thaliana* in its native range. New Phytologist 194: 1112–1122.

Ali ML, Sanchez PL, Yu S, Lorieux M, Eizenga GC. 2010. Chromosome Segment Substitution Lines: A Powerful Tool for the Introgression of Valuable Genes from *Oryza* Wild Species into Cultivated Rice (*O. sativa*). Rice 3: 218–234.

Alonso-Blanco C, Andrade J, Becker C, Bemm F, Bergelson J, Borgwardt KM, Cao J, Chae E, Dezwaan TM, Ding W, et al. 2016. 1,135 Genomes Reveal the Global Pattern of Polymorphism in *Arabidopsis thaliana*. Cell 166: 481–491.

Alonso-Blanco C, Bentsink L, Hanhart CJ, Vries HB, Koornneef M. 2003. Analysis of Natural Allelic Variation at Seed Dormancy Loci of *Arabidopsis thaliana*. Genetics 164: 711–729.

Amasino R. 2010. Seasonal and developmental timing of flowering. The Plant Journal 61: 1001–1013.

Anderson JT, Willis JH, Mitchell-Olds T. 2011. Evolutionary genetics of plant adaptation. Trends in Genetics 27: 258–266.

Bates D, Mächler M, Bolker B, Walker S. 2015. Fitting Linear Mixed-Effects Models Using lme4. Journal of Statistical Software 67: 1–48.

Beck JB, Schmuths H, Schaal BA. 2008. Native range genetic variation in *Arabidopsis thaliana* is strongly geographically structured and reflects Pleistocene glacial dynamics. Molecular Ecology 17: 902–915.

Bentsink L, Yuan K, Koornneef M, Vreugdenhil D. 2003. The genetics of phytate and phosphate accumulation in seeds and leaves of *Arabidopsis thaliana*, using natural variation. Theoretical and Applied Genetics 106: 1234–1243.

Bolger AM, Lohse M, Usadel B. 2014. Trimmomatic: a flexible trimmer for Illumina sequence data. Bioinformatics 30: 2114–2120.

Bomblies K, Peichel CL. 2022. Genetics of adaptation. Proceedings of the National Academy of Sciences 119: e2122152119.

Bouvet J-M, Saya A, Vigneron Ph. 2009. Trends in additive, dominance and environmental effects with age for growth traits in *Eucalyptus* hybrid populations. Euphytica 165: 35–54.

Catchen J, Hohenlohe PA, Bassham S, Amores A, Cresko WA. 2013. Stacks: an analysis tool set for population genomics. Molecular Ecology 22: 3124–3140.

Chen H, Yuan Y-W. 2024. Genetic basis of nectar guide trichome variation between bumblebee- and self-pollinated monkeyflowers (*Mimulus*): role of the MIXTA-like gene GUIDELESS. BMC Plant Biology 24: 62.

Chitwood DH, Kumar R, Headland LR, Ranjan A, Covington MF, Ichihashi Y, Fulop D, Jiménez-Gómez JM, Peng J, Maloof JN, et al. 2013. A Quantitative Genetic Basis for Leaf Morphology in a Set of Precisely Defined Tomato Introgression Lines. The Plant Cell 25: 2465–2481.

Danecek P, Auton A, Abecasis G, Albers CA, Banks E, DePristo MA, Handsaker RE, Lunter G, Marth GT, Sherry ST, et al. 2011. The variant call format and VCFtools. Bioinformatics 27: 2156–2158.

Danecek P, Bonfield JK, Liddle J, Marshall J, Ohan V, Pollard MO, Whitwham A, Keane T, McCarthy SA, Davies RM, et al. 2021. Twelve years of SAMtools and BCFtools. GigaScience 10: giab008.

Des Marais DL, Hernandez KM, Juenger TE. 2013. Genotype-by-Environment Interaction and Plasticity: Exploring Genomic Responses of Plants to the Abiotic Environment. Annual Review of Ecology, Evolution, and Systematics 44: 5–29.

Dittmar EL, Oakley CG, Ågren J, Schemske DW. 2014. Flowering time QTL in natural populations of *Arabidopsis thaliana* and implications for their adaptive value. Molecular Ecology 23: 4291–4303.

Dittmar EL, Oakley CG, Conner JK, Gould BA, Schemske DW. 2016. Factors influencing the effect size distribution of adaptive substitutions. Proceedings of the Royal Society B: Biological Sciences.

Dittrich-Reed DR, Fitzpatrick BM. 2013. Transgressive Hybrids as Hopeful Monsters. Evolutionary Biology 40: 310–315.

Durvasula A, Fulgione A, Gutaker RM, Alacakaptan SI, Flood PJ, Neto C, Tsuchimatsu T, Burbano HA, Picó FX, Alonso-Blanco C, et al. 2017. African genomes illuminate the early history and transition to selfing in *Arabidopsis thaliana*. Proceedings of the National Academy of Sciences 114: 5213–5218.

Eduardo I, Arús P, Monforte AJ. 2005. Development of a genomic library of near isogenic lines (NILs) in melon (*Cucumis melo* L.) from the exotic accession PI161375. Theoretical and Applied Genetics 112: 139–148.

Edwards K, Johnstone C, Thompson C. 1991. A simple and rapid method for the preparation of plant genomic DNA for PCR analysis. Nucleic Acids Research 19: 1349.

Edwards KD, Lynn JR, Gyula P, Nagy F, Millar AJ. 2005. Natural Allelic Variation in the Temperature-Compensation Mechanisms of the *Arabidopsis thaliana* Circadian Clock. Genetics 170: 387–400.

Ellis TJ, Postma FM, Oakley CG, Ågren J. 2021. Life-history trade-offs and the genetic basis of fitness in *Arabidopsis thaliana*. Molecular Ecology 30: 2846–2858.

Eshed Y, Zamir D. 1994. A genomic library of *Lycopersicon pennellii* in *L. esculentum*: A tool for fine mapping of genes. Euphytica 79: 175–179.

Eshed Y, Zamir D. 1995. An introgression line population of *Lycopersicon pennellii* in the cultivated tomato enables the identification and fine mapping of yield-associated QTL. Genetics 141: 1147–1162.

Fishman L, Sweigart AL, Kenney AM, Campbell S. 2014. Major quantitative trait loci control divergence in critical photoperiod for flowering between selfing and outcrossing species of monkeyflower (*Mimulus*). The New Phytologist 201: 1498–1507.

Fletcher RS, Mullen JL, Yoder S, Bauerle WL, Reuning G, Sen S, Meyer E, Juenger TE, McKay JK. 2013. Development of a next-generation NIL library in *Arabidopsis thaliana* for dissecting complex traits. BMC Genomics 14: 655.

Gnan S, Priest A, Kover PX. 2014. The Genetic Basis of Natural Variation in Seed Size and Seed Number and Their Trade-Off Using *Arabidopsis thaliana* MAGIC Lines. Genetics 198: 1751–1758.

Grillo MA, Li C, Hammond M, Wang L, Schemske DW. 2013. Genetic architecture of flowering time differentiation between locally adapted populations of *Arabidopsis thaliana*. New Phytologist 197: 1321–1331.

Guerrero RF, Muir CD, Josway S, Moyle LC. 2017. Pervasive antagonistic interactions among hybrid incompatibility loci. PLOS Genetics 13: e1006817.

Hepworth J, Antoniou-Kourounioti RL, Berggren K, Selga C, Tudor EH, Yates B, Cox D, Collier Harris BR, Irwin JA, Howard M, et al. 2020. Natural variation in autumn expression is the major adaptive determinant distinguishing *Arabidopsis* FLC haplotypes. eLife 9: e57671.

Hereford J. 2009. A Quantitative Survey of Local Adaptation and Fitness Trade-Offs. The American Naturalist 173: 579–588.

Herridge RP, Day RC, Baldwin S, Macknight RC. 2011. Rapid analysis of seed size in *Arabidopsis* for mutant and QTL discovery. Plant Methods 7: 3.

Jeuken MJW, Lindhout P. 2004. The development of lettuce backcross inbred lines (BILs) for exploitation of the *Lactuca saligna* (wild lettuce) germplasm. Theoretical and Applied Genetics 109: 394–401.

Josephs EB. 2018. Determining the evolutionary forces shaping G × E. New Phytologist 219: 31–36.

Kalisz S, Wardle GM. 1994. Life history variation in *Campanula americana* (Campanulaceae): population differentiation. American Journal of Botany 81: 521–527.

Kaur A, Khangura RS, Dilkes BP. 2025. A maize mutant in the glutamate receptor-like dwarf13 is modified by cis-acting natural variation and a cornichon homolog. PLOS Genetics 21: e1012006.

Kawecki TJ, Ebert D. 2004. Conceptual issues in local adaptation. Ecology Letters 7: 1225–1241.

Keurentjes JJB, Bentsink L, Alonso-Blanco C, Hanhart CJ, Blankestijn-De Vries H, Effgen S, Vreugdenhil D, Koornneef M. 2007. Development of a Near-Isogenic Line Population of *Arabidopsis thaliana* and Comparison of Mapping Power With a Recombinant Inbred Line Population. Genetics 175: 891–905.

Khangura RS, Marla S, Venkata BP, Heller NJ, Johal GS, Dilkes BP. 2019. A Very Oil Yellow1 Modifier of the Oil Yellow1-N1989 Allele Uncovers a Cryptic Phenotypic Impact of Cis-regulatory Variation in Maize. G3 Genes|Genomes|Genetics 9: 375–390.

Khangura RS, Venkata BP, Marla SR, Mickelbart MV, Dhungana S, Braun DM, Dilkes BP, Johal GS. 2020. Interaction Between Induced and Natural Variation at oil yellow1 Delays Reproductive Maturity in Maize. G3 Genes|Genomes|Genetics 10: 797–810.

Koornneef M, Hanhart CJ, van der Veen JH. 1991. A genetic and physiological analysis of late flowering mutants in *Arabidopsis thaliana*. Molecular and General Genetics MGG 229: 57–66.

Lacey EP. 1988. Latitudinal Variation in Reproductive Timing of a Short-Lived Monocarp, *Daucus Carota* (Apiaceae). Ecology 69: 220–232.

Le Corre V, Roux F, Reboud X. 2002. DNA Polymorphism at the FRIGIDA Gene in *Arabidopsis thaliana*: Extensive Nonsynonymous Variation Is Consistent with Local Selection for Flowering Time. Molecular Biology and Evolution 19: 1261–1271.

Lee G, Sanderson BJ, Ellis TJ, Dilkes BP, McKay JK, Ågren J, Oakley CG. 2024. A large-effect fitness trade-off across environments is explained by a single mutation affecting cold acclimation. Proceedings of the National Academy of Sciences 121: e2317461121.

Leimu R, Fischer M. 2008. A Meta-Analysis of Local Adaptation in Plants. PLOS ONE 3: e4010.

Leinonen PH, Remington DL, Leppälä J, Savolainen O. 2013. Genetic basis of local adaptation and flowering time variation in *Arabidopsis lyrata*. Molecular Ecology 22: 709–723.

Lenth R. 2025. Estimated Marginal Means, aka Least-Squares Means.

Li H. 2013. Aligning sequence reads, clone sequences and assembly contigs with BWA-MEM. *arXiv:1303.3997 [q-bio].*

Li H, Durbin R. 2009. Fast and accurate short read alignment with Burrows–Wheeler transform. Bioinformatics 25: 1754–1760.

Li P, Filiault D, Box MS, Kerdaffrec E, Oosterhout C van, Wilczek AM, Schmitt J, McMullan M, Bergelson J, Nordborg M, et al. 2014a. Multiple FLC haplotypes defined by independent cis-regulatory variation underpin life history diversity in Arabidopsis thaliana. Genes & Development 28: 1635–1640.

Li H, Handsaker B, Wysoker A, Fennell T, Ruan J, Homer N, Marth G, Abecasis G, Durbin R, 1000 Genome Project Data Processing Subgroup. 2009. The Sequence Alignment/Map format and SAMtools. Bioinformatics 25: 2078–2079.

Li C-Q, Song L, Zhao H-H, Wang Q-L, Fu Y-Z. 2014b. Identification of quantitative trait loci with main and epistatic effects for plant architecture traits in Upland cotton (*Gossypium hirsutum* L.). Plant Breeding 133: 390–400.

Lippman ZB, Semel Y, Zamir D. 2007. An integrated view of quantitative trait variation using tomato interspecific introgression lines. Current Opinion in Genetics & Development 17: 545–552.

Liu S, Zhou R, Dong Y, Li P, Jia J. 2006. Development, utilization of introgression lines using a synthetic wheat as donor. Theoretical and Applied Genetics 112: 1360–1373.

Lopez L, Lang PLM, Marciniak S, Kistler L, Latorre SM, Haile A, Cerda EV, Gamba D, Xu Y, Woods P, et al. 2025. Museum genomics reveals temporal genetic stasis and global genetic diversity in *Arabidopsis thaliana*. : 2025.02.06.636844.

Mackay TFC. 2014. Epistasis and quantitative traits: using model organisms to study gene–gene interactions. Nature Reviews Genetics 15: 22–33.

Mackay TFC, Anholt RRH. 2024. Pleiotropy, epistasis and the genetic architecture of quantitative traits. Nature Reviews Genetics 25: 639–657.

Mantel SJ, Sweigart AL. 2024. Postzygotic barriers persist despite ongoing introgression in hybridizing *Mimulus* species. Molecular Ecology n/a: e17261.

Marshall MM, Remington DL, Lacey EP. 2020. Two reproductive traits show contrasting genetic architectures in *Plantago lanceolata*. Molecular Ecology 29: 272–291.

McKenna A, Hanna M, Banks E, Sivachenko A, Cibulskis K, Kernytsky A, Garimella K, Altshuler D, Gabriel S, Daly M, et al. 2010. The Genome Analysis Toolkit: A MapReduce framework for analyzing next-generation DNA sequencing data. Genome Research 20: 1297–1303.

Melchinger AE, Utz HF, Piepho H-P, Zeng Z-B, Schön CC. 2007. The Role of Epistasis in the Manifestation of Heterosis: A Systems-Oriented Approach. Genetics 177: 1815–1825.

Mojica JP, Mullen J, Lovell JT, Monroe JG, Paul JR, Oakley CG, McKay JK. 2016. Genetics of water use physiology in locally adapted *Arabidopsis thaliana*. Plant Science 251: 12–22.

Monforte AJ, Tanksley SD. 2000. Development of a set of near isogenic and backcross recombinant inbred lines containing most of the *Lycopersicon hirsutum* genome in a *L. esculentum* genetic background: A tool for gene mapping and gene discovery. Genome 43: 803–813.

Muir CD, Moyle LC. 2009. Antagonistic epistasis for ecophysiological trait differences between *Solanum* species. New Phytologist 183: 789–802.

Oakley CG, Ågren J, Atchison RA, Schemske DW. 2014. QTL mapping of freezing tolerance: links to fitness and adaptive trade-offs. Molecular Ecology 23: 4304–4315.

Oakley CG, Ågren J, Schemske DW. 2015. Heterosis and outbreeding depression in crosses between natural populations of *Arabidopsis thaliana*. Heredity 115: 73–82.

Oakley CG, Lundemo S, Ågren J, Schemske DW. 2019. Heterosis is common and inbreeding depression absent in natural populations of *Arabidopsis thaliana*. Journal of Evolutionary Biology 32: 592–603.

Oakley CG, Savage L, Lotz S, Larson GR, Thomashow MF, Kramer DM, Schemske DW. 2018. Genetic basis of photosynthetic responses to cold in two locally adapted populations of *Arabidopsis thaliana*. Journal of Experimental Botany 69: 699–709.

Oakley CG, Schemske DW, McKay JK, Ågren J. 2023. Ecological genetics of local adaptation in *Arabidopsis*: An 8-year field experiment. Molecular Ecology 32: 4570–4583.

Okonechnikov K, Conesa A, García-Alcalde F. 2016. Qualimap 2: advanced multi-sample quality control for high-throughput sequencing data. Bioinformatics 32: 292–294.

Olsson K, Ågren J. 2002. Latitudinal population differentiation in phenology, life history and flower morphology in the perennial herb *Lythrum salicaria*. Journal of Evolutionary Biology 15: 983–996.

Orr HA. 1998. The Population Genetics of Adaptation: The Distribution of Factors Fixed During Adaptive Evolution. Evolution 52: 935–949.

Pabuayon ICM, Kitazumi A, Cushman KR, Singh RK, Gregorio GB, Dhatt B, Zabet-Moghaddam M, Walia H, de los Reyes BG. 2021. Novel and Transgressive Salinity Tolerance in Recombinant Inbred Lines of Rice Created by Physiological Coupling-Uncoupling and Network Rewiring Effects. Frontiers in Plant Science 12.

Phillips PC. 2008. Epistasis — the essential role of gene interactions in the structure and evolution of genetic systems. Nature Reviews Genetics 9: 855–867.

Postma FM, Ågren J. 2015. Maternal environment affects the genetic basis of seed dormancy in *Arabidopsis thaliana*. Molecular Ecology 24: 785–797.

Postma FM, Ågren J. 2016. Early life stages contribute strongly to local adaptation in *Arabidopsis thaliana*. Proceedings of the National Academy of Sciences 113: 7590–7595.

Postma FM, Ågren J. 2018. Among-year variation in selection during early life stages and the genetic basis of fitness in *Arabidopsis thaliana*. Molecular Ecology 27: 2498–2511.

Ramsay LD, Jennings DE, Kearsey MJ, Marshall DF, Bohuon EJR, Arthur AE, Lydiate DJ. 1996. The construction of a substitution library of recombinant backcross lines in *Brassica oleracea* for the precision mapping of quantitative trait loci. Genome 39: 558–567.

Reif JC, Kusterer B, Piepho H-P, Meyer RC, Altmann T, Schön CC, Melchinger AE. 2009. Unraveling Epistasis With Triple Testcross Progenies of Near-Isogenic Lines. Genetics 181: 247–257.

Rieseberg LH, Archer MA, Wayne RK. 1999. Transgressive segregation, adaptation and speciation. Heredity 83: 363–372.

Rieseberg LH, Raymond O, Rosenthal DM, Lai Z, Livingstone K, Nakazato T, Durphy JL, Schwarzbach AE, Donovan LA, Lexer C. 2003. Major Ecological Transitions in Wild Sunflowers Facilitated by Hybridization. Science 301: 1211–1216.

Robinson JT, Thorvaldsdóttir H, Winckler W, Guttman M, Lander ES, Getz G, Mesirov JP. 2011. Integrative genomics viewer. Nature Biotechnology 29: 24–26.

Rojas-Gutierrez JD, Mantel SJ, Oakley CG. 2026. Environment-dependent and often antagonistic effects of dominance and epistasis on heterosis in crosses between natural populations. BioRxiv, Preprint

Sanderson BJ, Park S, Jameel MI, Kraft JC, Thomashow MF, Schemske DW, Oakley CG. 2020. Genetic and physiological mechanisms of freezing tolerance in locally adapted populations of a winter annual. American Journal of Botany 107: 250–261.

Schindelin J, Arganda-Carreras I, Frise E, Kaynig V, Longair M, Pietzsch T, Preibisch S, Rueden C, Saalfeld S, Schmid B, et al. 2012. Fiji: an open-source platform for biological-image analysis. Nature Methods 9: 676–682.

Schluter D, Marchinko KB, Arnegard ME, Zhang H, Brady SD, Jones FC, Bell MA, Kingsley DM. 2021. Fitness maps to a large-effect locus in introduced stickleback populations. Proceedings of the National Academy of Sciences 118: e1914889118.

Schmalenbach I, March TJ, Bringezu T, Waugh R, Pillen K. 2011. High-Resolution Genotyping of Wild Barley Introgression Lines and Fine-Mapping of the Threshability Locus thresh-1 Using the Illumina GoldenGate Assay. G3 Genes|Genomes|Genetics 1: 187–196.

Shindo C, Lister C, Crevillen P, Nordborg M, Dean C. 2006. Variation in the epigenetic silencing of FLC contributes to natural variation in *Arabidopsis* vernalization response. Genes & Development 20: 3079–3083.

Shuang LS, Cuevas H, Lemke C, Kim C, Shehzad T, Paterson AH. 2023. Genetic dissection of morphological variation between cauliflower and a rapid cycling *Brassica oleracea* line. G3 Genes|Genomes|Genetics 13: jkad163.

Simpson GG, Dean C. 2002. *Arabidopsis*, the Rosetta Stone of Flowering Time? Science 296: 285–289.

Stinchcombe JR, Hoekstra HE. 2008. Combining population genomics and quantitative genetics: finding the genes underlying ecologically important traits. Heredity 100: 158–170.

Sung S, Amasino RM. 2004. Vernalization and epigenetics: how plants remember winter. Current Opinion in Plant Biology 7: 4–10.

Szalma SJ, Hostert BM, LeDeaux JR, Stuber CW, Holland JB. 2007. QTL mapping with near-isogenic lines in maize. Theoretical and Applied Genetics 114: 1211–1228.

Törjék O, Meyer RC, Zehnsdorf M, Teltow M, Strompen G, Witucka-Wall H, Blacha A, Altmann T. 2008. Construction and Analysis of 2 Reciprocal *Arabidopsis* Introgression Line Populations. Journal of Heredity 99: 396–406.

Ungerer MC. 2005. A Primer of Ecological Genetics. BioScience 55: 283–285.

Van Daele I, Gonzalez N, Vercauteren I, de Smet L, Inzé D, Roldán-Ruiz I, Vuylsteke M. 2012. A comparative study of seed yield parameters in *Arabidopsis thaliana* mutants and transgenics. Plant Biotechnology Journal 10: 488–500.

VanWallendael A, Soltani A, Emery NC, Peixoto MM, Olsen J, Lowry DB. 2019. A Molecular View of Plant Local Adaptation: Incorporating Stress-Response Networks. Annual Review of Plant Biology 70: 559–583.

Wadgymar SM, DeMarche ML, Josephs EB, Sheth SN, Anderson JT. 2022. Local Adaptation: Causal Agents of Selection and Adaptive Trait Divergence. Annual Review of Ecology, Evolution, and Systematics 53: 87–111.

Wang Y, Li W, Wang L, Yan J, Lu G, Yang N, Xu J, Wang Y, Gui S, Chen G, et al. 2022. Three types of genes underlying the Gametophyte factor1 locus cause unilateral cross incompatibility in maize. Nature Communications 13: 4498.

Wang S, Meyer E, McKay JK, Matz MV. 2012. 2b-RAD: a simple and flexible method for genome-wide genotyping. Nature Methods 9: 808–810.

Wardyn BM, Edwards JW, Lamkey KR. 2007. The Genetic Structure of a Maize Population: The Role of Dominance. Crop Science 47: 467–474.

Whittaker C, Dean C. 2017. The FLC Locus: A Platform for Discoveries in Epigenetics and Adaptation. Annual Review of Cell and Developmental Biology 33: 555–575.

Wijnen CL, Botet R, van de Belt J, Deurhof L, de Jong H, de Snoo CB, Dirks R, Boer MP, van Eeuwijk FA, Wijnker E, et al. 2024. A complete chromosome substitution mapping panel reveals genome-wide epistasis in *Arabidopsis*. Heredity 133: 198–205.

Yan Z, Liang D, Liu H, Zheng G. 2010. FLC: A key regulator of flowering time in *Arabidopsis*. Russian Journal of Plant Physiology 57: 166–174.

Yang Z, Jin L, Zhu H, Wang S, Zhang G, Liu G. 2018. Analysis of Epistasis among QTLs on Heading Date based on Single Segment Substitution Lines in Rice. Scientific Reports 8: 3059.

Zhao L, Zhou H, Lu L, Liu L, Li X, Lin Y, Yu S. 2009. Identification of quantitative trait loci controlling rice mature seed culturability using chromosomal segment substitution lines. Plant Cell Reports 28: 247–256.

Zhu P, Schon M, Questa J, Nodine M, Dean C. 2023. Causal role of a promoter polymorphism in natural variation of the *Arabidopsis* floral repressor gene FLC. Current Biology 33: 4381–4391.e3.

