## Supplementary material for "A panel of near-isogenic lines derived from locally adapted populations of a wild plant: A powerful tool for dissecting additive and non-additive effects on ecologically important traits": Fig. S2

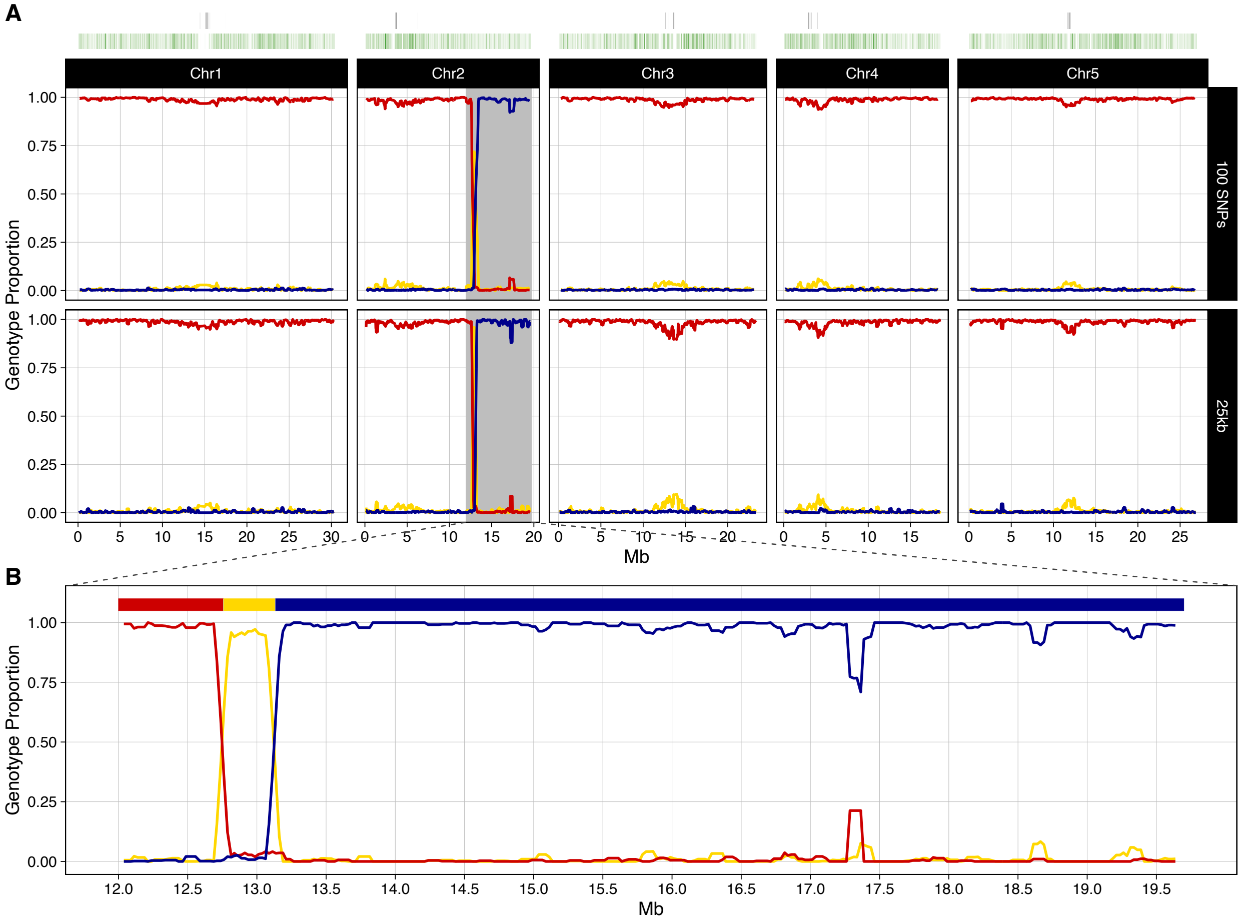
A
Chr1
Chr2
Chr3
Chr4
Chr5
100 SNPs
25kb
Genotype Proportion
1.00
0.75
0.50
0.25
0.00
Mb
B
12.0
12.5
13.0
13.5
14.0
14.5
15.0
15.5
16.0
16.5
17.0
17.5
18.0
18.5
19.0
19.5

**C_001**

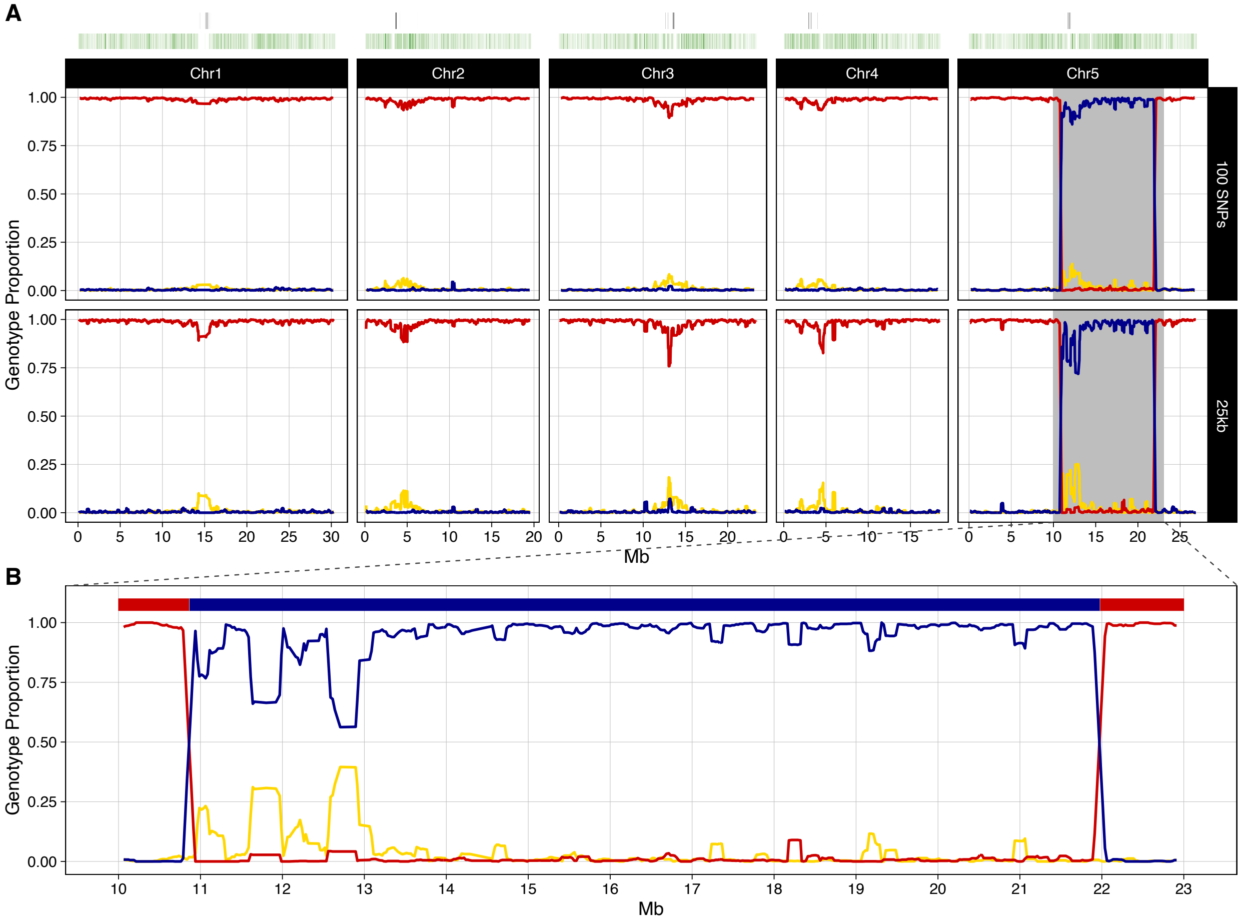
A
Chr1
Chr2
Chr3
Chr4
Chr5
100 SNPs
25kb
Genotype Proportion
1.00
0.75
0.50
0.25
0.00
0
5
10
15
20
25
30
Mb
B
Genotype Proportion
Mb

**C_003**

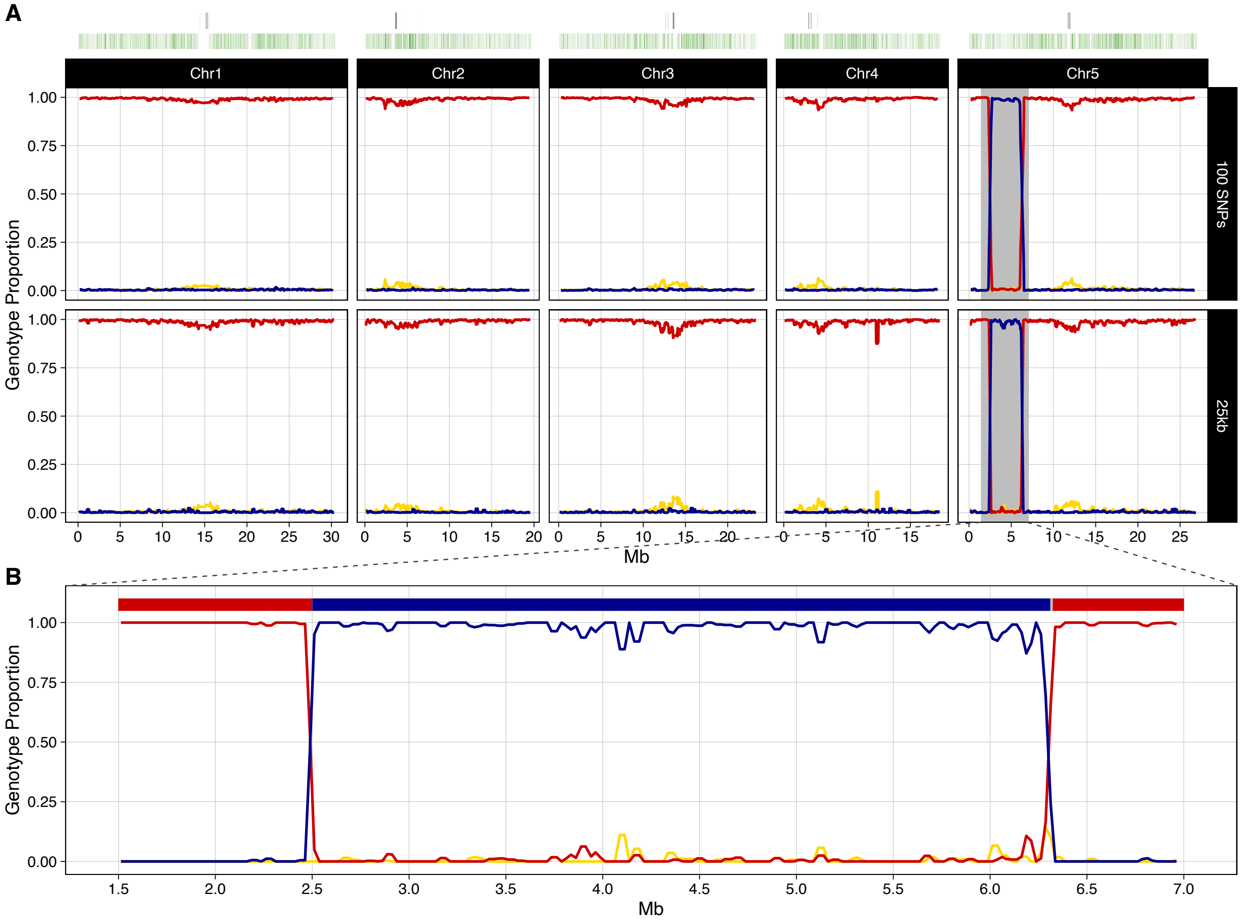
A
Chr1
Chr2
Chr3
Chr4
Chr5
100 SNPs
25kb
Genotype Proportion
1.00
0.75
0.50
0.25
0.00
Mb
B
Genotype Proportion
Mb
1.5
2.0
2.5
3.0
3.5
4.0
4.5
5.0
5.5
6.0
6.5
7.0

**C_005**

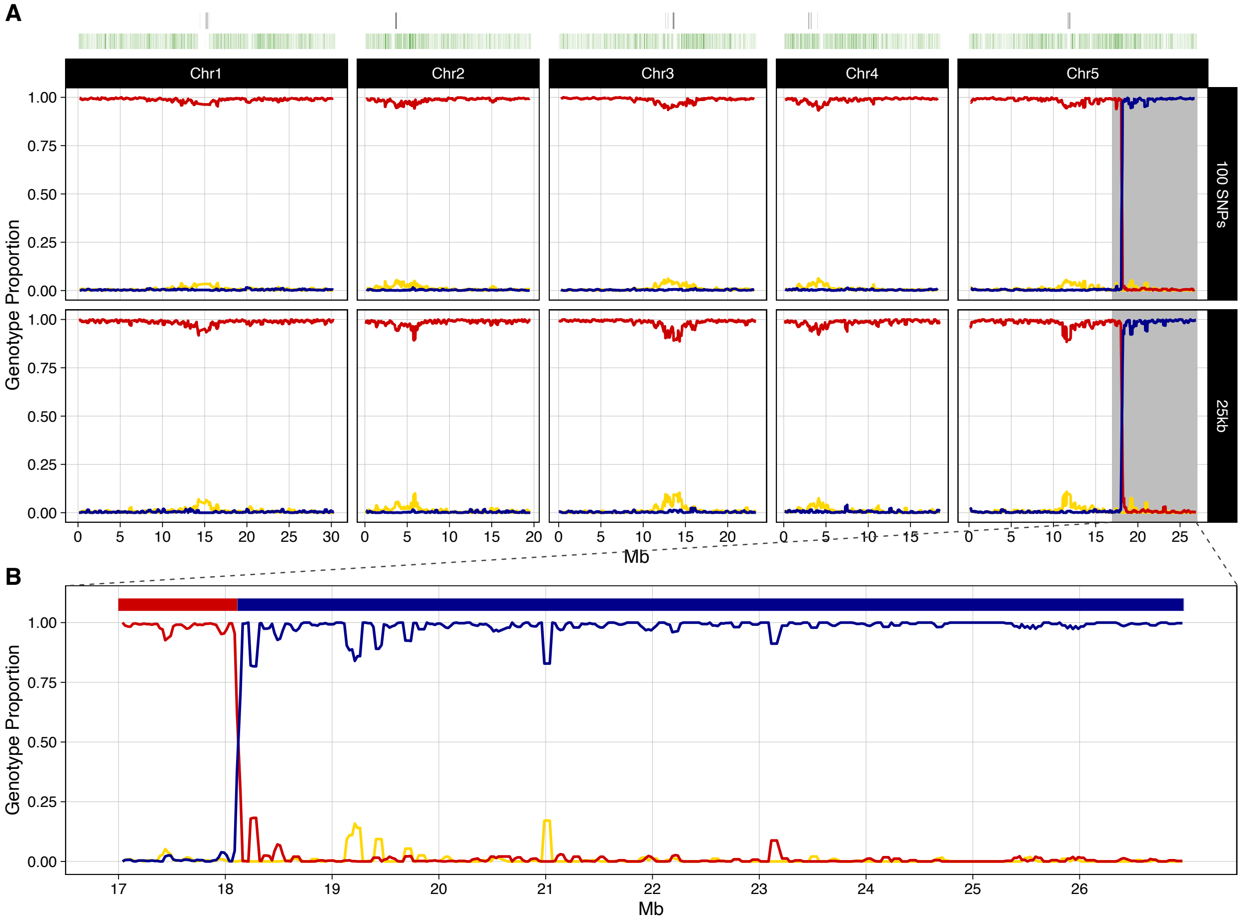
A
Chr1
Chr2
Chr3
Chr4
Chr5
100 SNPs
25kb
Genotype Proportion
1.00
0.75
0.50
0.25
0.00
0 5 10 15 20 25 30
0 5 10 15 20
0 5 10 15 20
0 5 10 15
0 5 10 15 20 25
Mb
B
Genotype Proportion
1.00
0.75
0.50
0.25
0.00
17 18 19 20 21 22 23 24 25 26
Mb

**C_007**

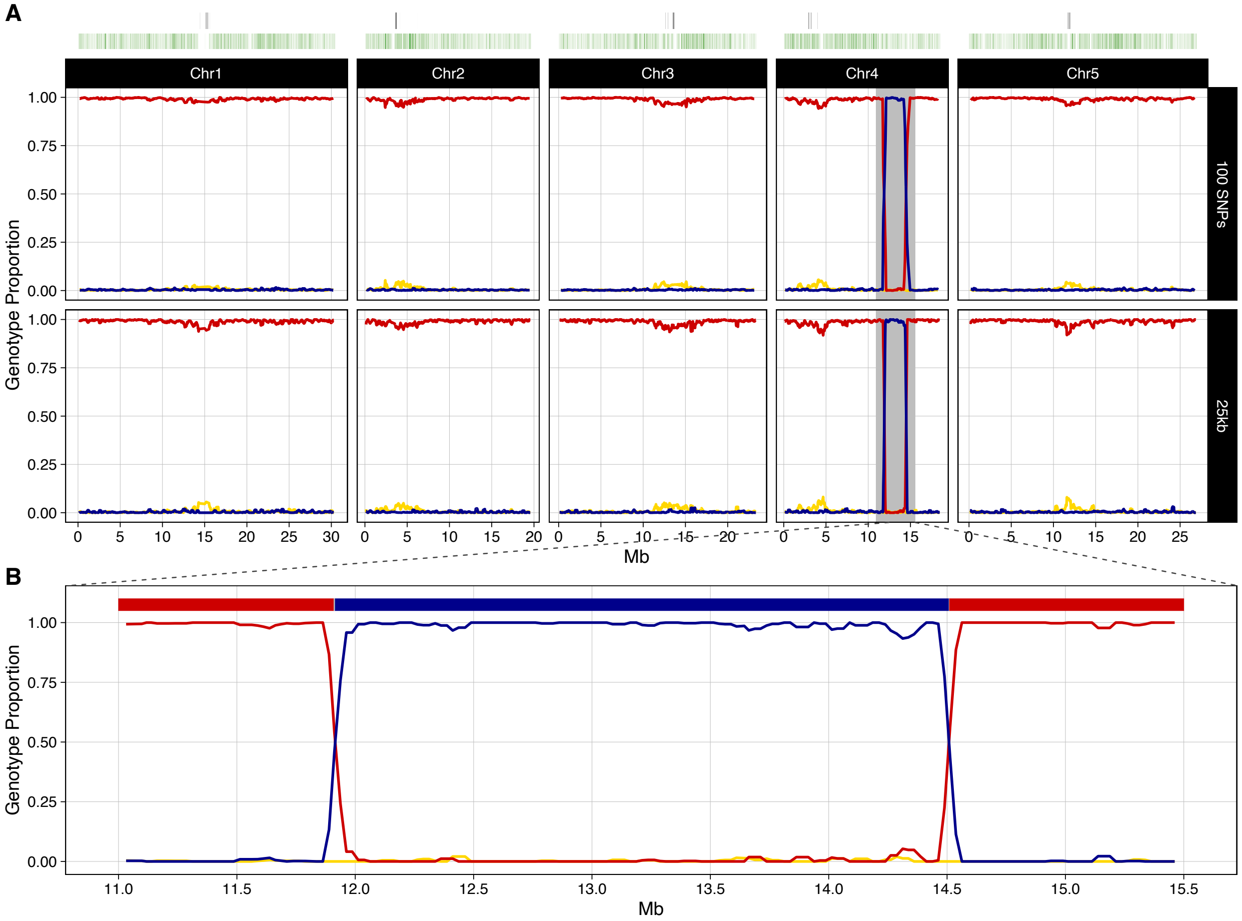
A
Chr1
Chr2
Chr3
Chr4
Chr5
100 SNPs
25kb
Genotype Proportion
Mb
B
Genotype Proportion
Mb

C_019

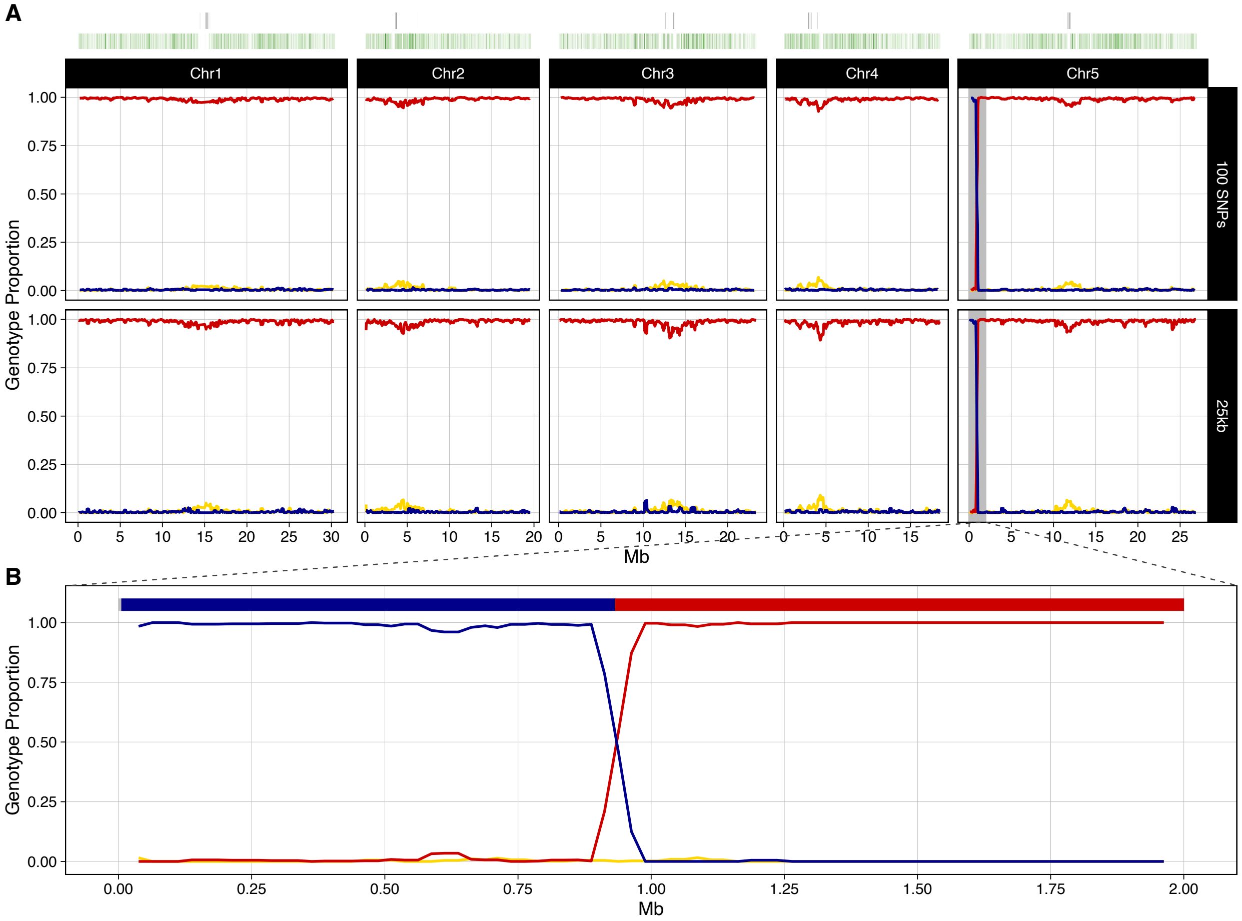
A
Chr1
Chr2
Chr3
Chr4
Chr5
100 SNPs
25kb
Genotype Proportion
Mb
B
Genotype Proportion
Mb

# C_021

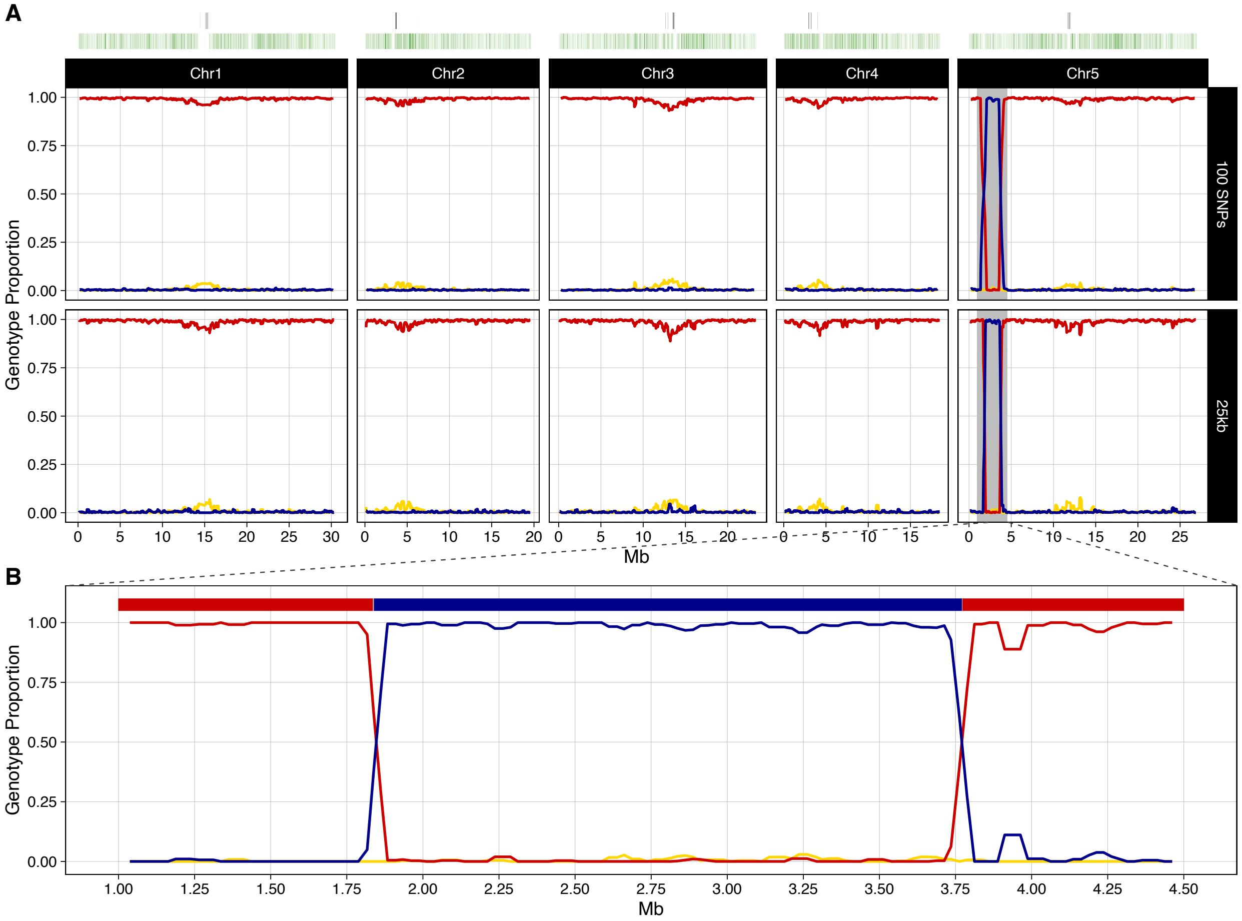
A
Chr1
Chr2
Chr3
Chr4
Chr5
100 SNPs
25kb
Genotype Proportion
1.00
0.75
0.50
0.25
0.00
Mb
B
Genotype Proportion
Mb

# C_023

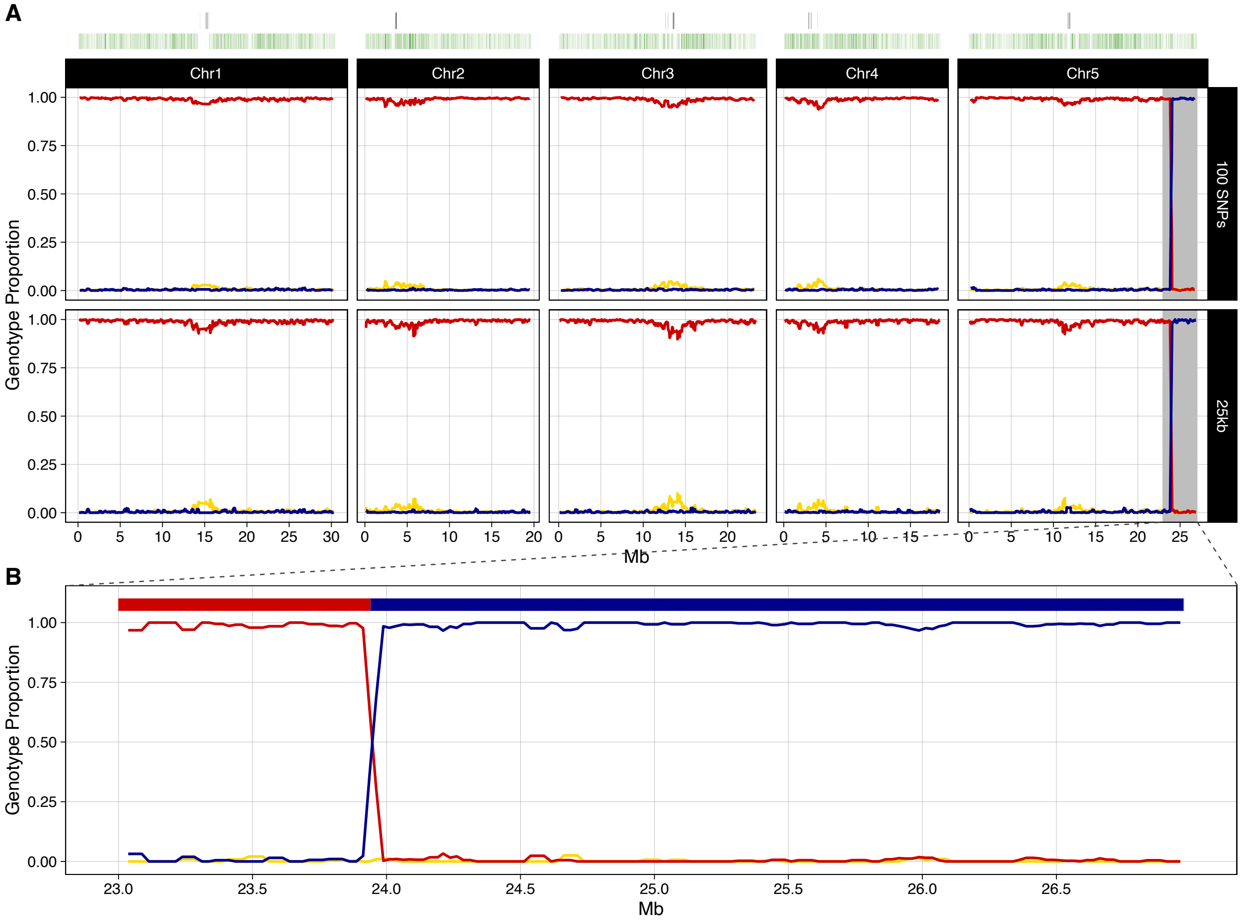
A
Chr1
Chr2
Chr3
Chr4
Chr5
100 SNPs
25kb
Genotype Proportion
1.00
0.75
0.50
0.25
0.00
0 5 10 15 20 25 30
0 5 10 15 20
0 5 10 15 20
0 5 10 15
0 5 10 15 20 25
Mb
B
Genotype Proportion
1.00
0.75
0.50
0.25
0.00
23.0 23.5 24.0 24.5 25.0 25.5 26.0 26.5
Mb

**C_026**

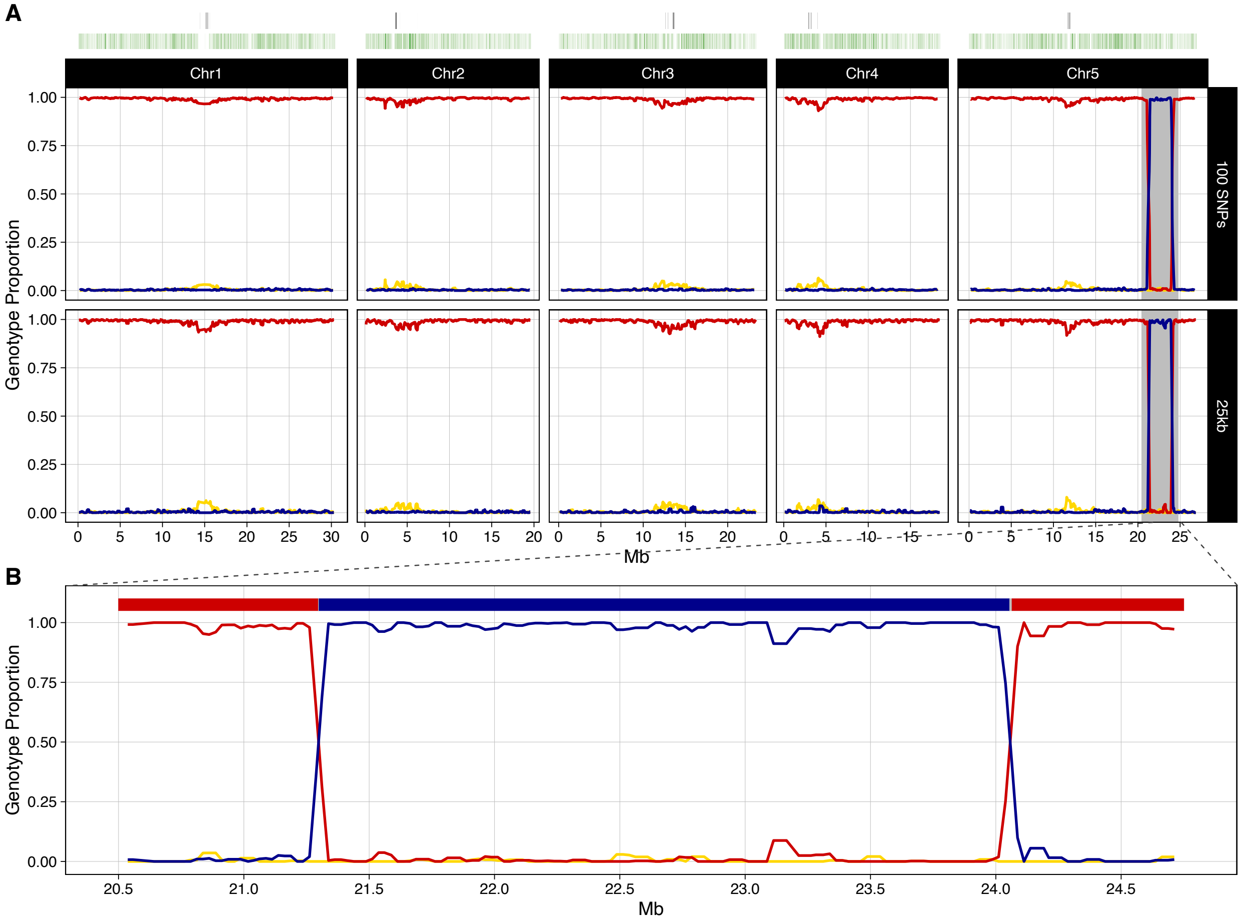
A
Chr1
Chr2
Chr3
Chr4
Chr5
100 SNPs
25kb
Genotype Proportion
1.00
0.75
0.50
0.25
0.00
Mb
B
Genotype Proportion
20.5
21.0
21.5
22.0
22.5
23.0
23.5
24.0
24.5
Mb

**C_031**

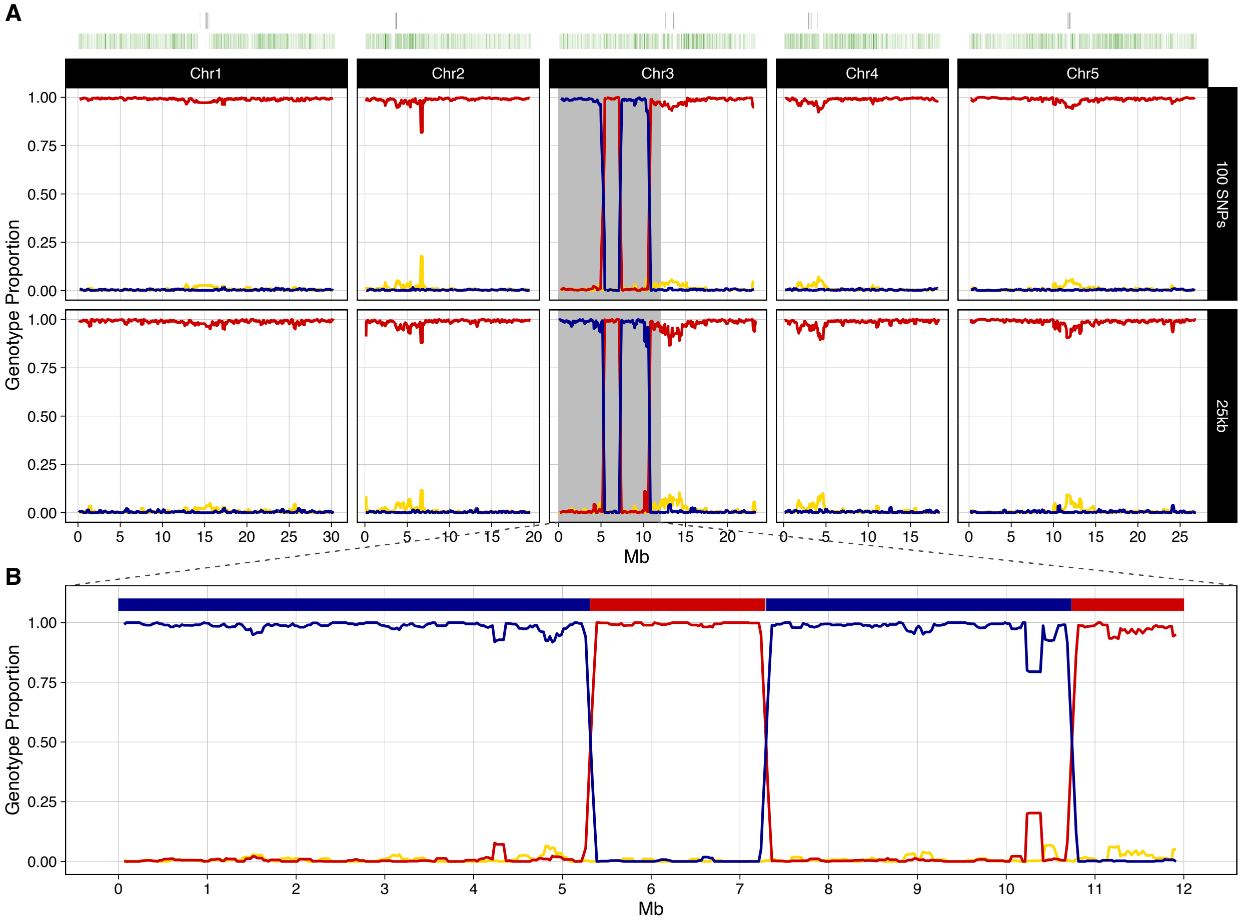
A
Chr1
Chr2
Chr3
Chr4
Chr5
100 SNPs
25kb
Genotype Proportion
1.00
0.75
0.50
0.25
0.00
0 5 10 15 20 25 30
Mb
B
0 1 2 3 4 5 6 7 8 9 10 11 12

**C_058**

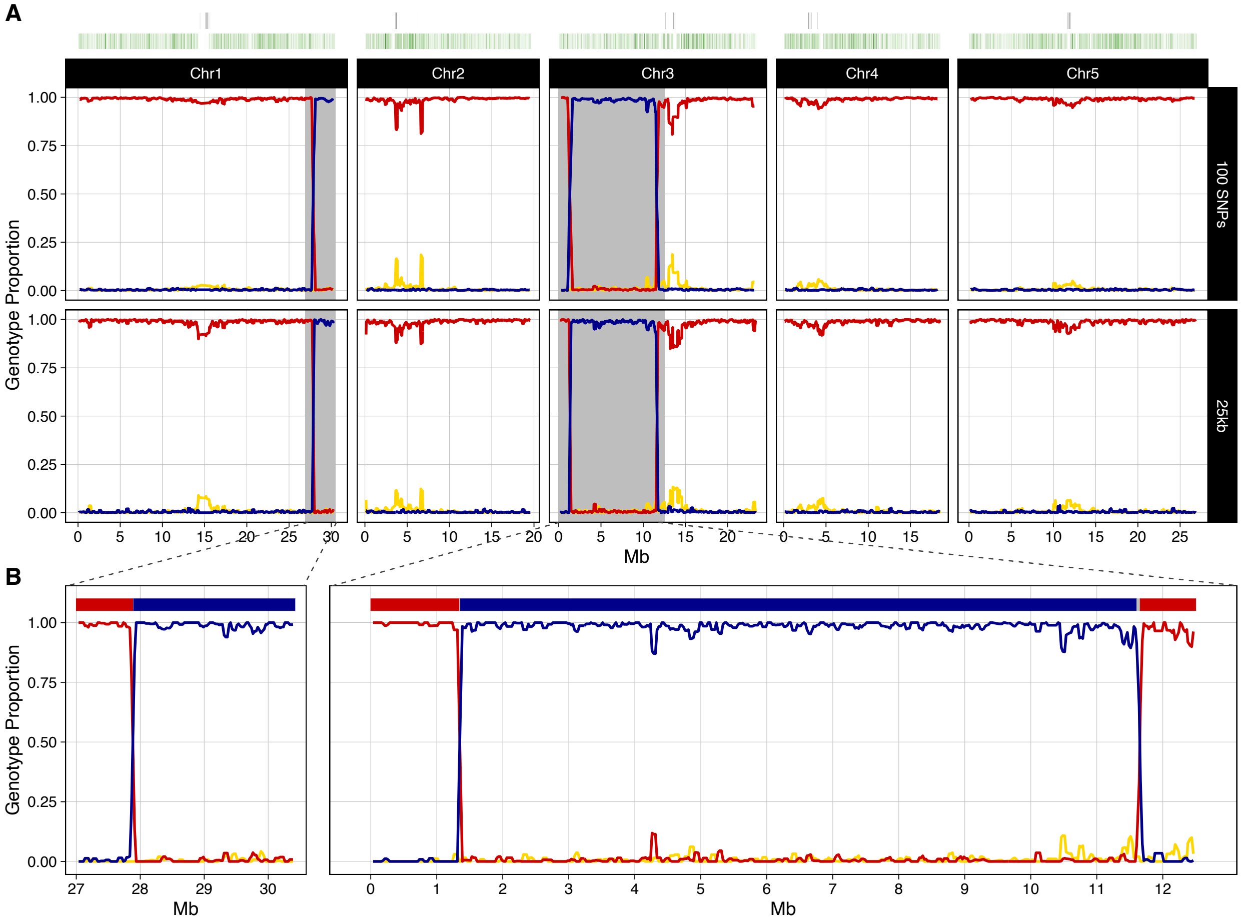

A
Chr1
Chr2
Chr3
Chr4
Chr5
100 SNPs
25kb
Genotype Proportion
1.00
0.75
0.50
0.25
0.00
0
5
10
15
20
25
30
Mb
B
27
28
29
30
1
2
3
4
6
7
8
9
11
12

**C_059**

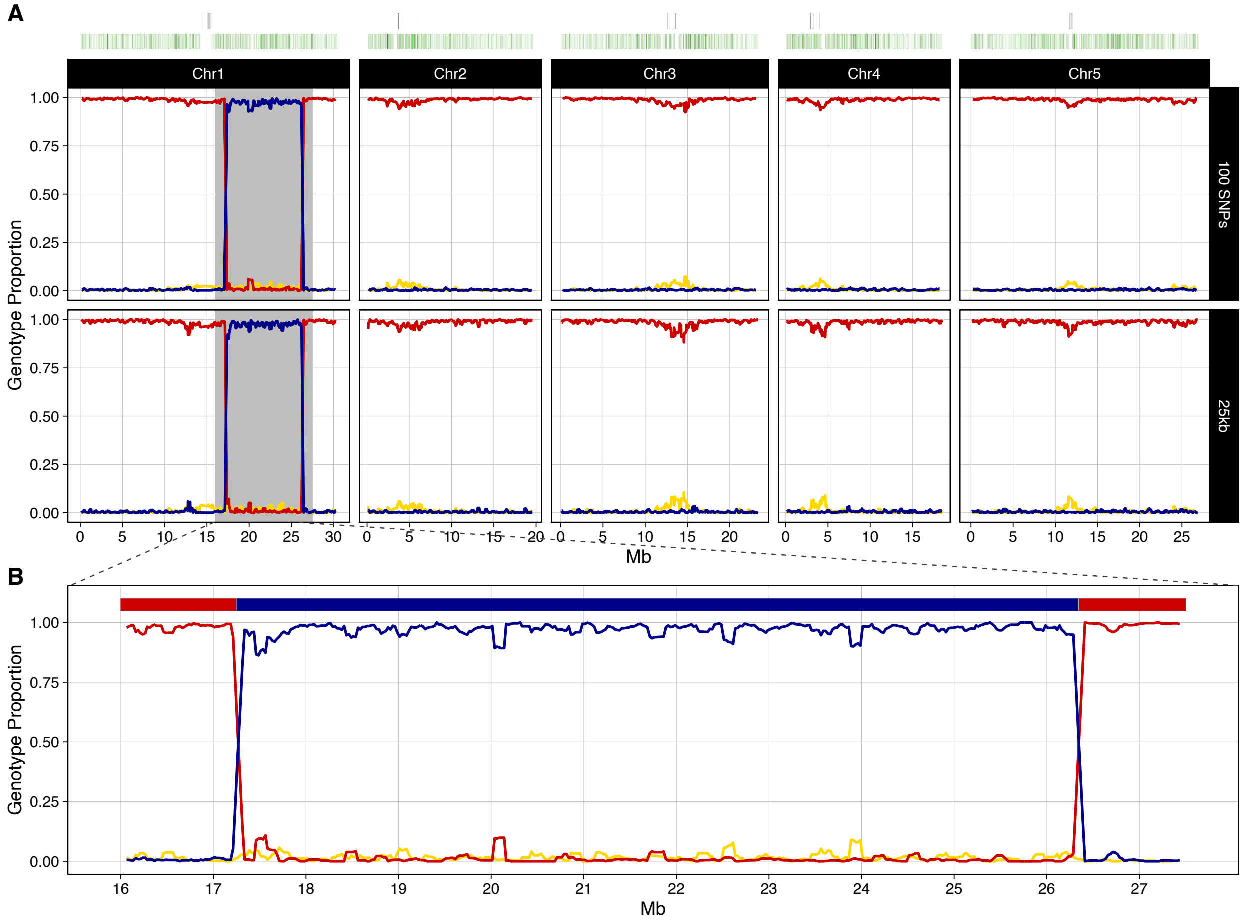
A
Chr1
Chr2
Chr3
Chr4
Chr5
100 SNPs
25kb
Genotype Proportion
Mb
B
Genotype Proportion
Mb

**C_086**

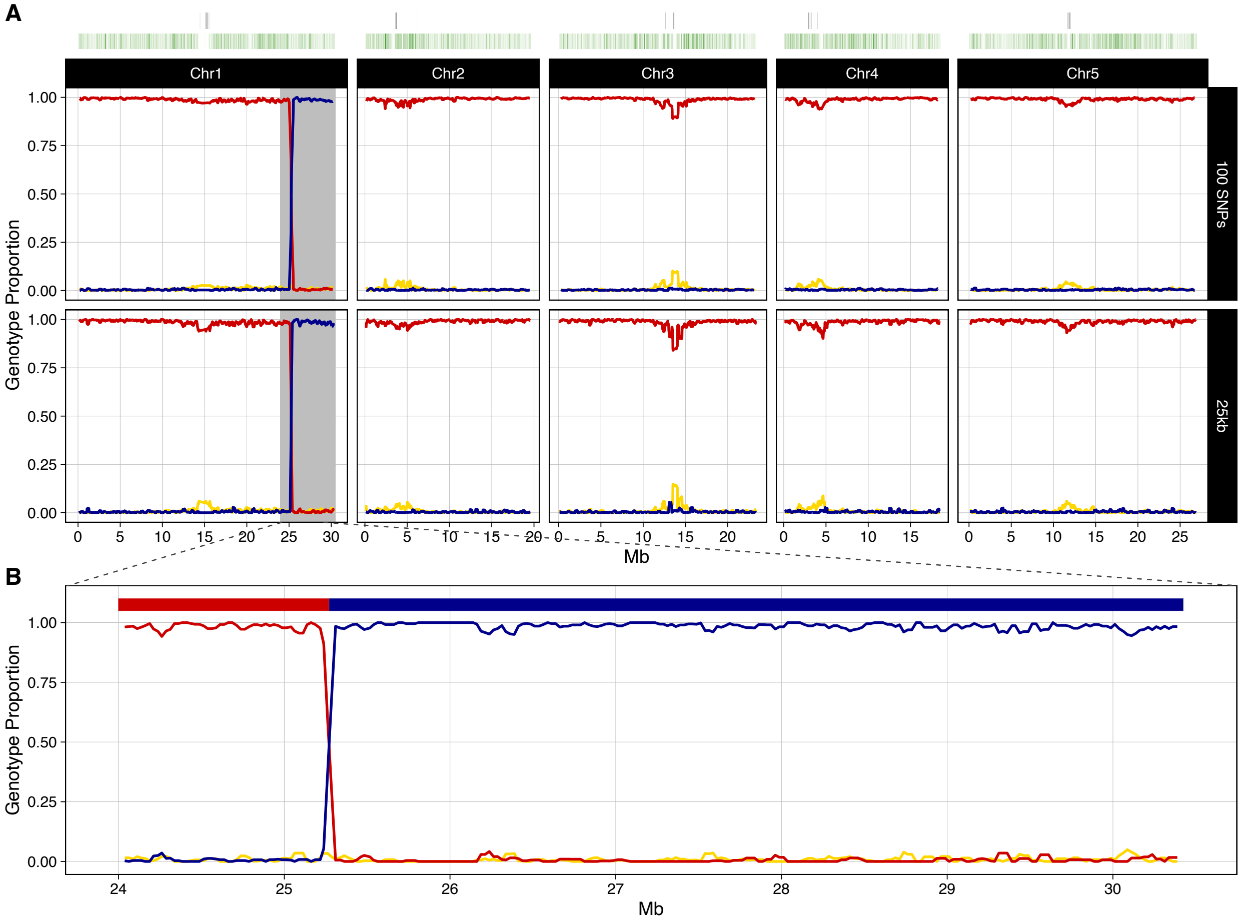
A
B
Chr1
Chr2
Chr3
Chr4
Chr5
100 SNPs
25kb
Genotype Proportion
Mb

**C_087**

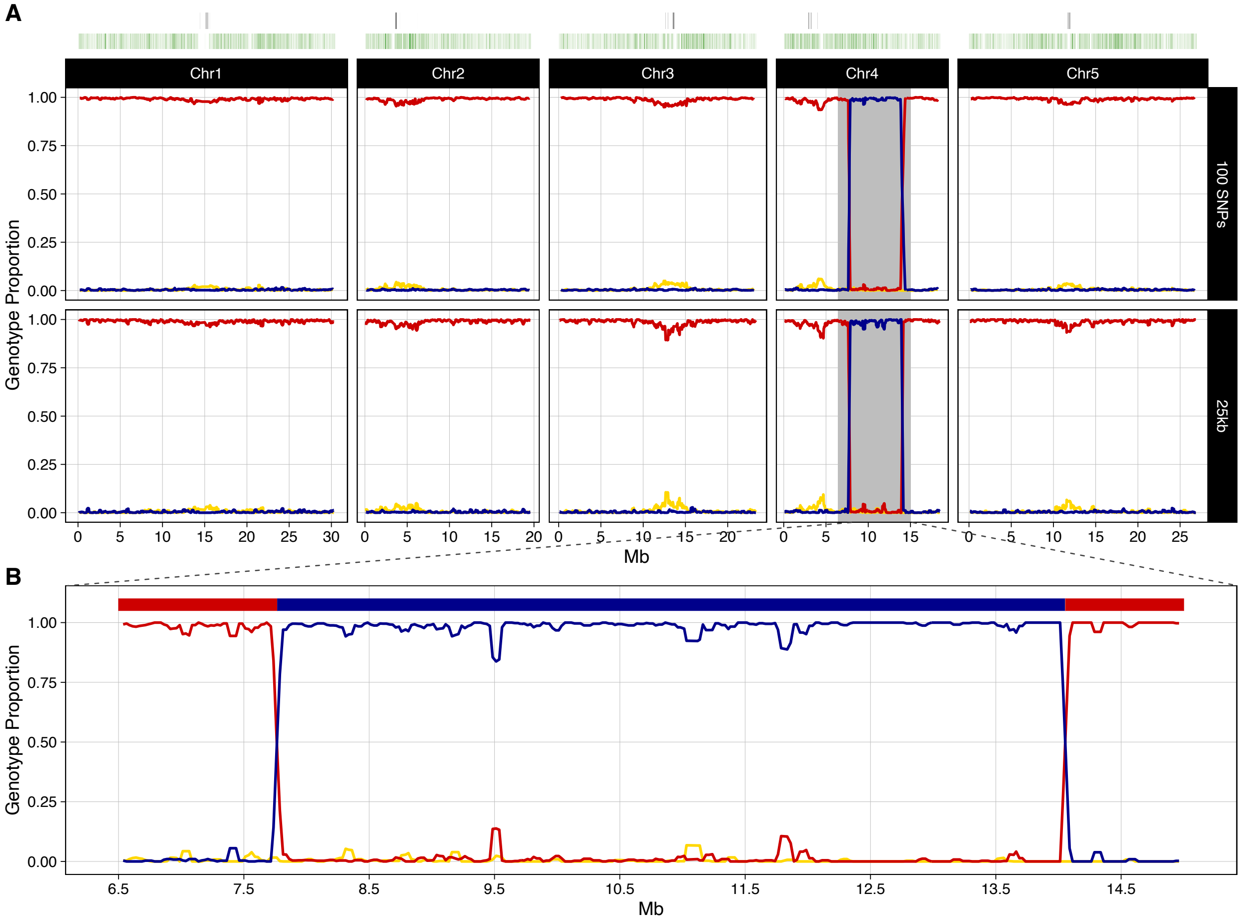
A
Chr1
Chr2
Chr3
Chr4
Chr5
100 SNPs
25kb
Genotype Proportion
1.00
0.75
0.50
0.25
0.00
Mb
B
6.5
7.5
8.5
9.5
10.5
11.5
12.5
13.5
14.5

**C_106**

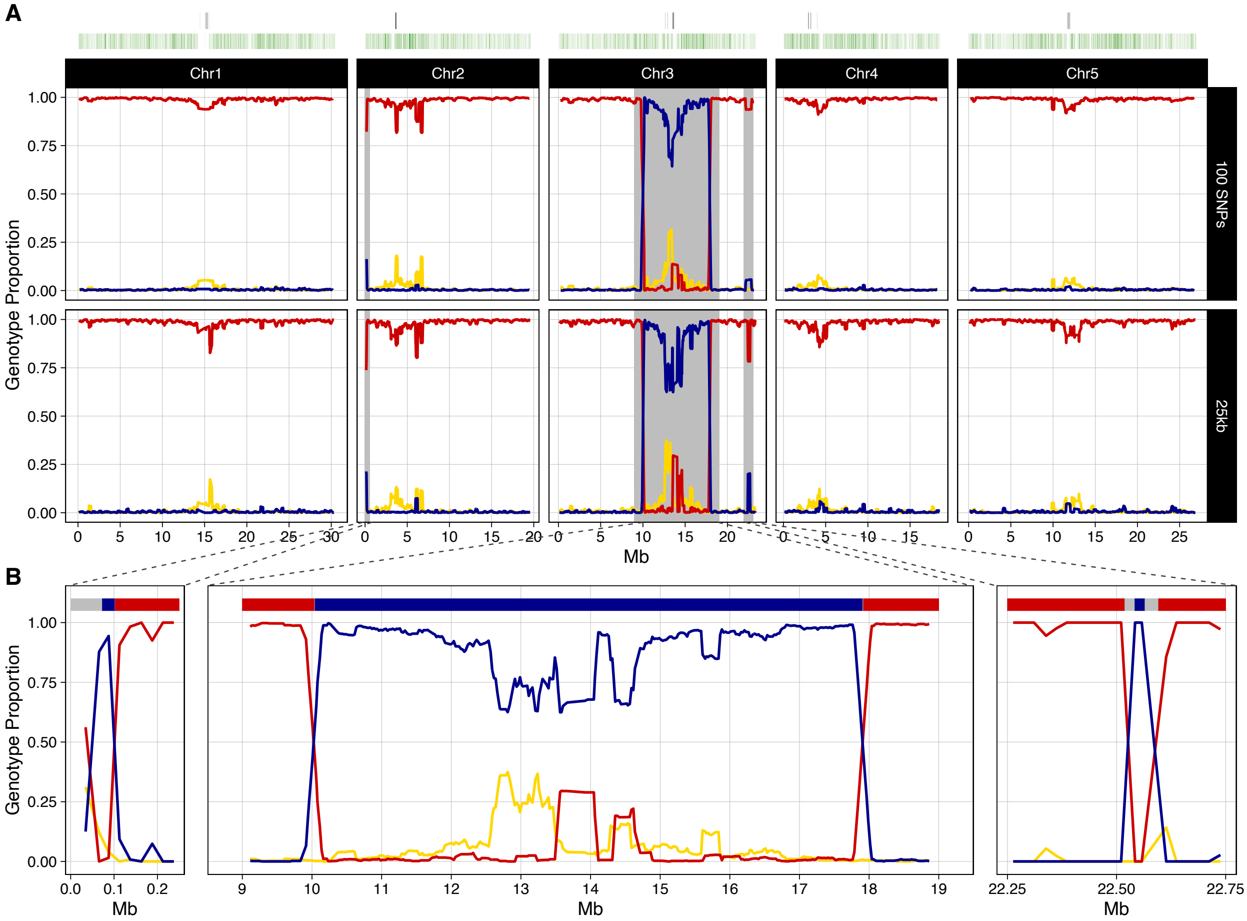
A
Chr1
Chr2
Chr3
Chr4
Chr5
100 SNPs
25kb
Genotype Proportion
1.00
0.75
0.50
0.25
0.00
Mb
B
Genotype Proportion
Mb
Mb
Mb

C_109

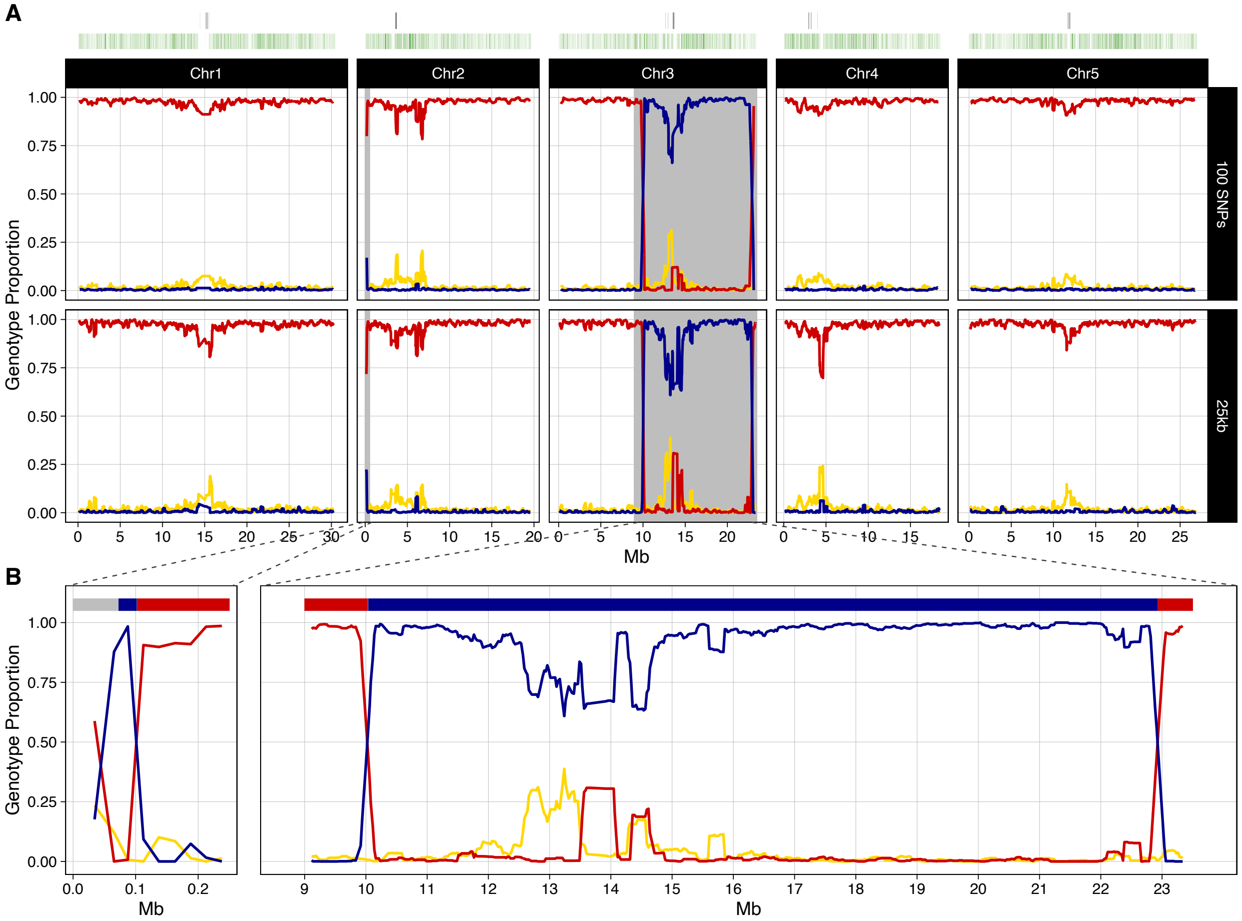

A
Chr1
Chr2
Chr3
Chr4
Chr5
100 SNPs
25kb
Genotype Proportion
1.00
0.75
0.50
0.25
0.00
Mb
B
Mb

**C_110**

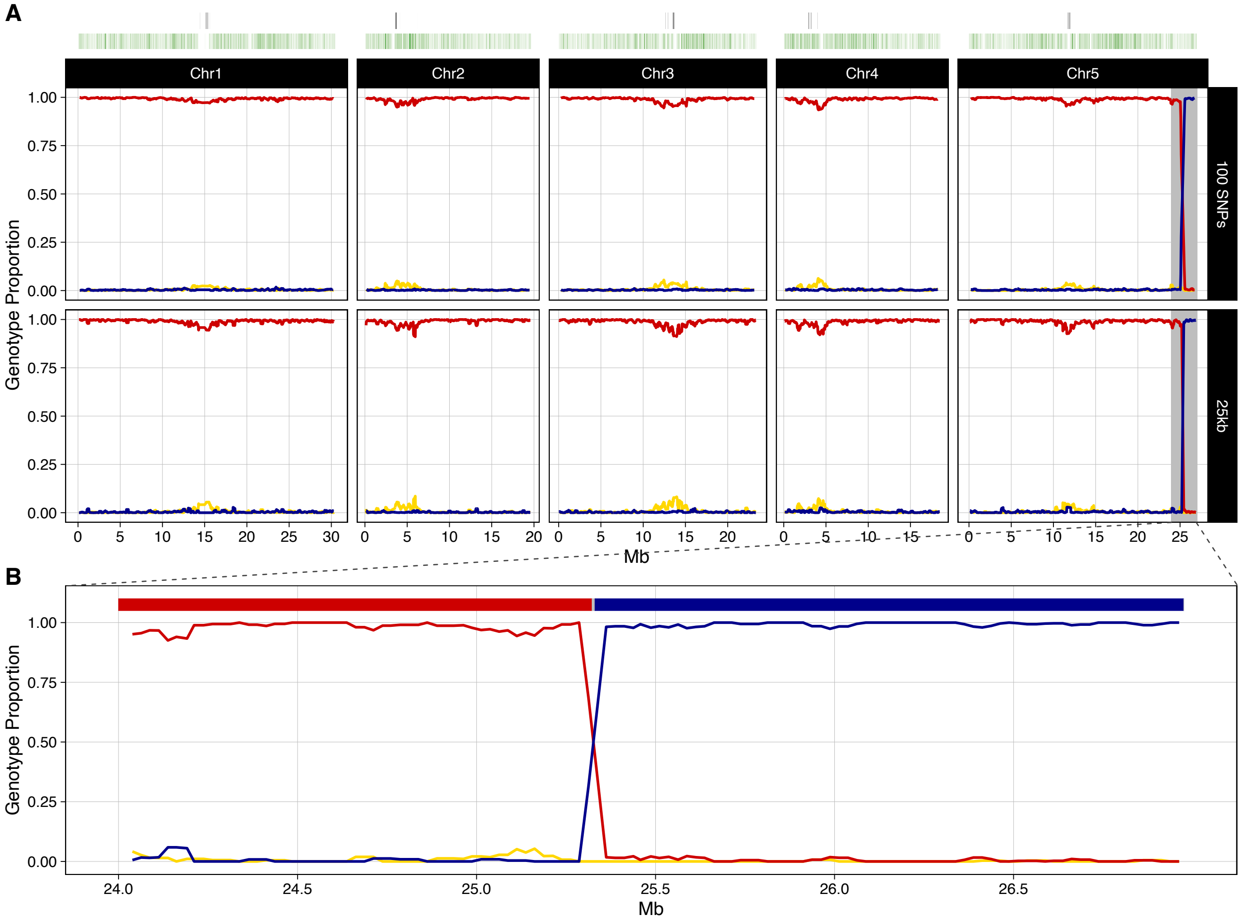
A
Chr1
Chr2
Chr3
Chr4
Chr5
100 SNPs
25kb
Genotype Proportion
1.00
0.75
0.50
0.25
0.00
Mb
B
24.0
24.5
25.0
25.5
26.0
26.5

**C_111**

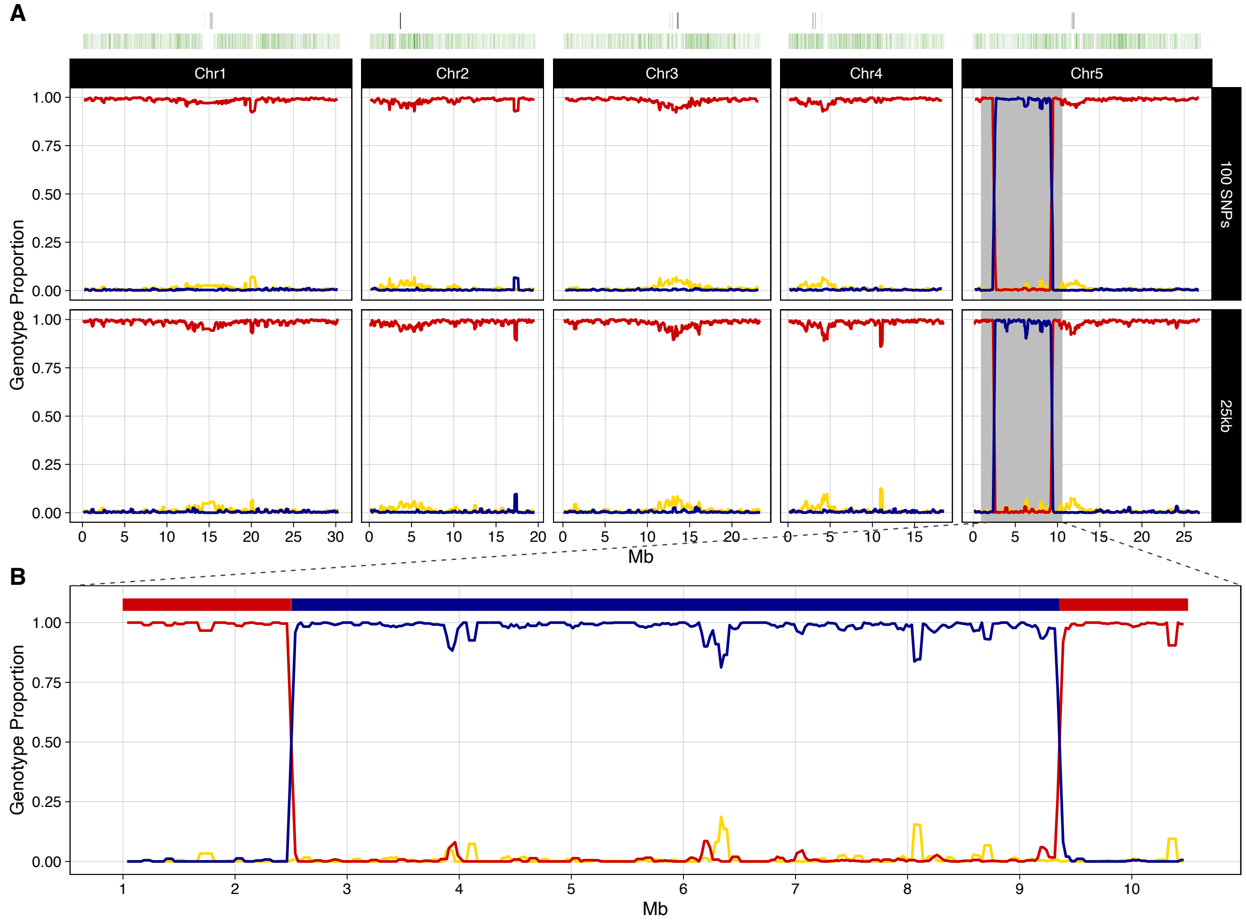
A
Chr1
Chr2
Chr3
Chr4
Chr5
100 SNPs
25kb
Genotype Proportion
1.00
0.75
0.50
0.25
0.00
0
5
10
15
20
25
30
Mb
B
Genotype Proportion
1
2
3
4
5
6
7
8
9
10
Mb

**C_115**

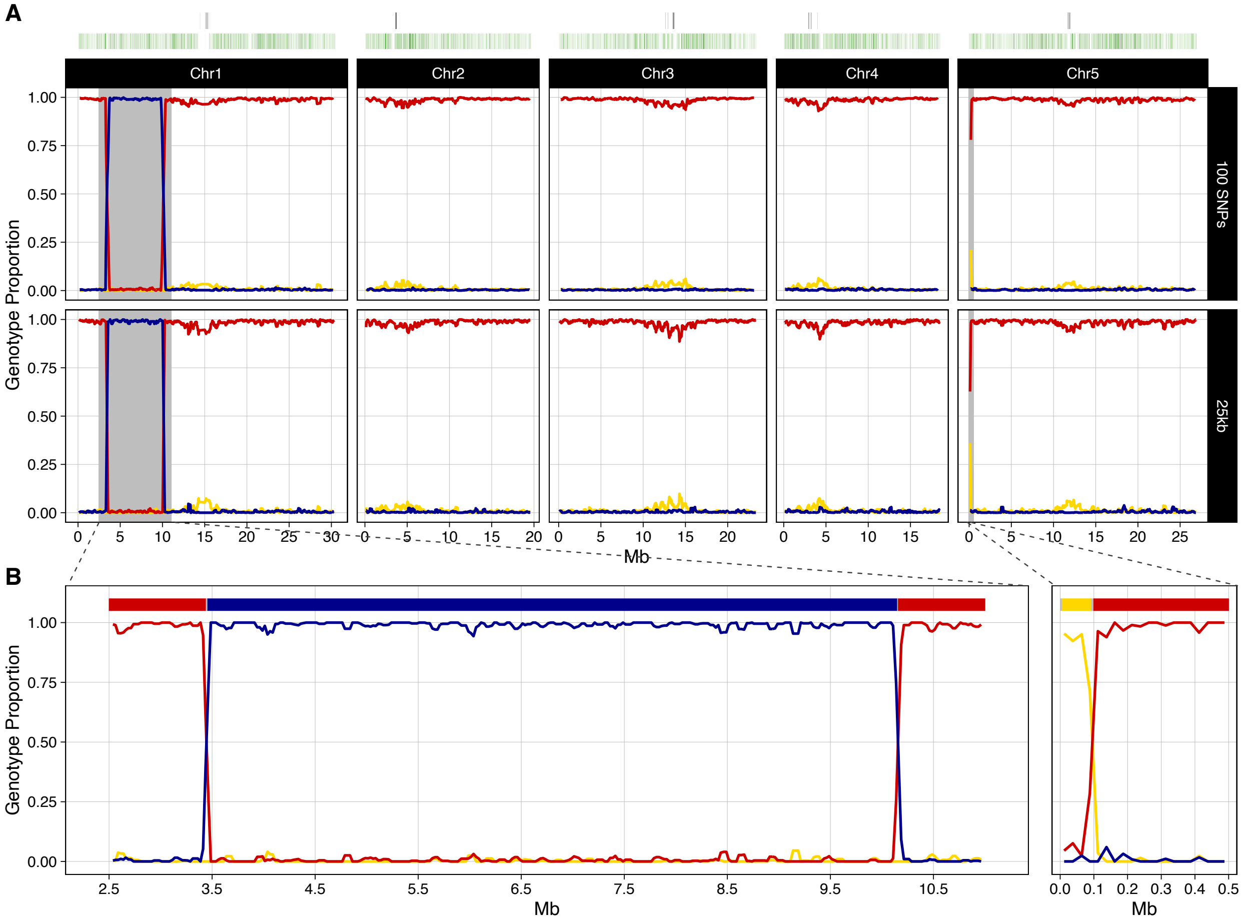
A
Chr1
Chr2
Chr3
Chr4
Chr5
100 SNPs
25kb
Genotype Proportion
1.00
0.75
0.50
0.25
0.00
Mb
B
2.5
3.5
4.5
5.5
6.5
7.5
8.5
9.5
10.5
0.0 0.1 0.2 0.3 0.4 0.5

**C_117**

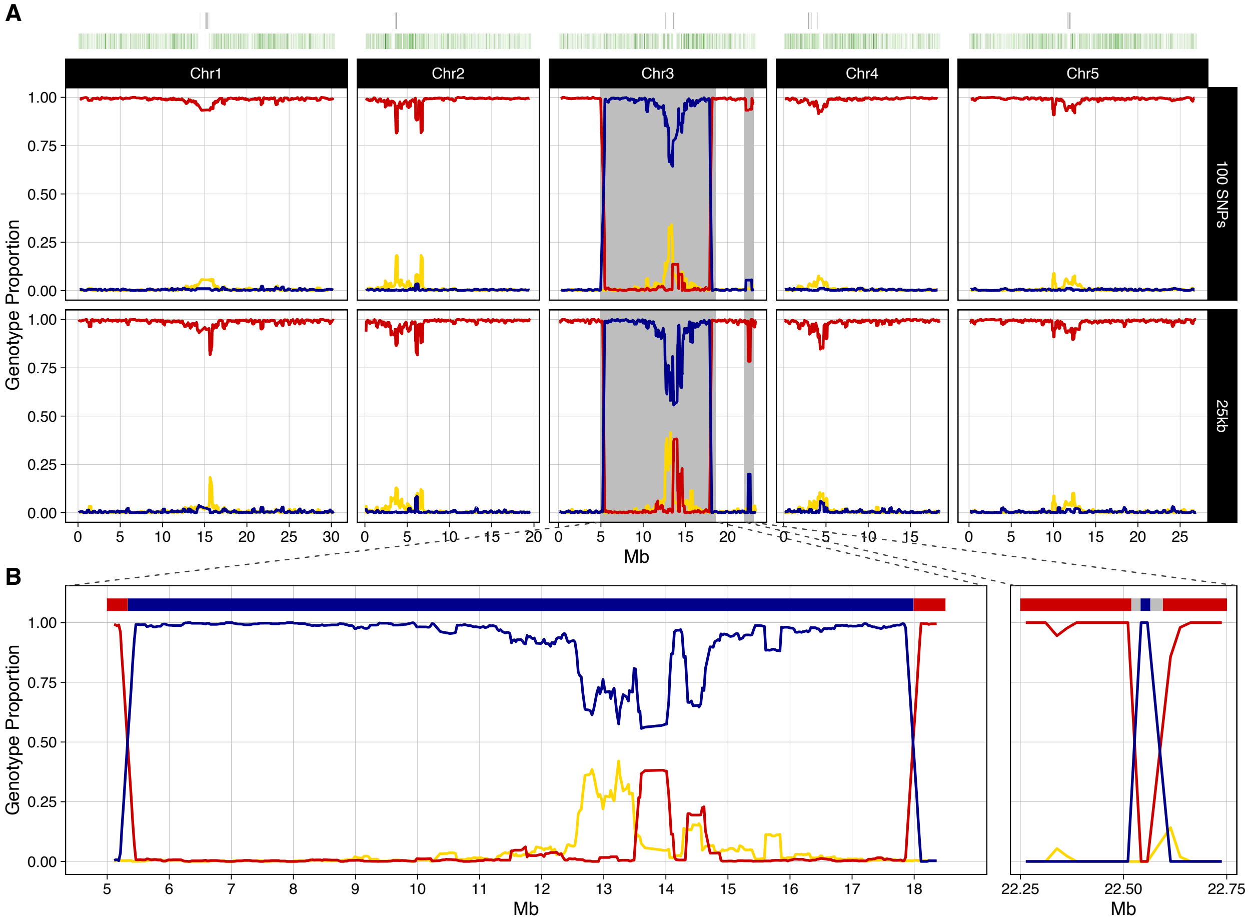
A
Chr1
Chr2
Chr3
Chr4
Chr5
100 SNPs
25kb
Genotype Proportion
Mb
B
Genotype Proportion
Mb
Mb

**C_118**

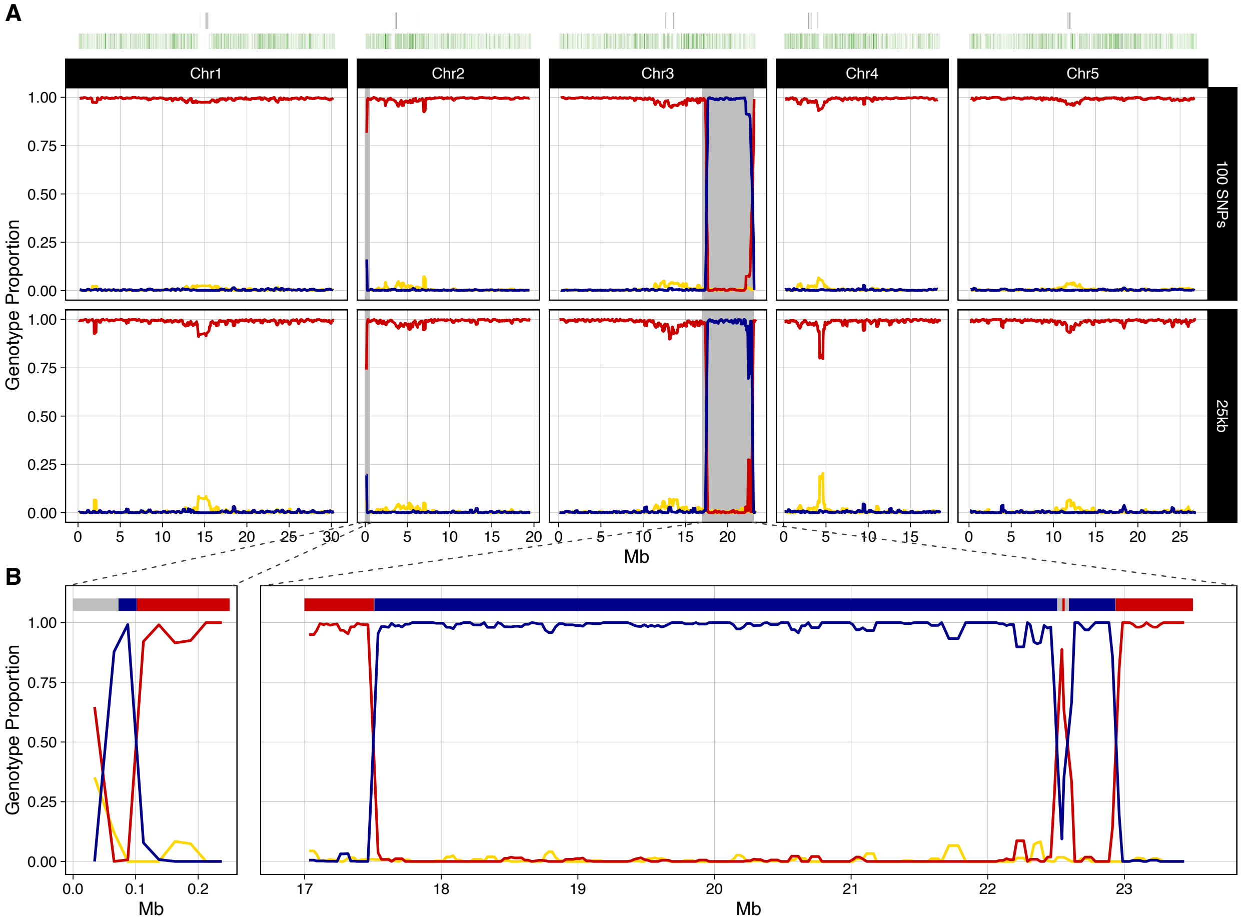
A
Chr1
Chr2
Chr3
Chr4
Chr5
100 SNPs
25kb
Genotype Proportion
1.00
0.75
0.50
0.25
0.00
Mb
B
Genotype Proportion
0.0
0.1
0.2
Mb
17
18
19
20
21
22
23
Mb

**C_148**

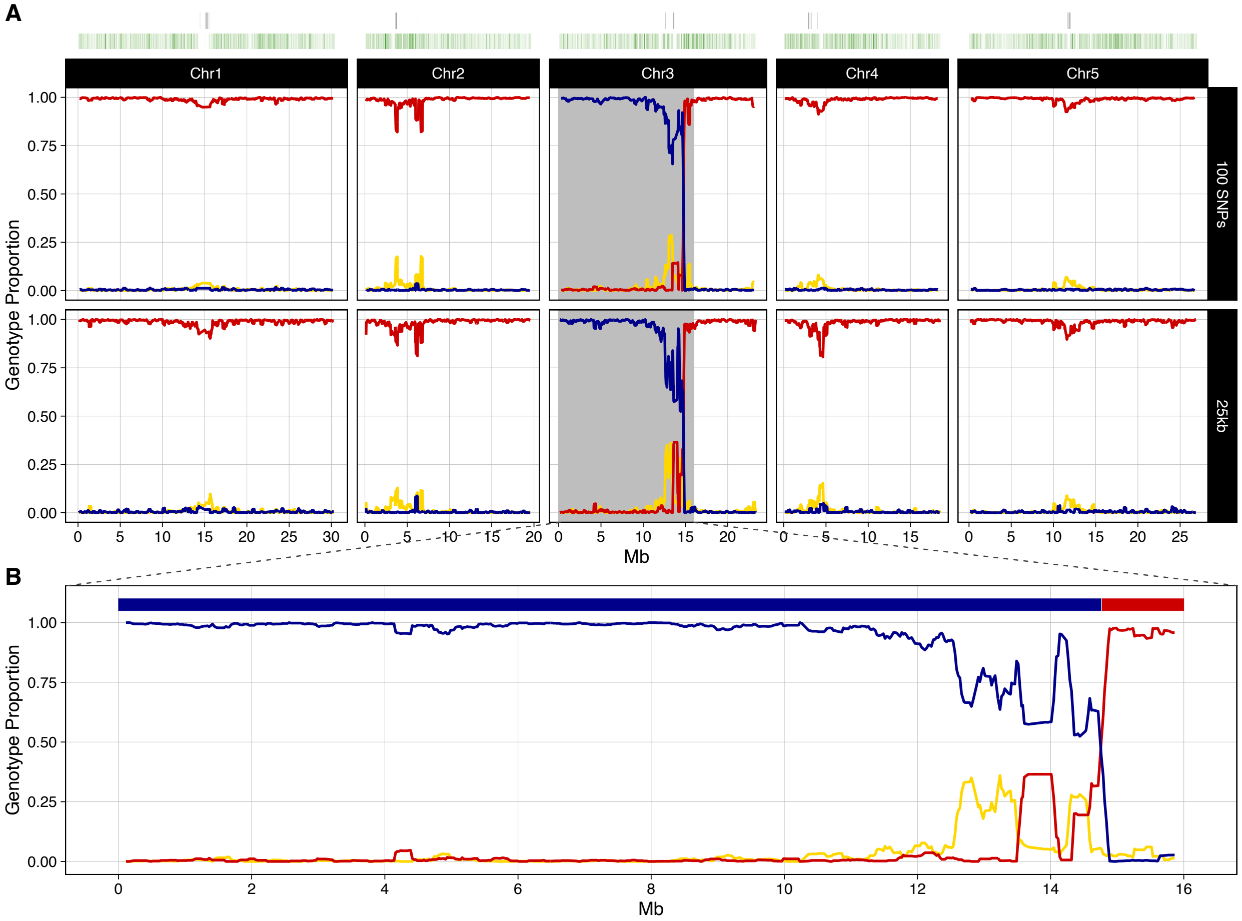
A
Chr1
Chr2
Chr3
Chr4
Chr5
100 SNPs
25kb
Genotype Proportion
1.00
0.75
0.50
0.25
0.00
0 5 10 15 20 25 30
0 5 10 15 20
0 5 10 15 20
0 5 10 15
0 5 10 15 20 25
Mb
B
Genotype Proportion
1.00
0.75
0.50
0.25
0.00
0 2 4 6 8 10 12 14 16
Mb

**C_149**

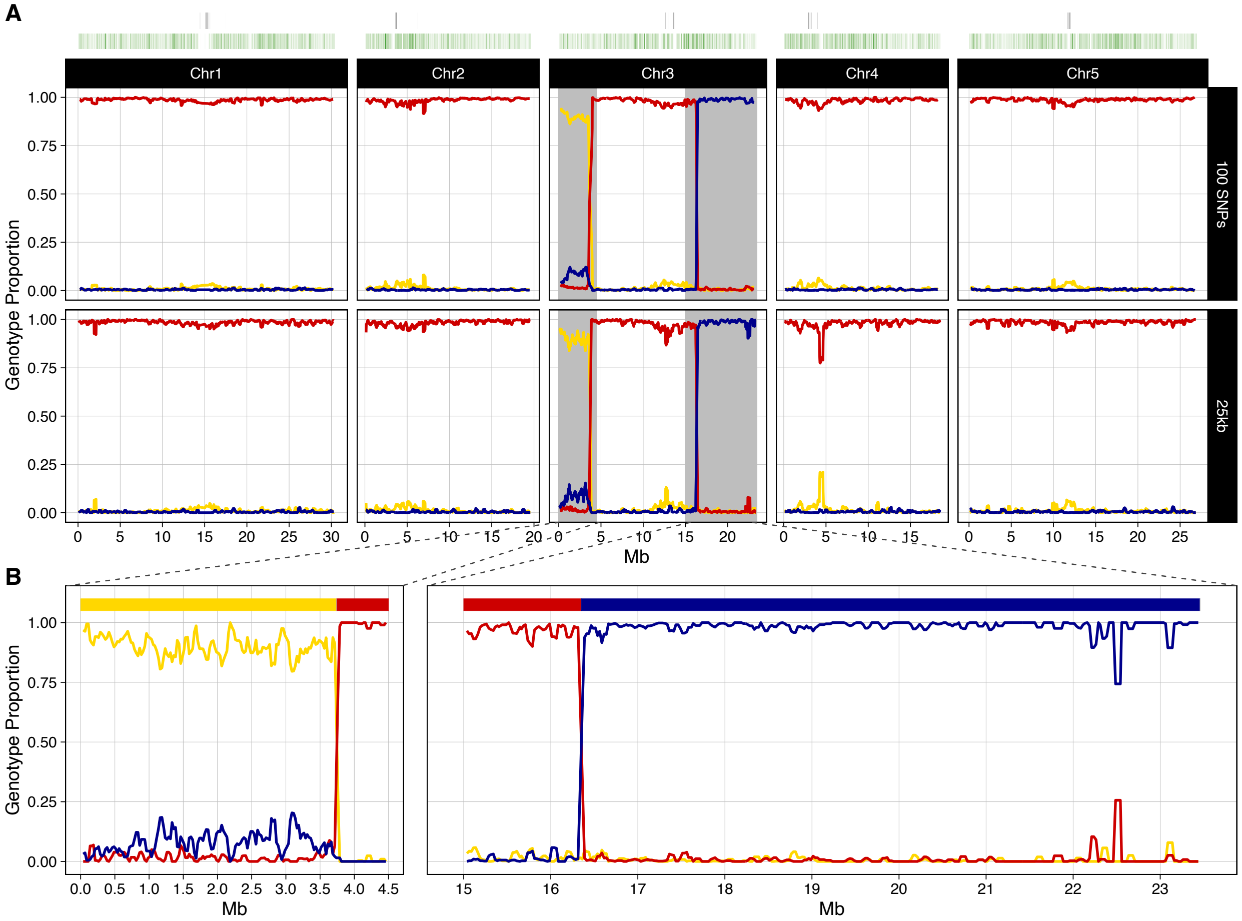
A
Chr1
Chr2
Chr3
Chr4
Chr5
100 SNPs
25kb
Genotype Proportion
1.00
0.75
0.50
0.25
0.00
0 5 10 15 20 25 30
Mb
B
Genotype Proportion
0.0 0.5 1.0 1.5 2.0 2.5 3.0 3.5 4.0 4.5
Mb
15 16 17 18 19 20 21 22 23
Mb

**C_152**

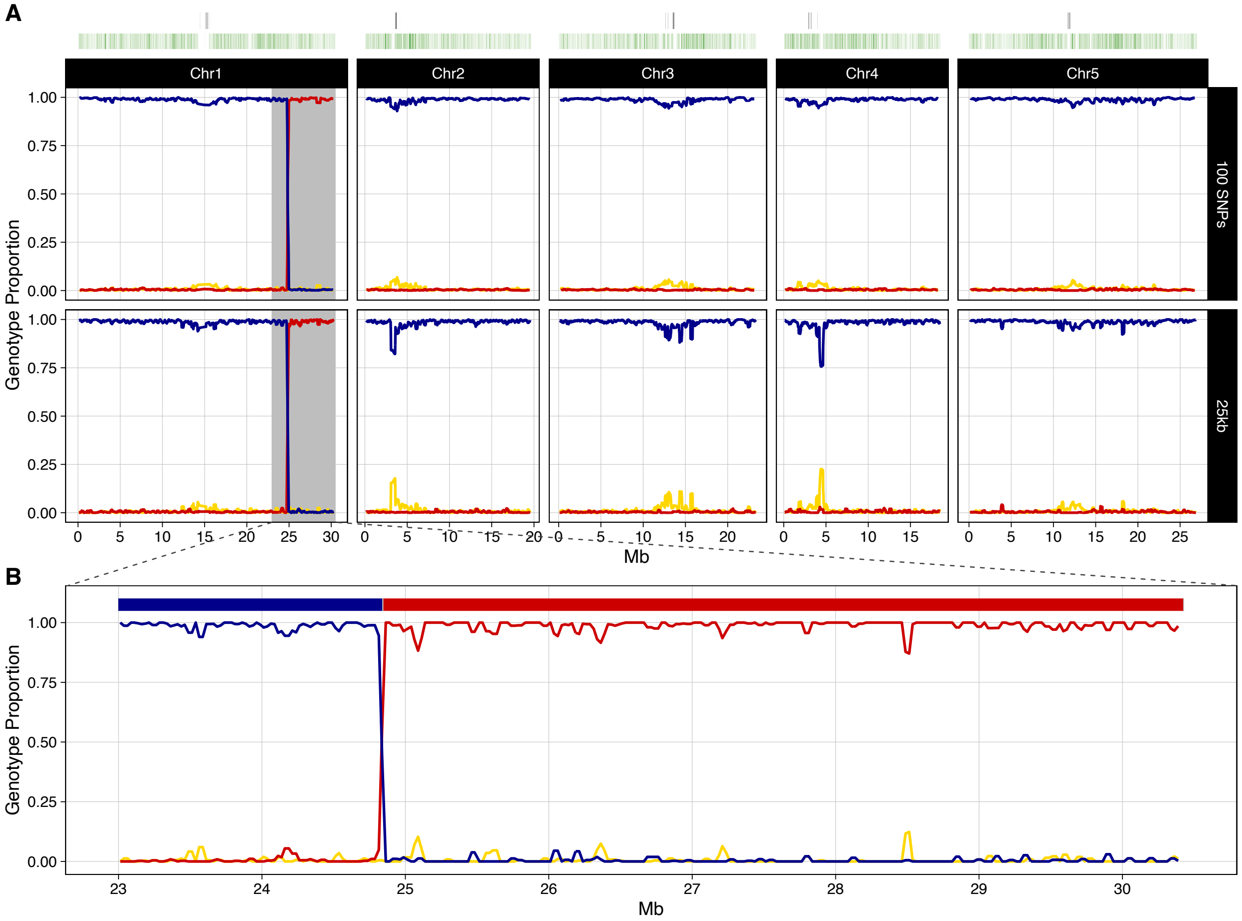
A
Chr1
Chr2
Chr3
Chr4
Chr5
100 SNPs
25kb
Genotype Proportion
1.00
0.75
0.50
0.25
0.00
Mb
B
Genotype Proportion
Mb
23
24
25
26
27
28
29
30

**R_008**

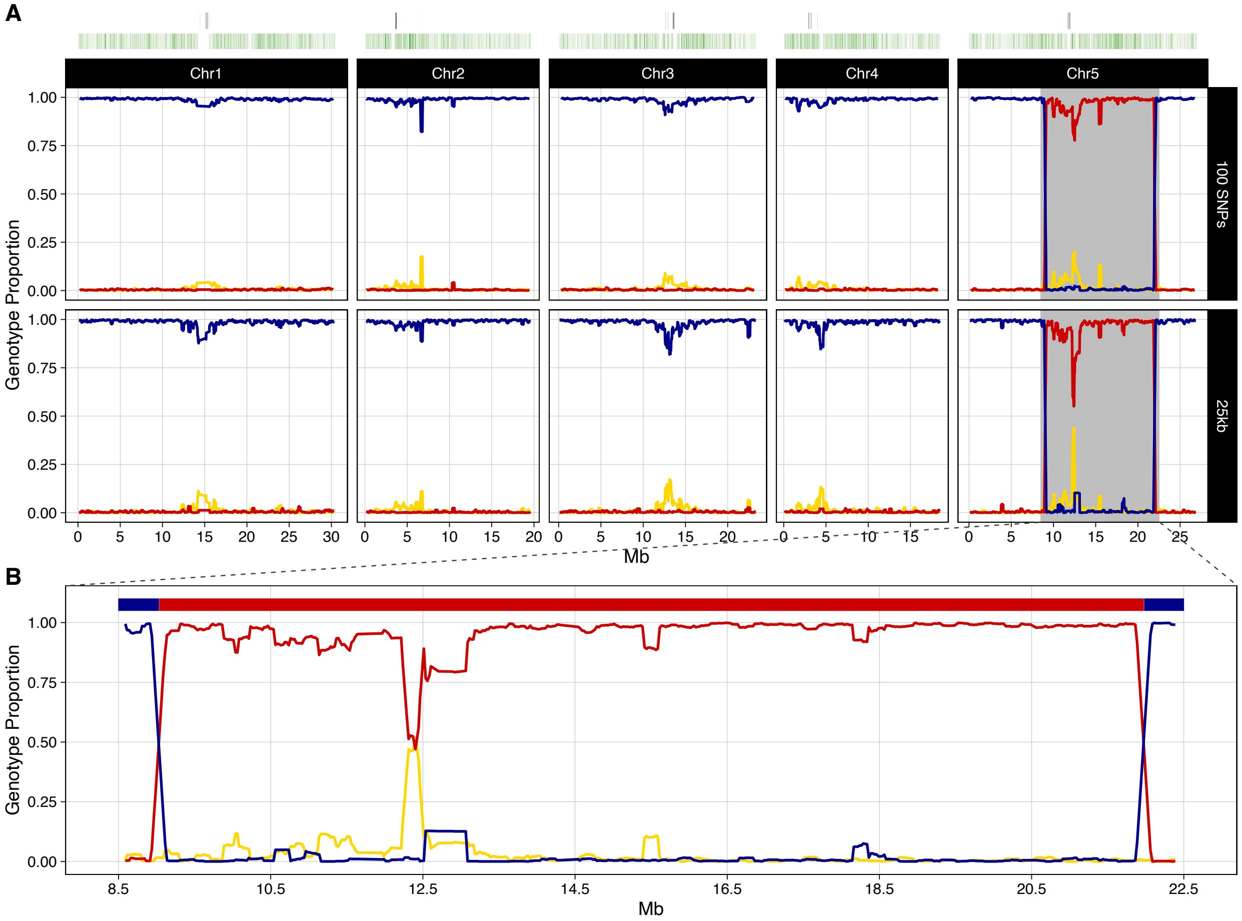

A
Chr1
Chr2
Chr3
Chr4
Chr5
100 SNPs
25kb
Genotype Proportion
1.00
0.75
0.50
0.25
0.00
0 5 10 15 20 25 30
Mb
B
8.5 10.5 12.5 14.5 16.5 18.5 20.5 22.5

**R_011**

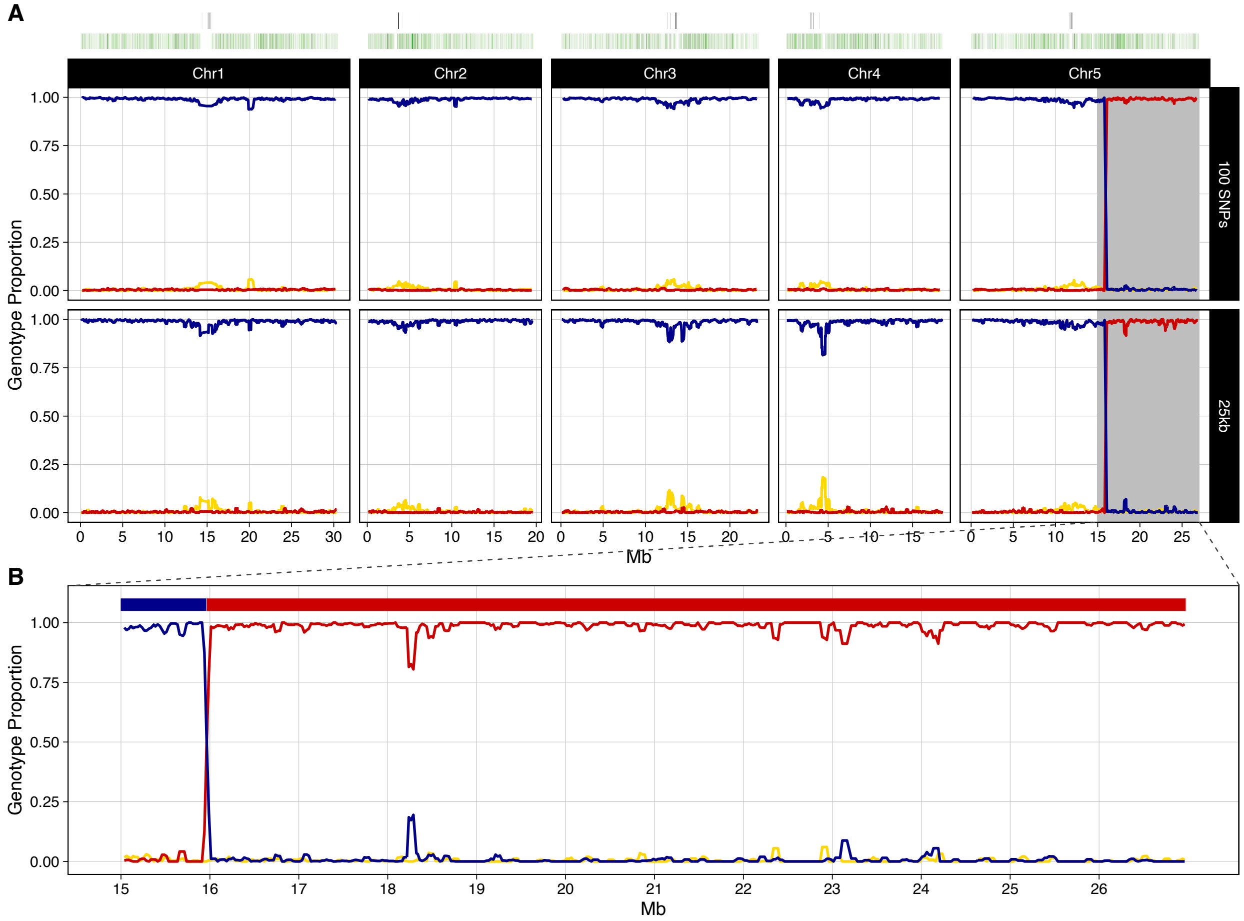
A
Chr1
Chr2
Chr3
Chr4
Chr5
100 SNPs
25kb
Genotype Proportion
1.00
0.75
0.50
0.25
0.00
0 5 10 15 20 25 30
0 5 10 15 20
0 5 10 15 20
0 5 10 15
0 5 10 15 20 25
Mb
B
Genotype Proportion
15 16 17 18 19 20 21 22 23 24 25 26
Mb

**R_013**

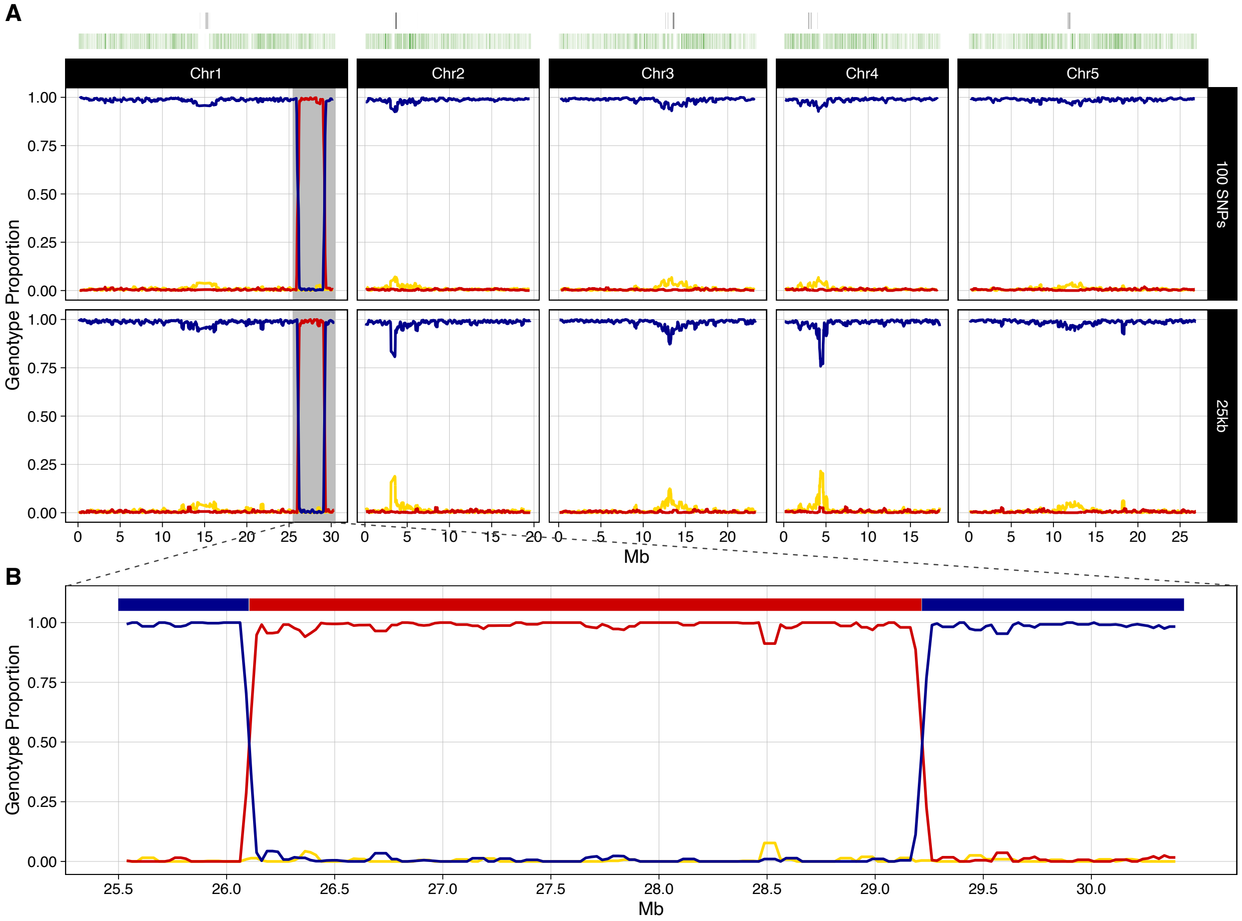
A
Chr1
Chr2
Chr3
Chr4
Chr5
100 SNPs
25kb
Genotype Proportion
1.00
0.75
0.50
0.25
0.00
0 5 10 15 20 25 30
Mb
B
25.5 26.0 26.5 27.0 27.5 28.0 28.5 29.0 29.5 30.0

**R_035**

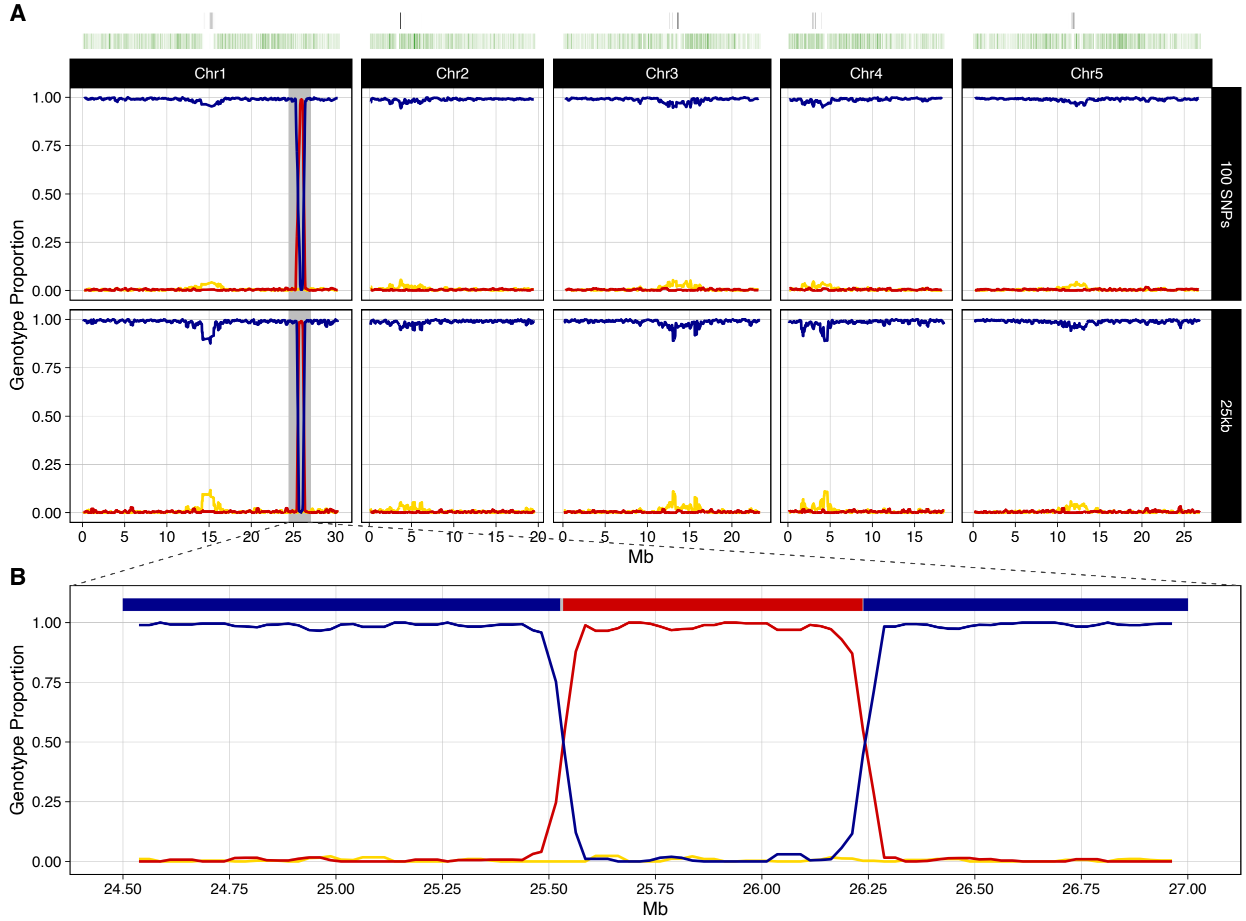
A
Chr1
Chr2
Chr3
Chr4
Chr5
100 SNPs
25kb
Genotype Proportion
1.00
0.75
0.50
0.25
0.00
Mb
B
24.50
24.75
25.00
25.25
25.50
25.75
26.00
26.25
26.50
26.75
27.00

**R_036**

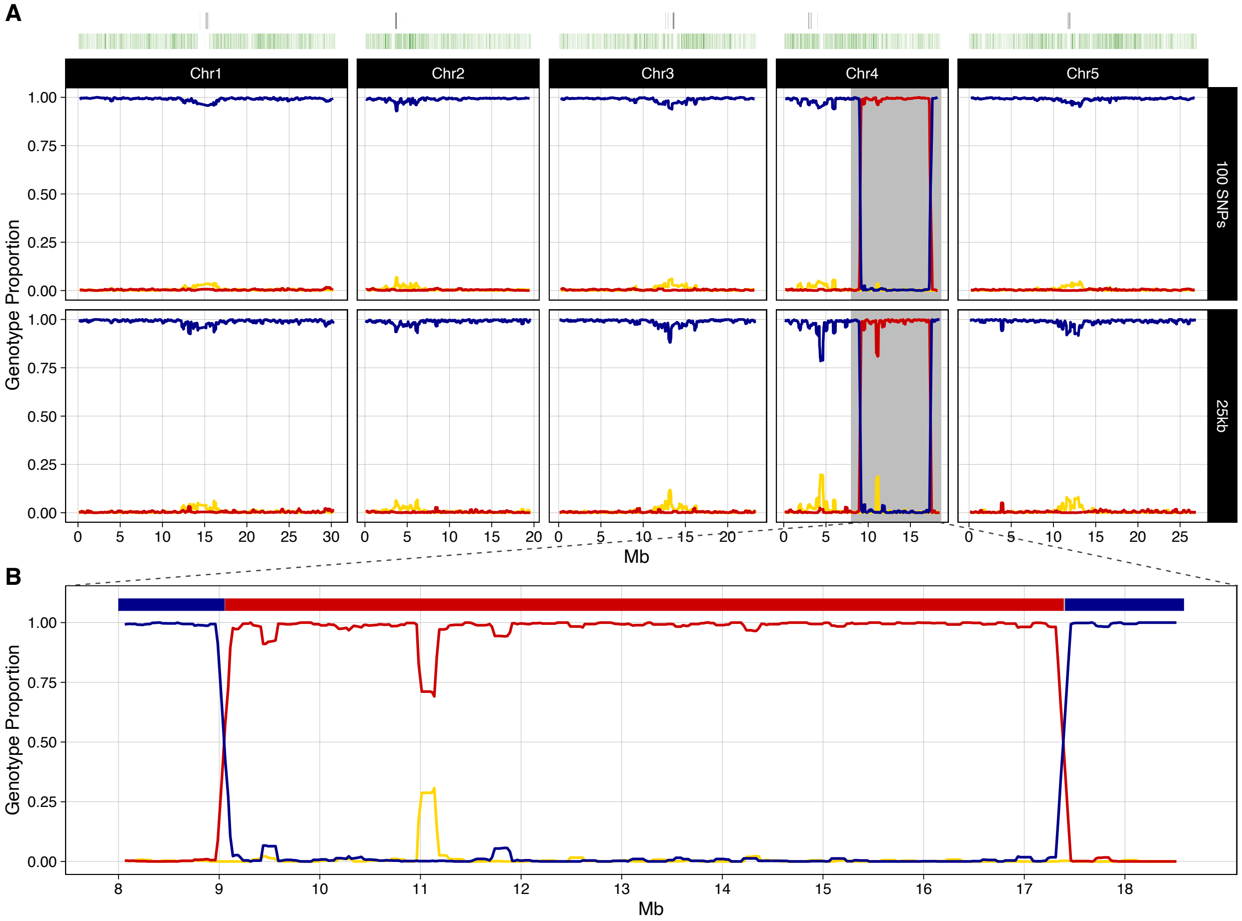
A
Chr1
Chr2
Chr3
Chr4
Chr5
100 SNPs
25kb
Genotype Proportion
1.00
0.75
0.50
0.25
0.00
Mb
B
Genotype Proportion
Mb

R_038

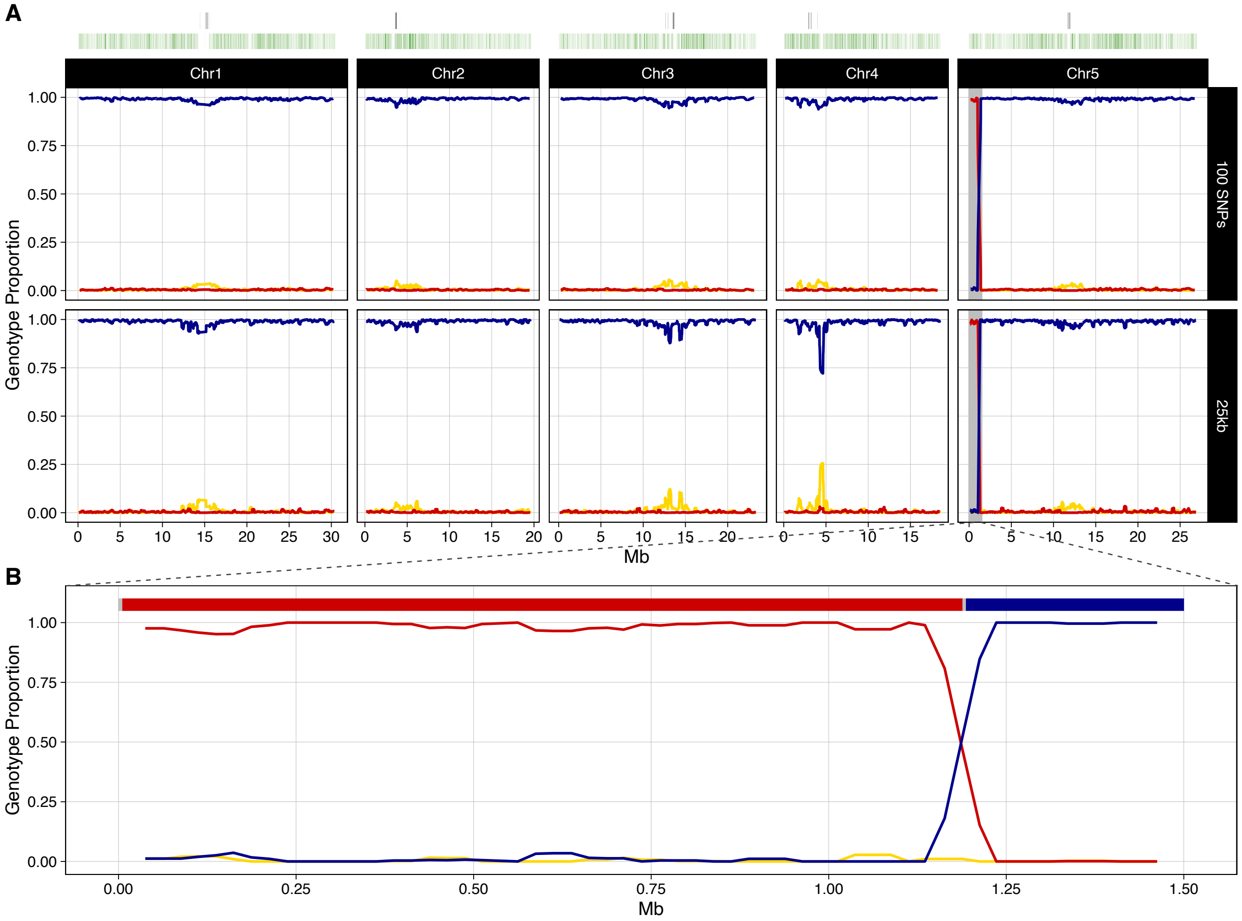
A
Chr1
Chr2
Chr3
Chr4
Chr5
100 SNPs
25kb
Genotype Proportion
1.00
0.75
0.50
0.25
0.00
Mb
B
Genotype Proportion
Mb

**R_040**

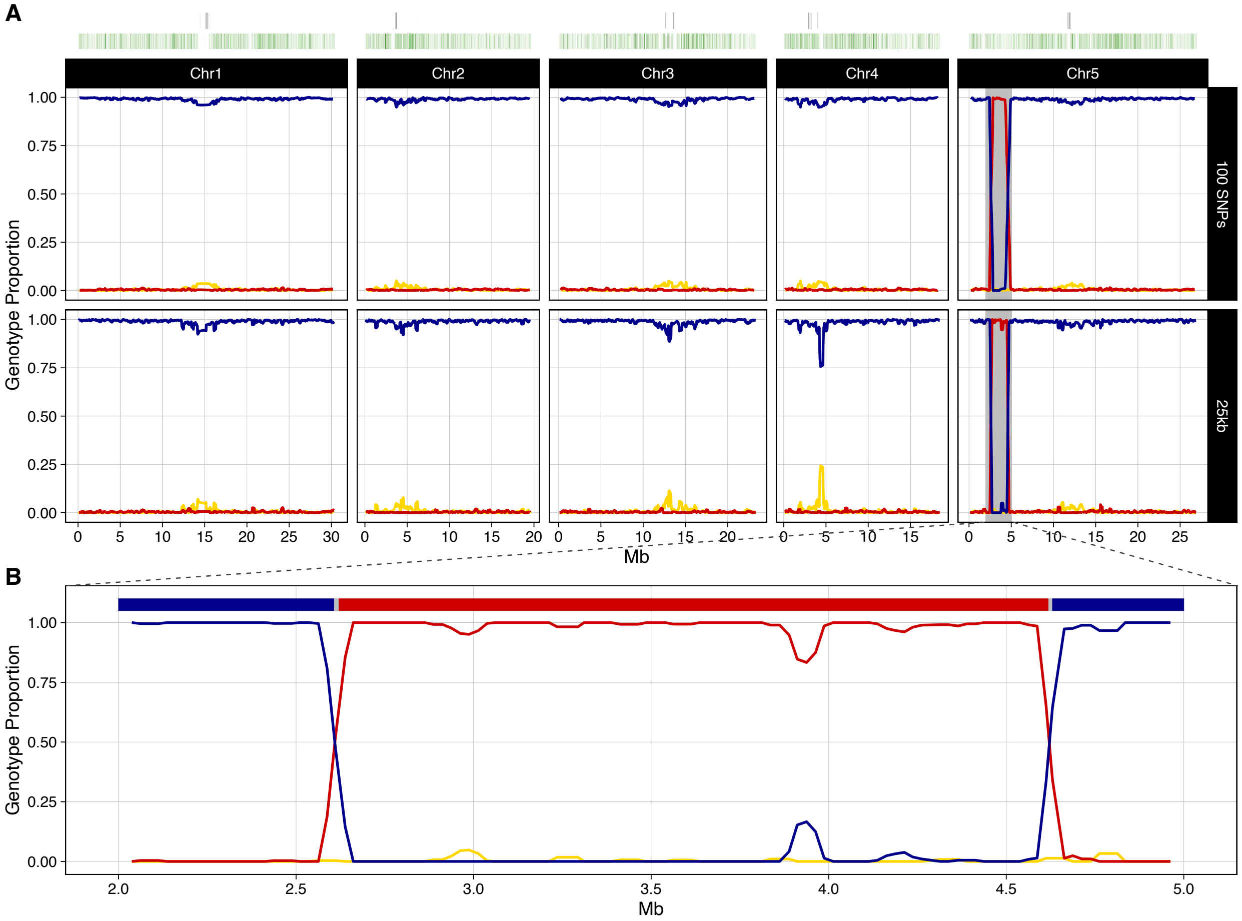
A
Chr1
Chr2
Chr3
Chr4
Chr5
100 SNPs
25kb
Genotype Proportion
1.00
0.75
0.50
0.25
0.00
0 5 10 15 20 25 30
0 5 10 15 20
0 5 10 15 20
0 5 10 15
0 5 10 15 20 25
Mb
B
Genotype Proportion
1.00
0.75
0.50
0.25
0.00
2.0 2.5 3.0 3.5 4.0 4.5 5.0
Mb

**R_041**

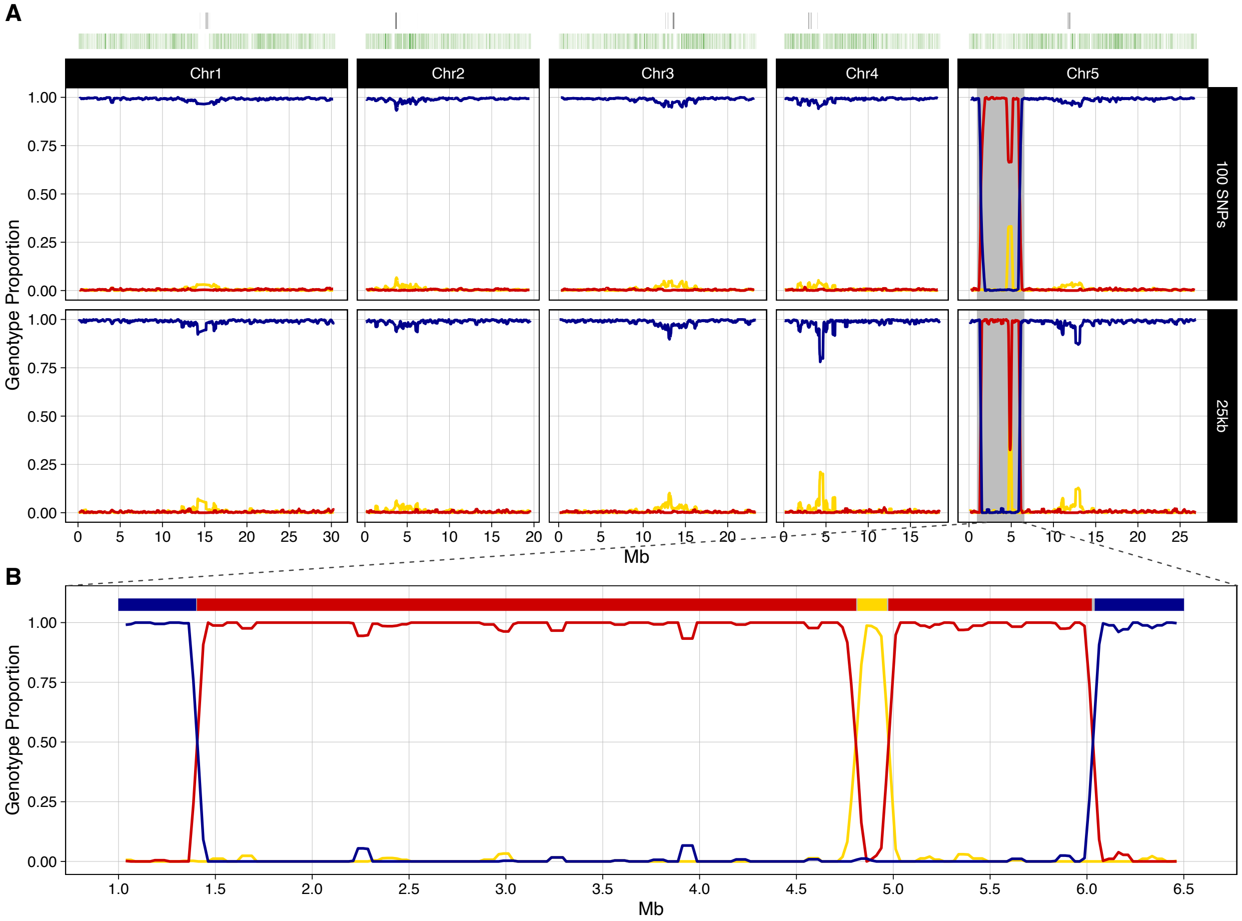
A
Chr1
Chr2
Chr3
Chr4
Chr5
100 SNPs
25kb
Genotype Proportion
1.00
0.75
0.50
0.25
0.00
0 5 10 15 20 25 30
Mb
B
Genotype Proportion
1.0 1.5 2.0 2.5 3.0 3.5 4.0 4.5 5.0 5.5 6.0 6.5
Mb

**R_042**

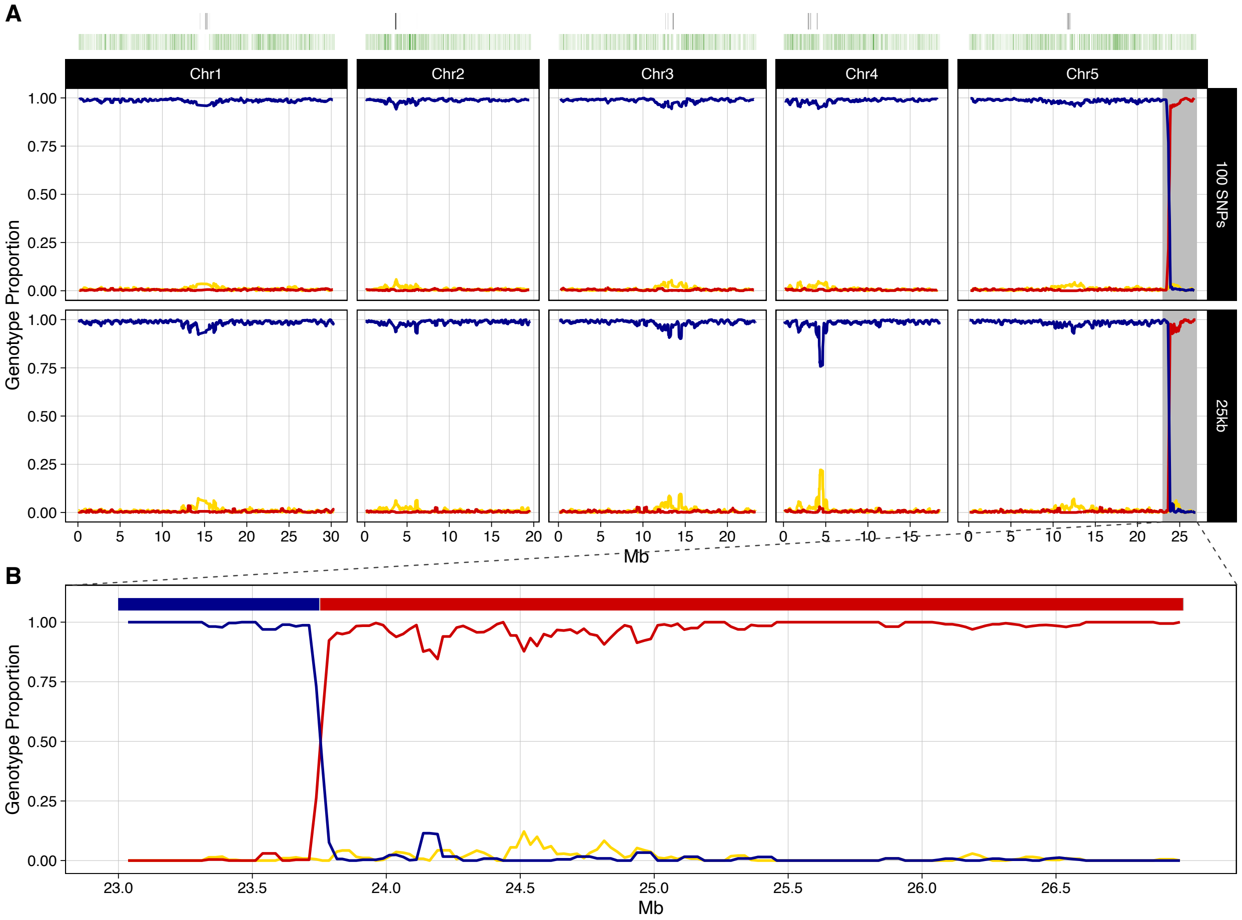
A
Chr1
Chr2
Chr3
Chr4
Chr5
100 SNPs
25kb
Genotype Proportion
1.00
0.75
0.50
0.25
0.00
Mb
B
23.0
23.5
24.0
24.5
25.0
25.5
26.0
26.5

**R_043**

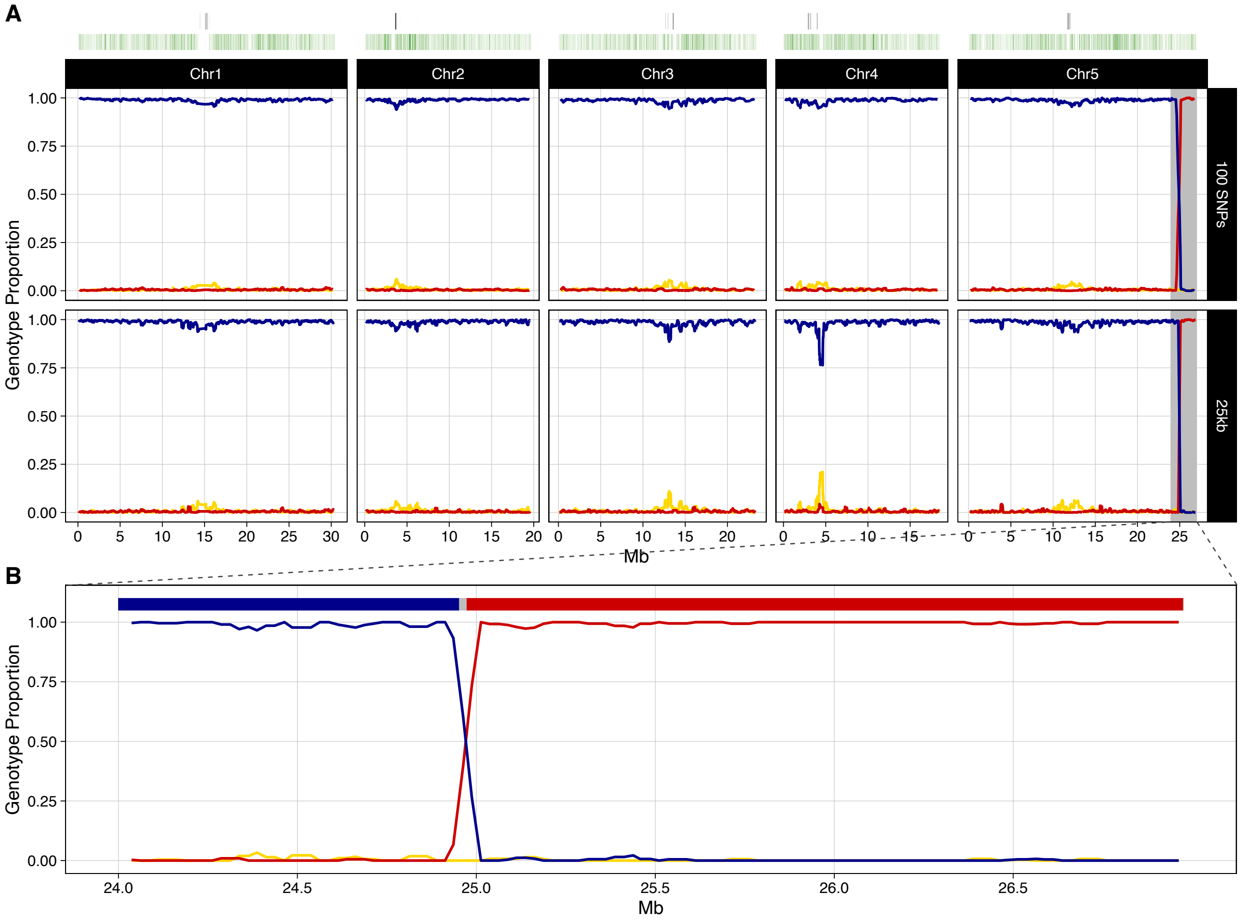
A
Chr1
Chr2
Chr3
Chr4
Chr5
100 SNPs
25kb
Genotype Proportion
1.00
0.75
0.50
0.25
0.00
Mb
B
24.0
24.5
25.0
25.5
26.0
26.5

**R_044**

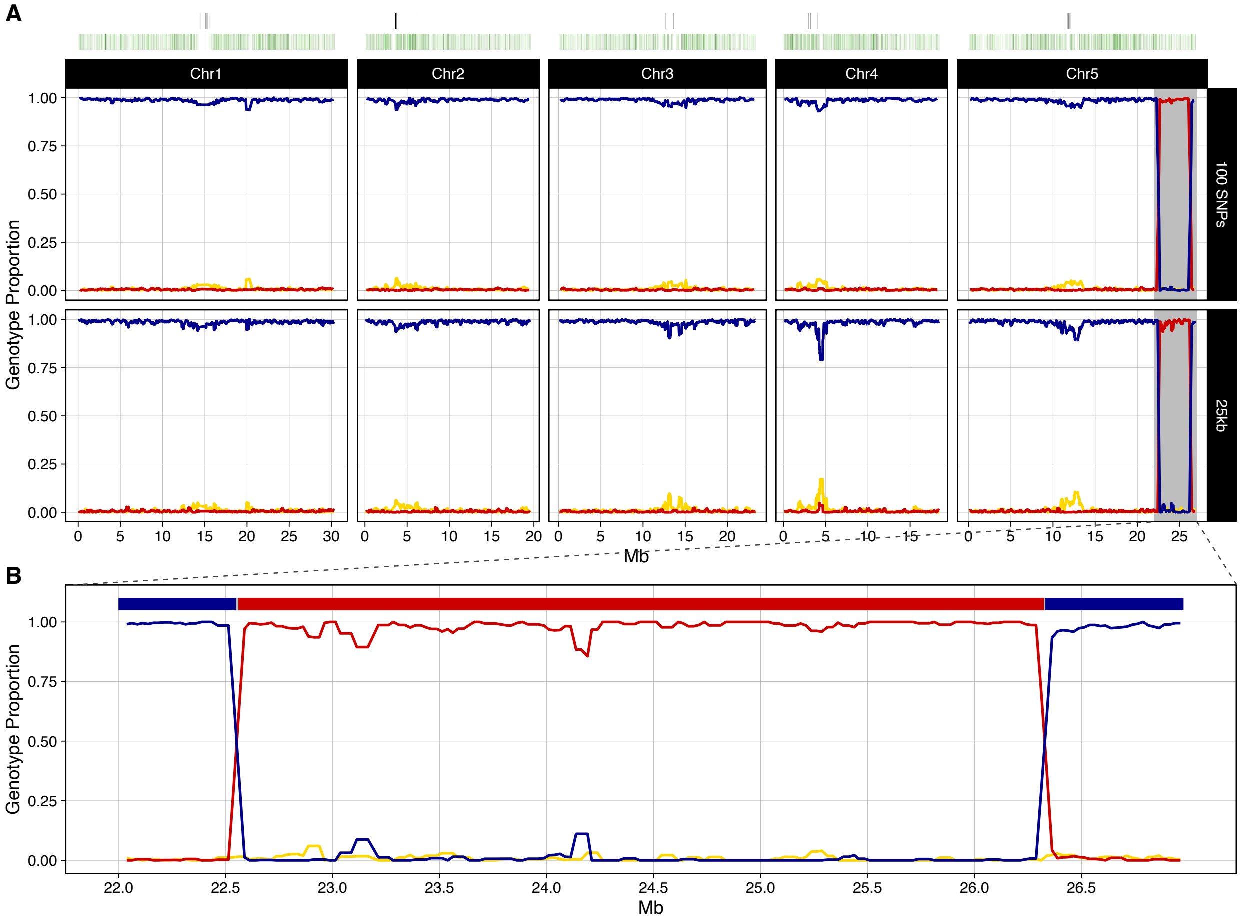

A
Chr1
Chr2
Chr3
Chr4
Chr5
100 SNPs
25kb
Genotype Proportion
Mb
B
Genotype Proportion
Mb

**R_048**

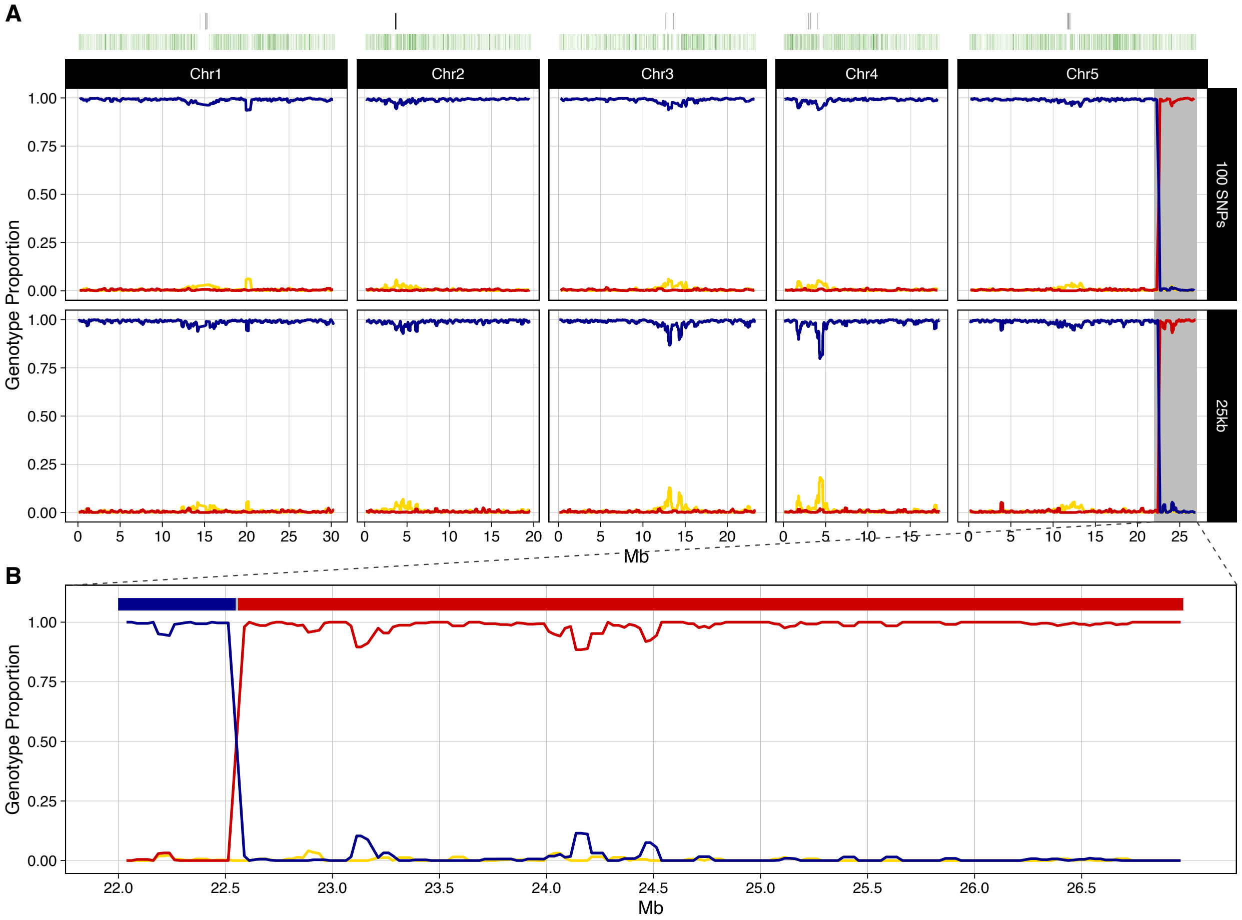

A
Chr1
Chr2
Chr3
Chr4
Chr5
100 SNPs
25kb
Genotype Proportion
1.00
0.75
0.50
0.25
0.00
Mb
B
22.0
22.5
23.0
23.5
24.0
24.5
25.0
25.5
26.0
26.5

**R_049**

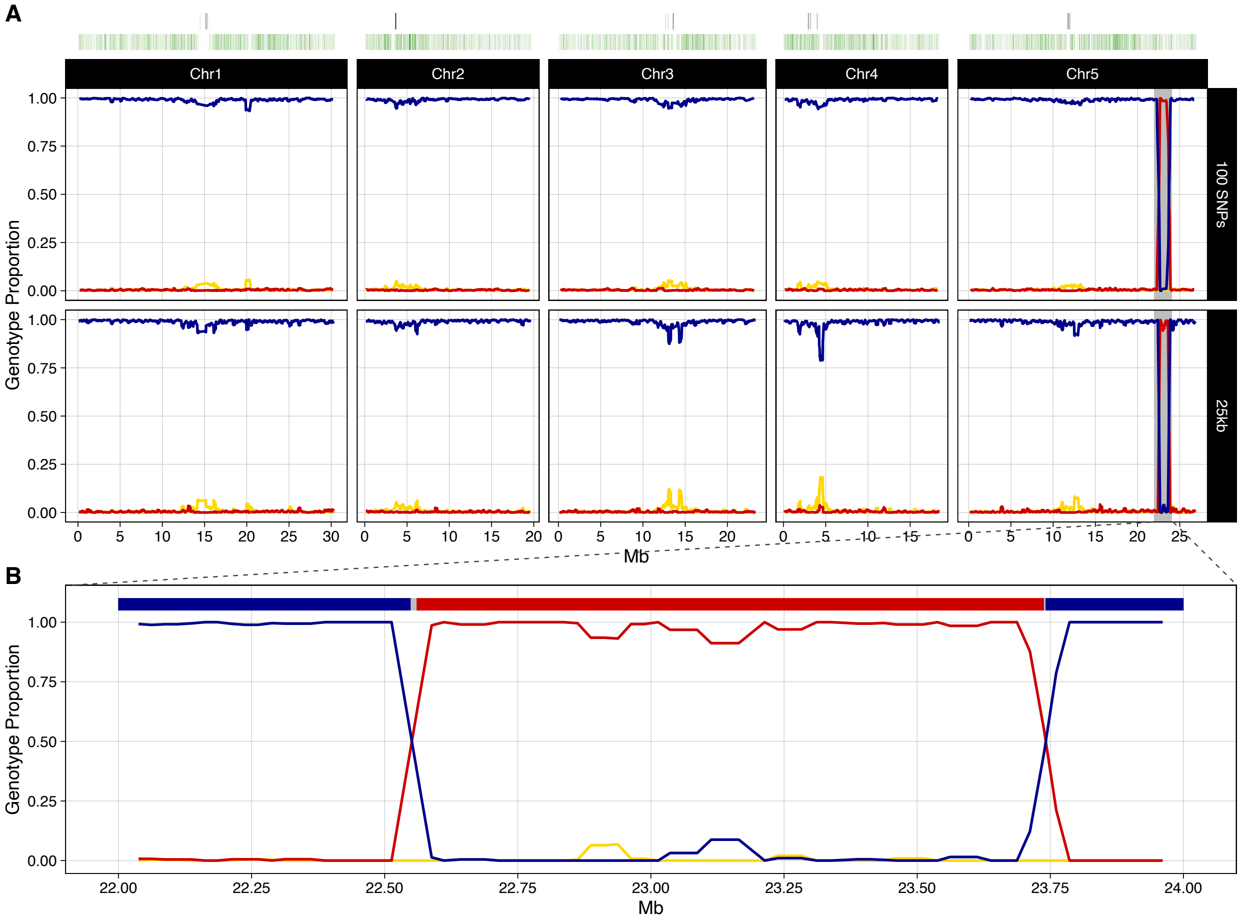
A
Chr1
Chr2
Chr3
Chr4
Chr5
100 SNPs
25kb
Genotype Proportion
Mb
B
Genotype Proportion
Mb

**R_051**

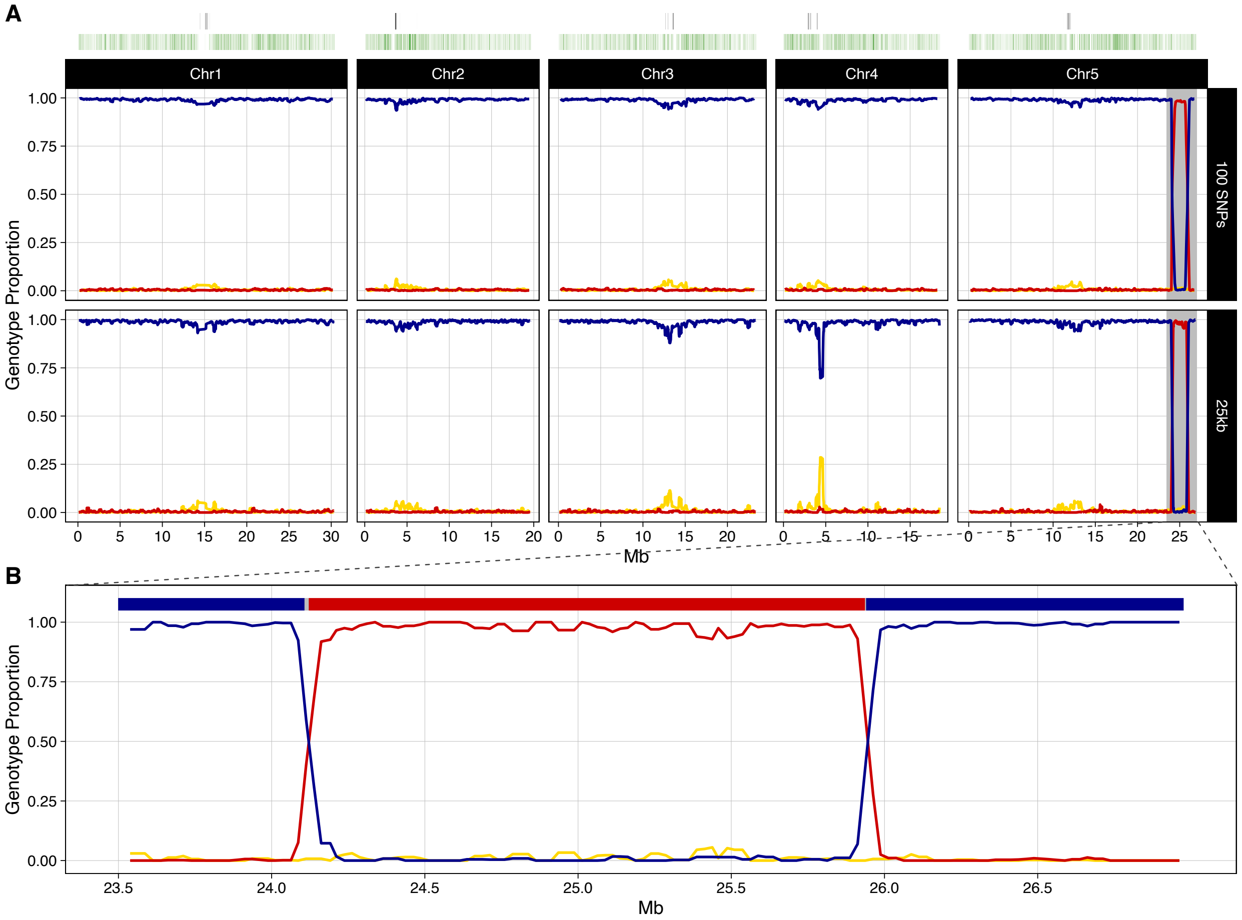
A
Chr1
Chr2
Chr3
Chr4
Chr5
100 SNPs
25kb
Genotype Proportion
1.00
0.75
0.50
0.25
0.00
0
5
10
15
20
25
30
Mb
B
23.5
24.0
24.5
25.0
25.5
26.0
26.5

**R_053**

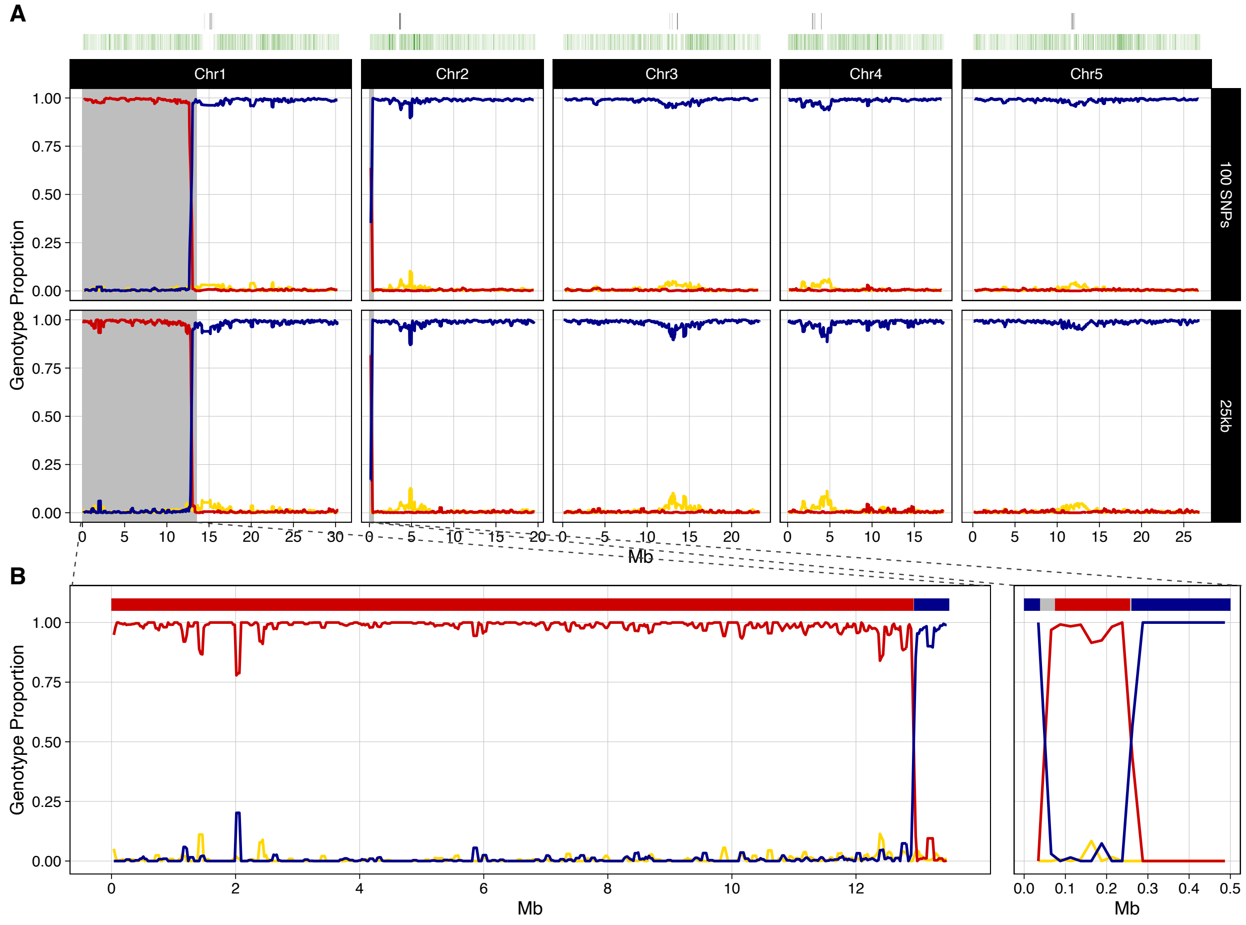
A
Chr1
Chr2
Chr3
Chr4
Chr5
100 SNPs
25kb
Genotype Proportion
1.00
0.75
0.50
0.25
0.00
0 5 10 15 20 25 30
0 5 10 15 20
0 5 10 15 20
0 5 10 15
0 5 10 15 20 25
Mb
B
0 2 4 6 8 10 12
0.0 0.1 0.2 0.3 0.4 0.5

**R_064**

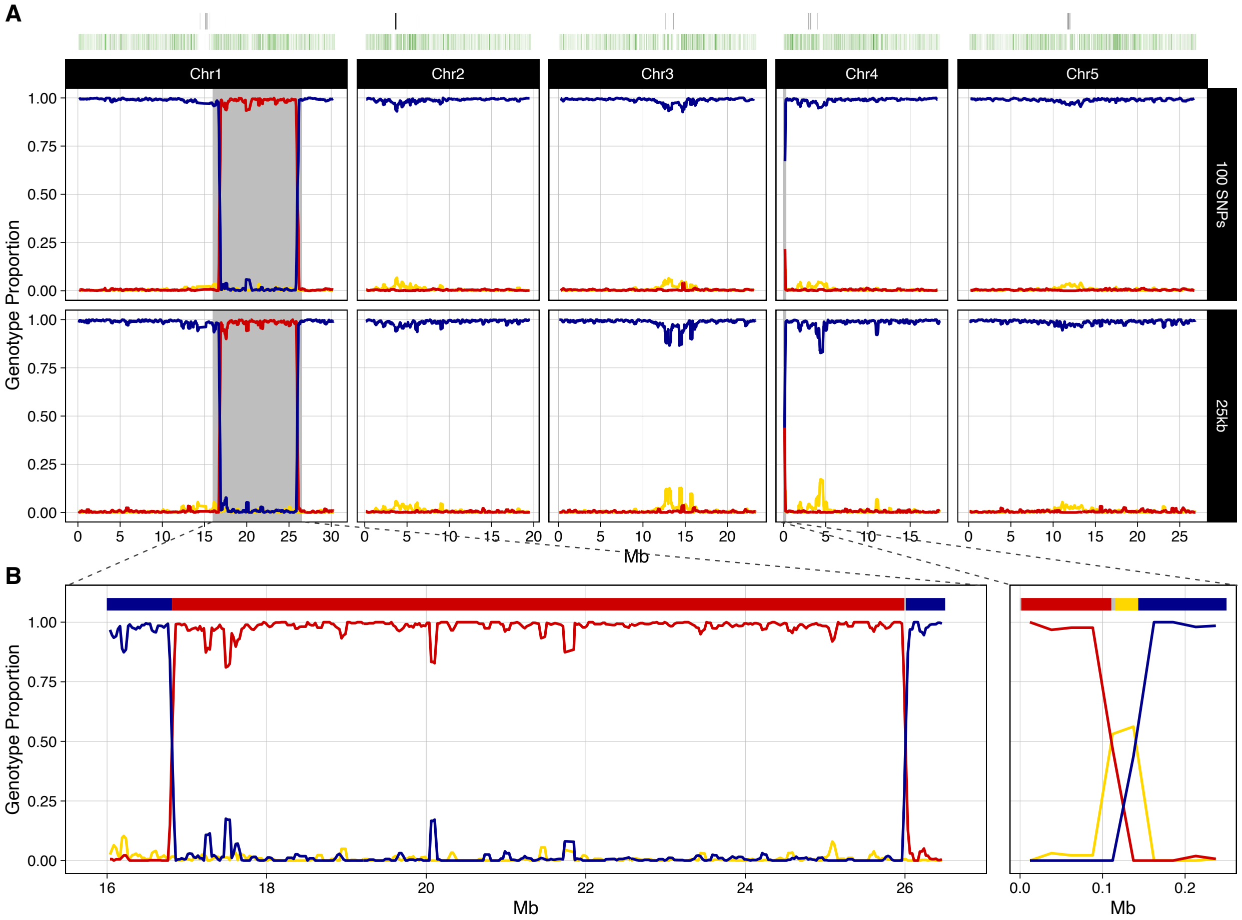
A
Chr1
Chr2
Chr3
Chr4
Chr5
100 SNPs
25kb
Genotype Proportion
1.00
0.75
0.50
0.25
0.00
Mb
B
Genotype Proportion
16
18
20
22
24
26
Mb
0.0
0.1
0.2
Mb

**R_065**

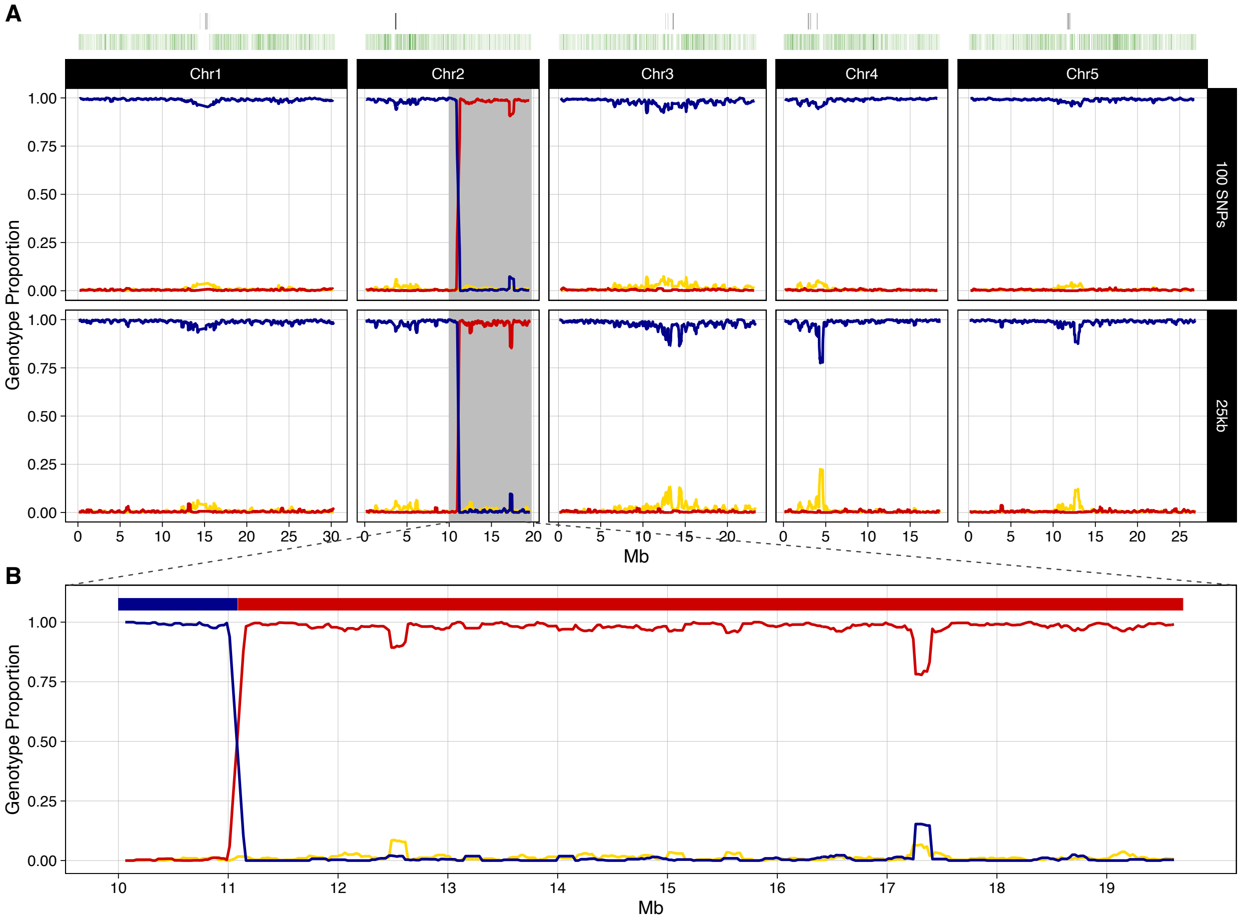
A
Chr1
Chr2
Chr3
Chr4
Chr5
100 SNPs
25kb
Genotype Proportion
1.00
0.75
0.50
0.25
0.00
0
5
10
15
20
25
30
Mb
B
Mb

**R_067**

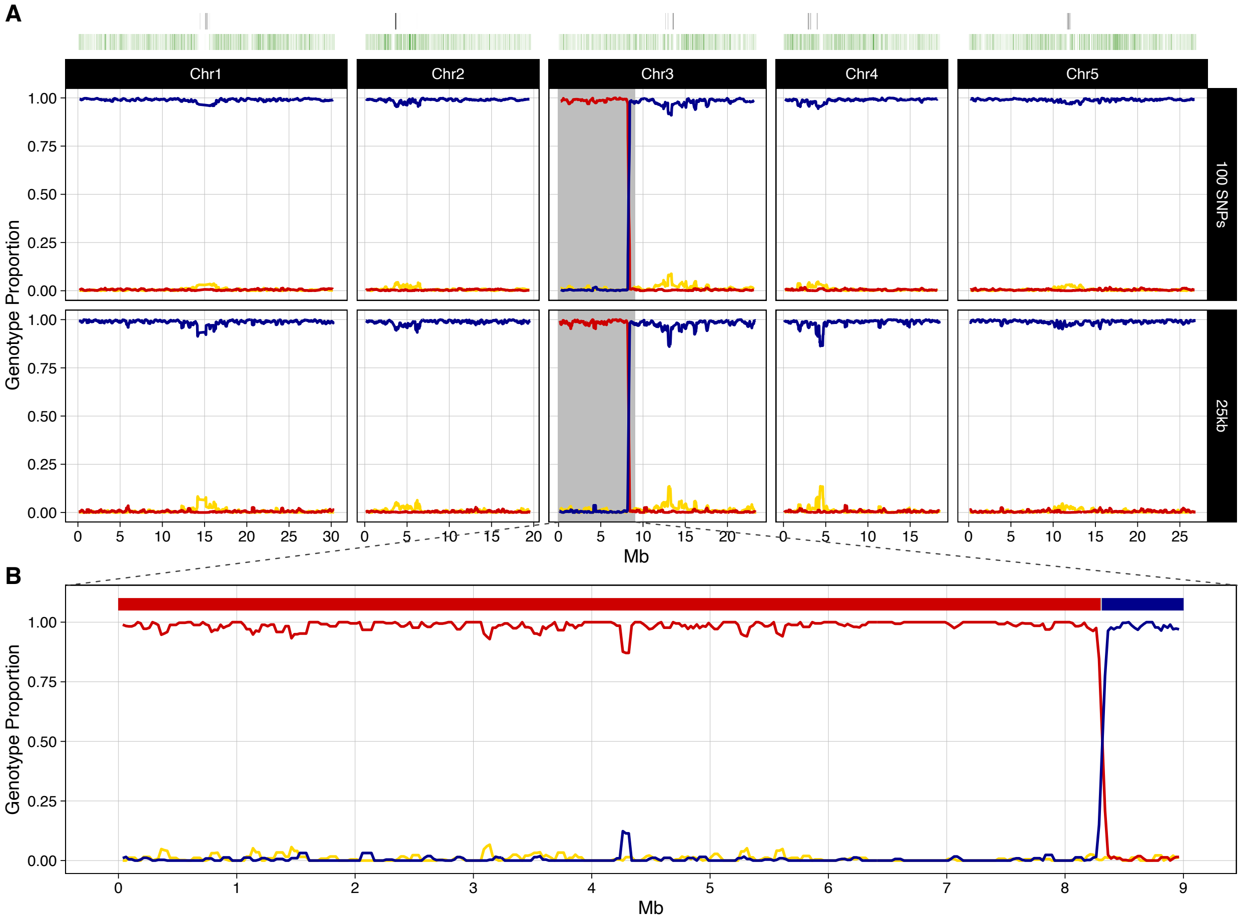
A
Chr1
Chr2
Chr3
Chr4
Chr5
100 SNPs
25kb
Genotype Proportion
Mb
B
Genotype Proportion
Mb

**R_068**

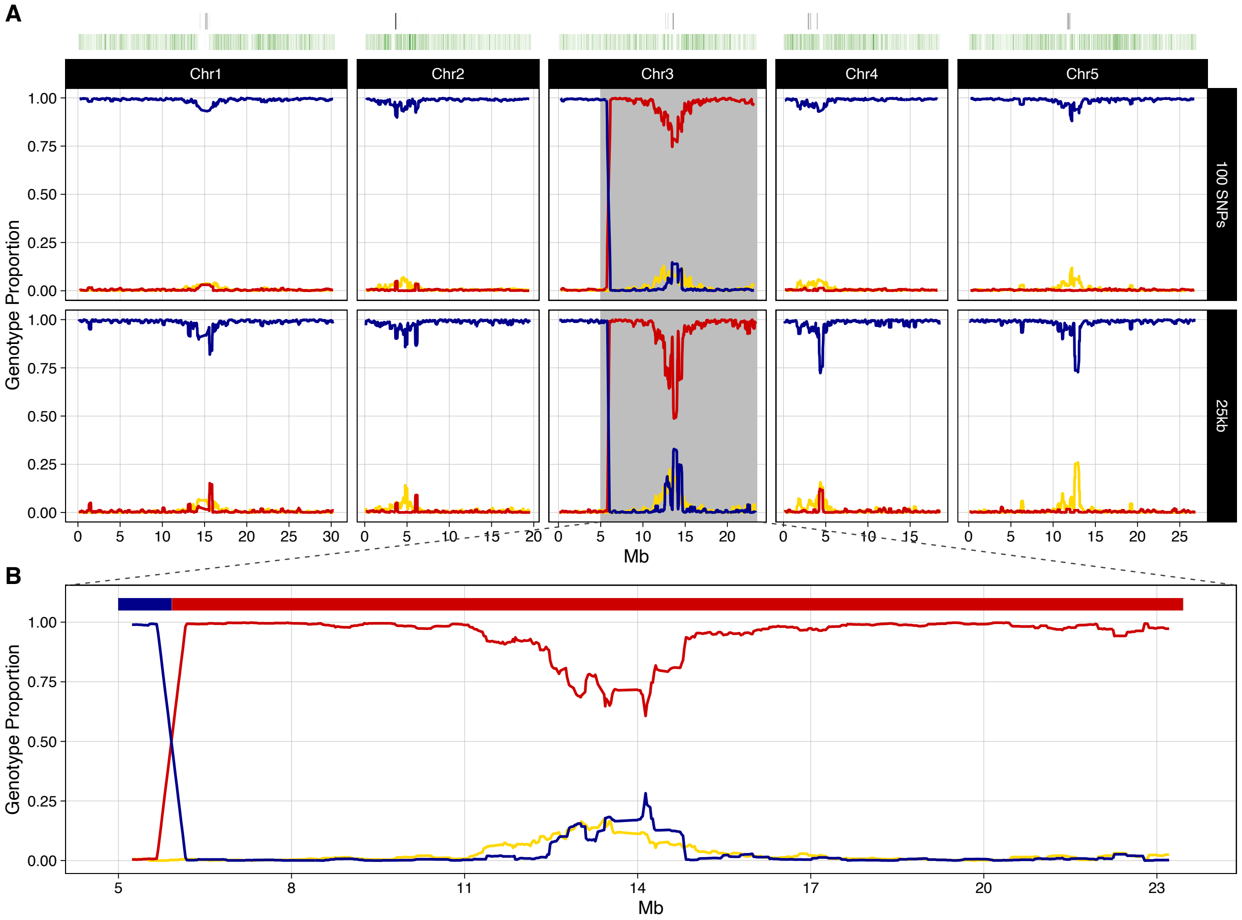
A
Chr1
Chr2
Chr3
Chr4
Chr5
100 SNPs
25kb
Genotype Proportion
1.00
0.75
0.50
0.25
0.00
Mb
B
Genotype Proportion
5
8
11
14
17
20
23
Mb

**R_069**

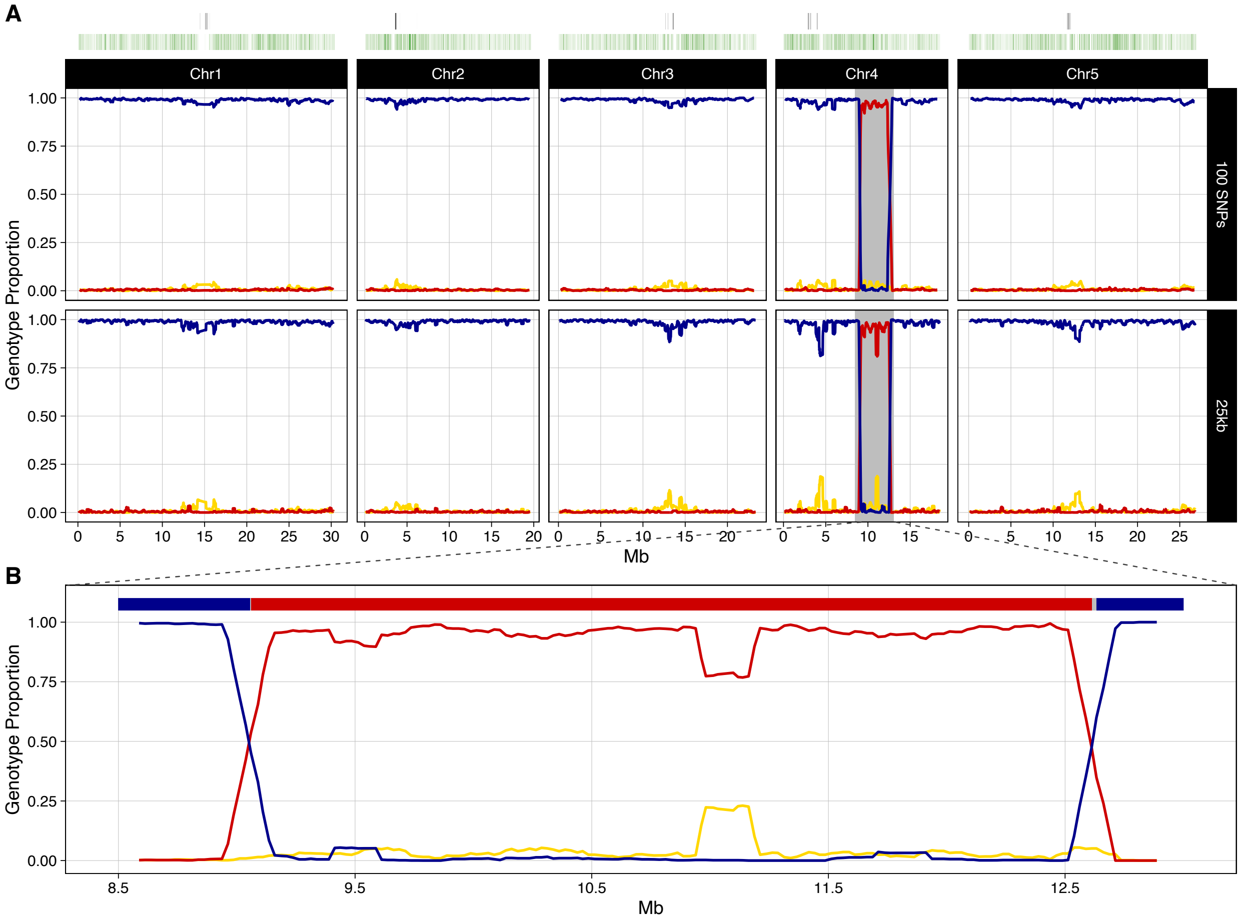

A
Chr1
Chr2
Chr3
Chr4
Chr5
100 SNPs
25kb
Genotype Proportion
1.00
0.75
0.50
0.25
0.00
Mb
B
Genotype Proportion
Mb
8.5
9.5
10.5
11.5
12.5

**R_072**

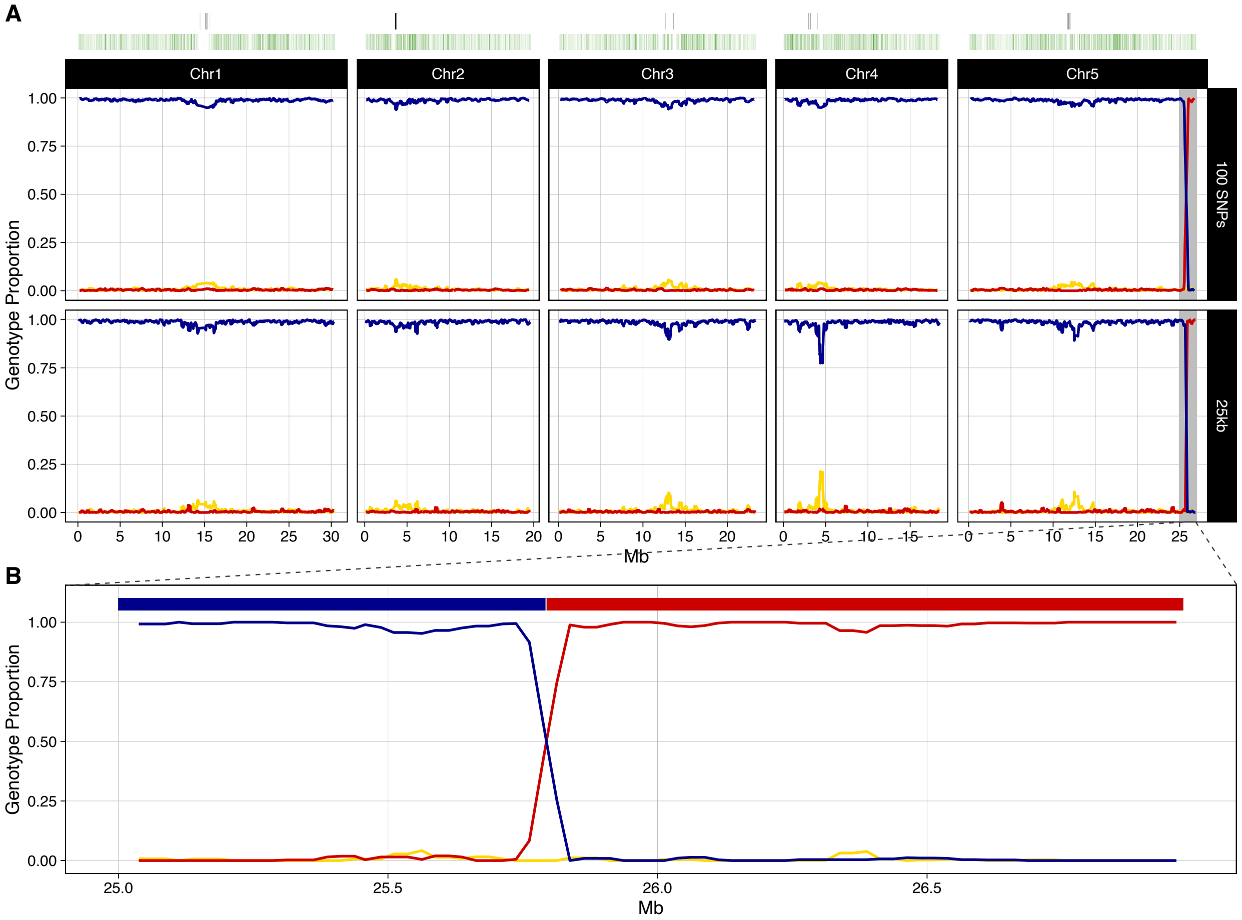

A
Chr1
Chr2
Chr3
Chr4
Chr5
100 SNPs
25kb
Genotype Proportion
1.00
0.75
0.50
0.25
0.00
0 5 10 15 20 25 30
Mb
B
25.0
25.5
26.0
26.5

**R_095**

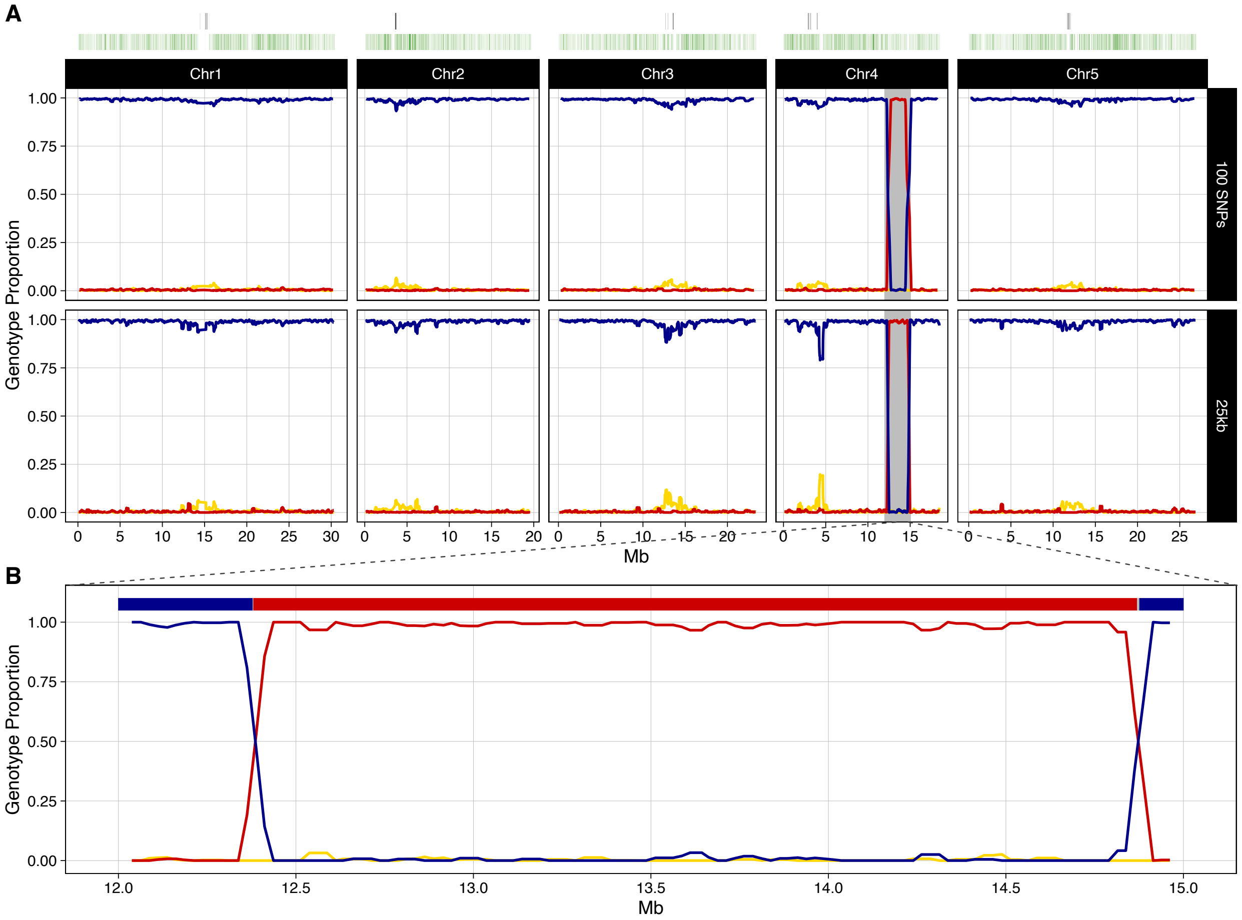
A
Chr1
Chr2
Chr3
Chr4
Chr5
100 SNPs
25kb
Genotype Proportion
1.00
0.75
0.50
0.25
0.00
0 5 10 15 20 25 30
Mb
B
Genotype Proportion
12.0
12.5
13.0
13.5
14.0
14.5
15.0
Mb

**R_103**

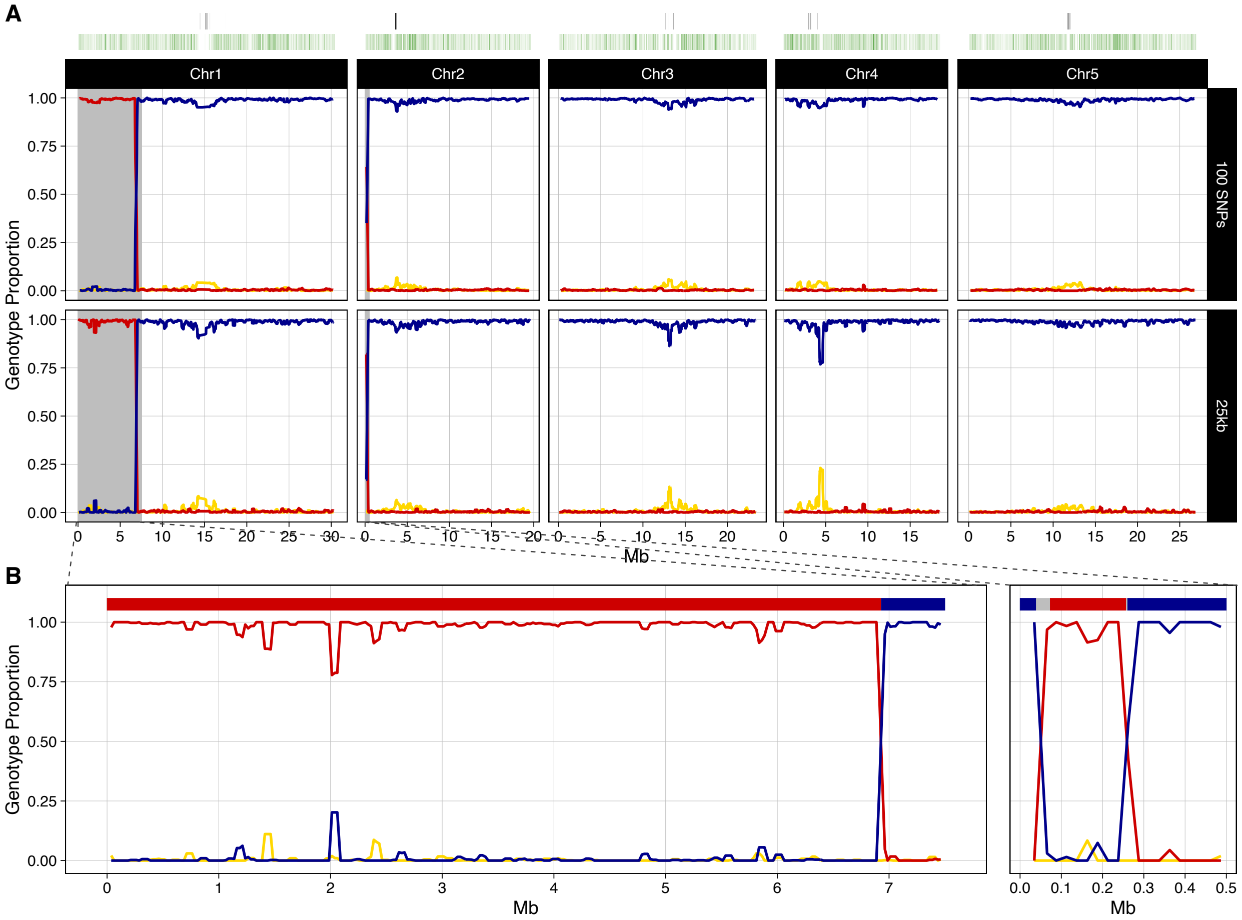
A
Chr1
Chr2
Chr3
Chr4
Chr5
100 SNPs
25kb
Genotype Proportion
1.00
0.75
0.50
0.25
0.00
Mb
B
Genotype Proportion
Mb
Mb

**R_112**

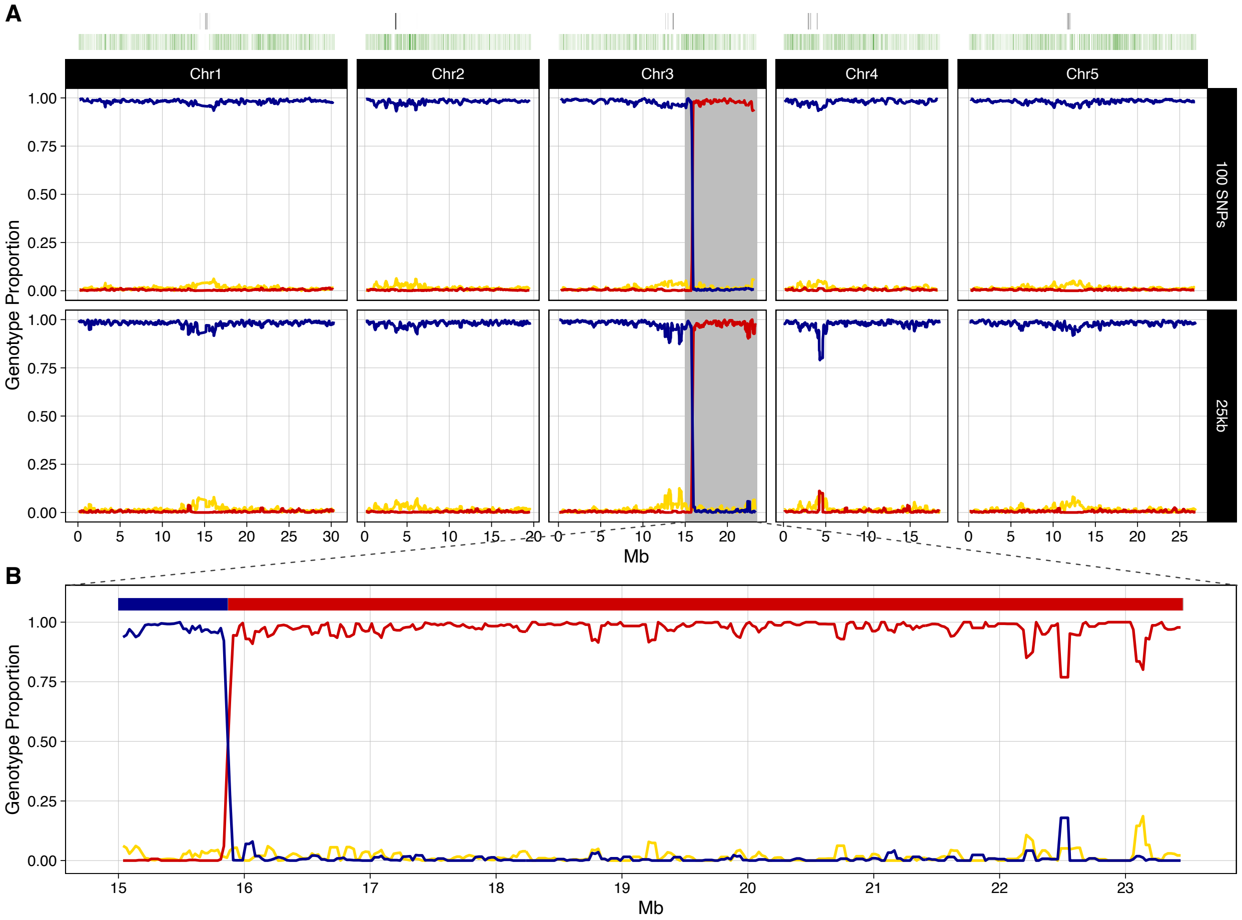
A
Chr1
Chr2
Chr3
Chr4
Chr5
100 SNPs
25kb
Genotype Proportion
1.00
0.75
0.50
0.25
0.00
Mb
B
Genotype Proportion
Mb

**R_113**

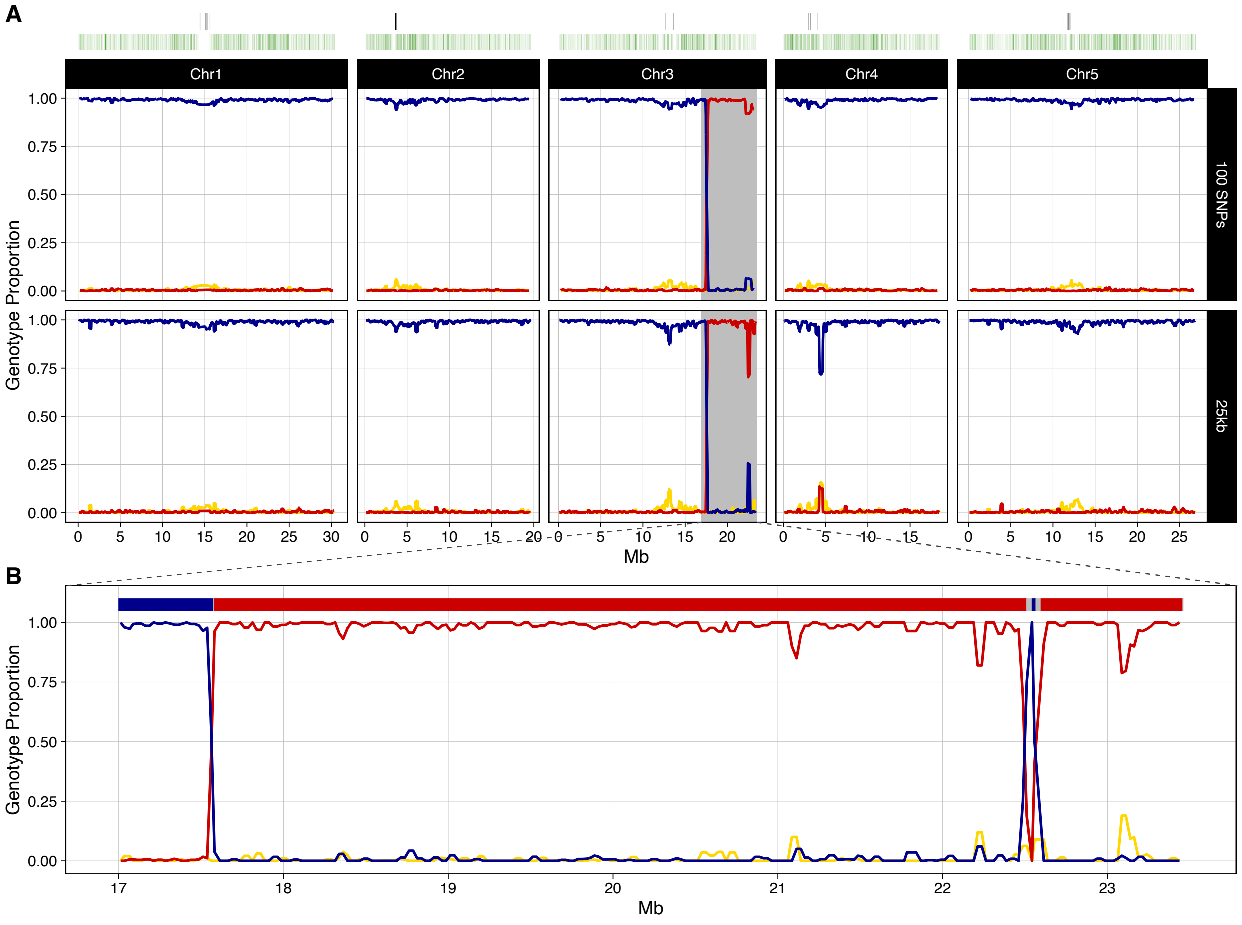
A
Chr1
Chr2
Chr3
Chr4
Chr5
100 SNPs
25kb
Genotype Proportion
1.00
0.75
0.50
0.25
0.00
0
5
10
15
20
25
30
Mb
B
17
18
19
20
21
22
23

**R_114**

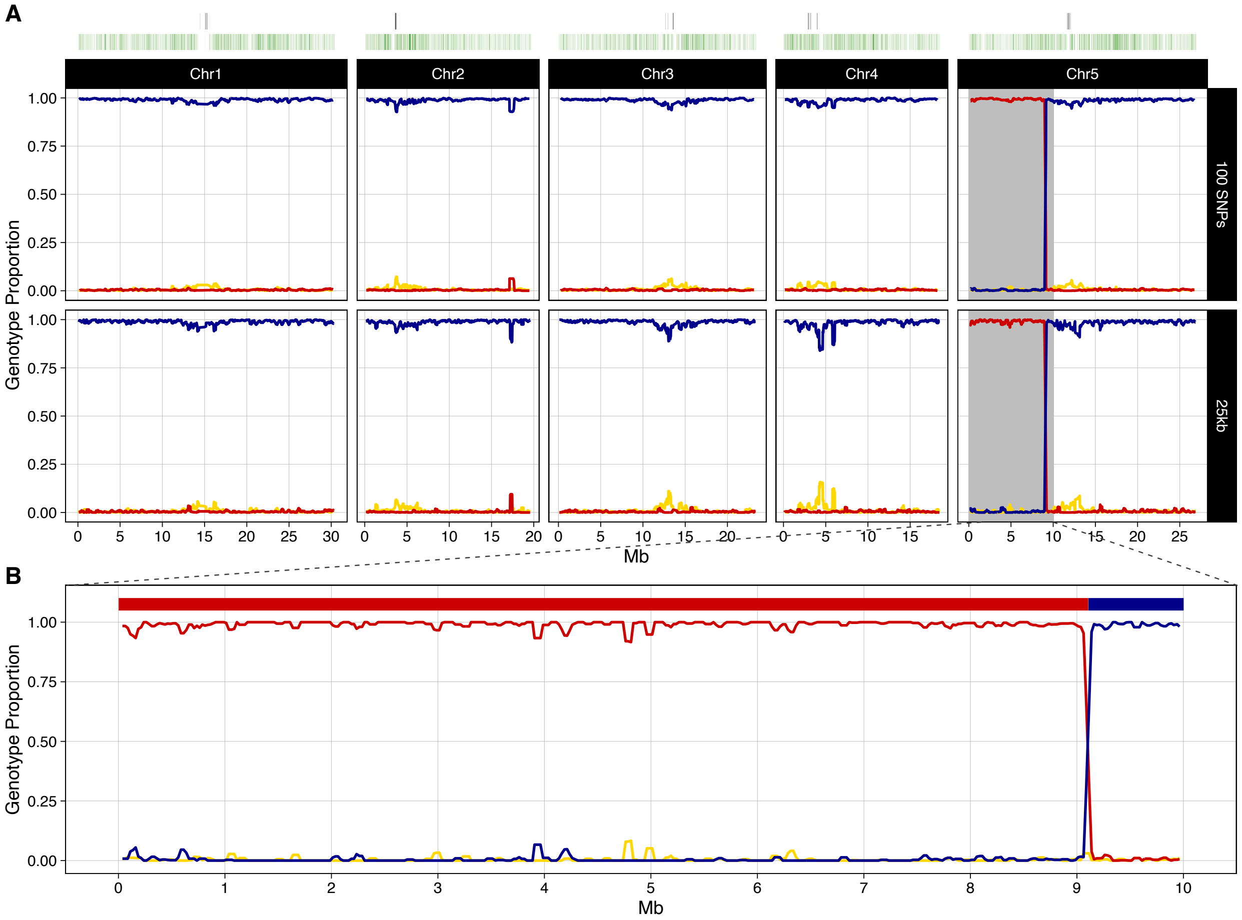
A
Chr1
Chr2
Chr3
Chr4
Chr5
100 SNPs
25kb
Genotype Proportion
1.00
0.75
0.50
0.25
0.00
Mb
B
Genotype Proportion
Mb

**R_119**

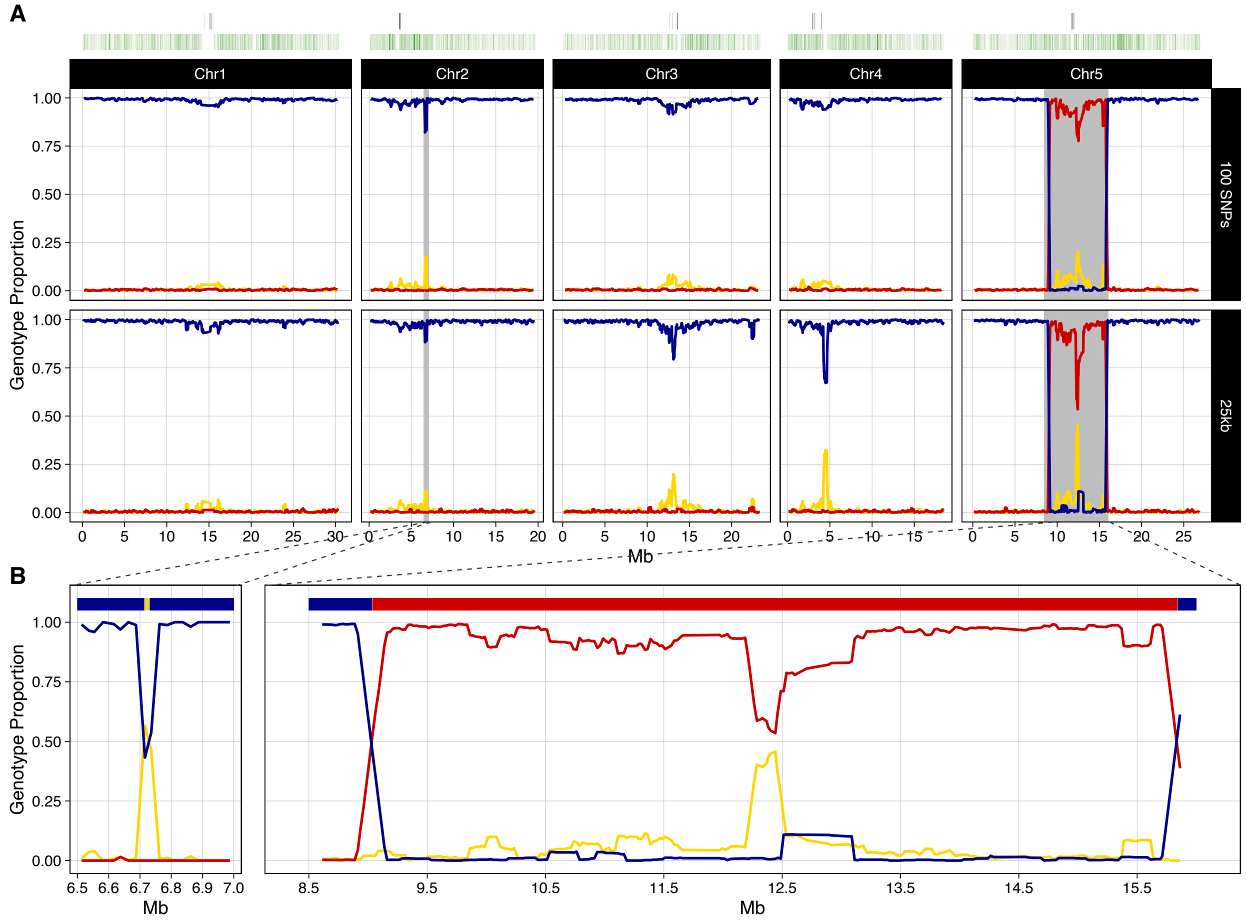
A
Chr1
Chr2
Chr3
Chr4
Chr5
100 SNPs
25kb
Genotype Proportion
1.00
0.75
0.50
0.25
0.00
0
5
10
15
20
25
30
Mb
B
6.5
6.6
6.7
6.8
6.9
7.0
8.5
9.5
10.5
11.5
12.5
13.5
14.5
15.5

**R_144**

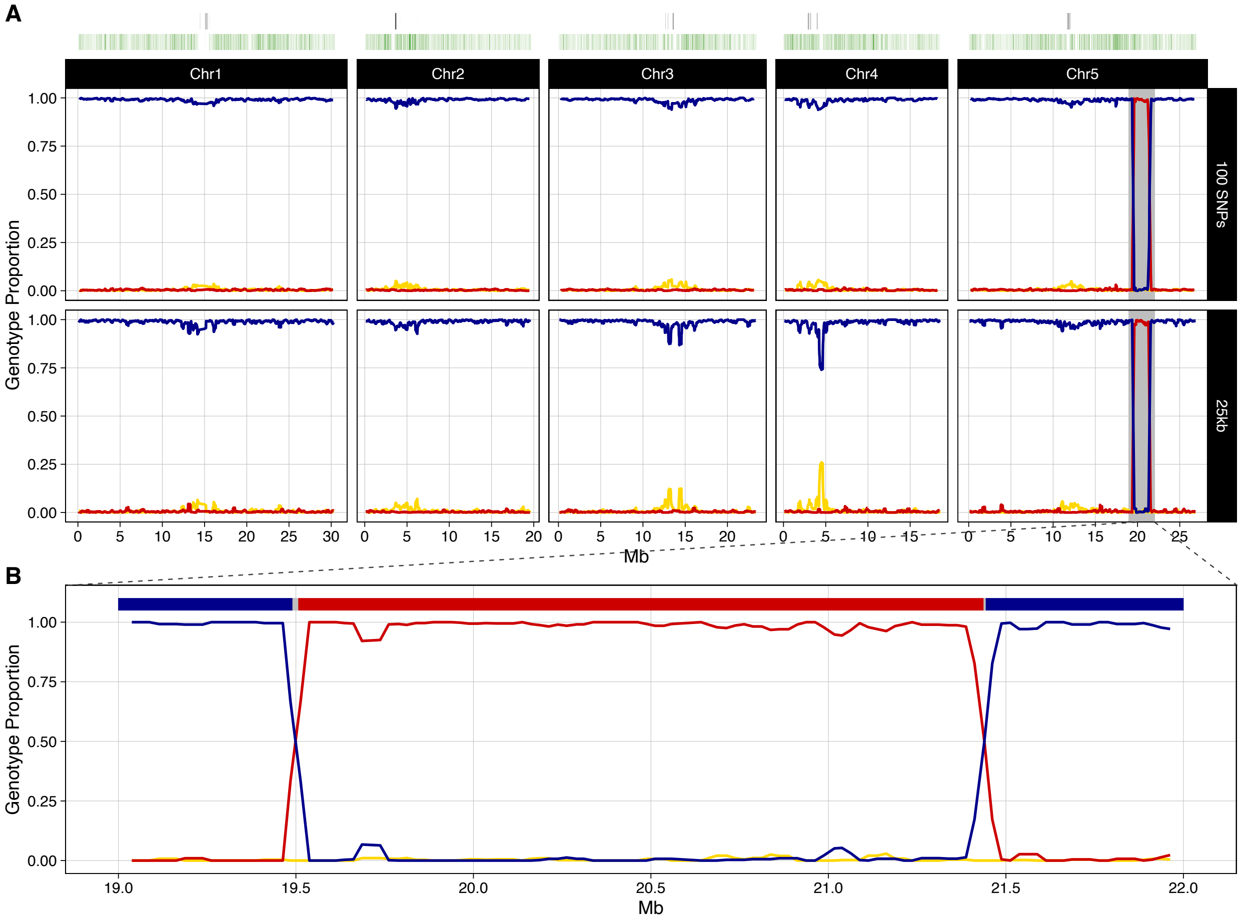
A
Chr1
Chr2
Chr3
Chr4
Chr5
100 SNPs
25kb
Genotype Proportion
1.00
0.75
0.50
0.25
0.00
Mb
B
19.0
19.5
20.0
20.5
21.0
21.5
22.0

**R_145**

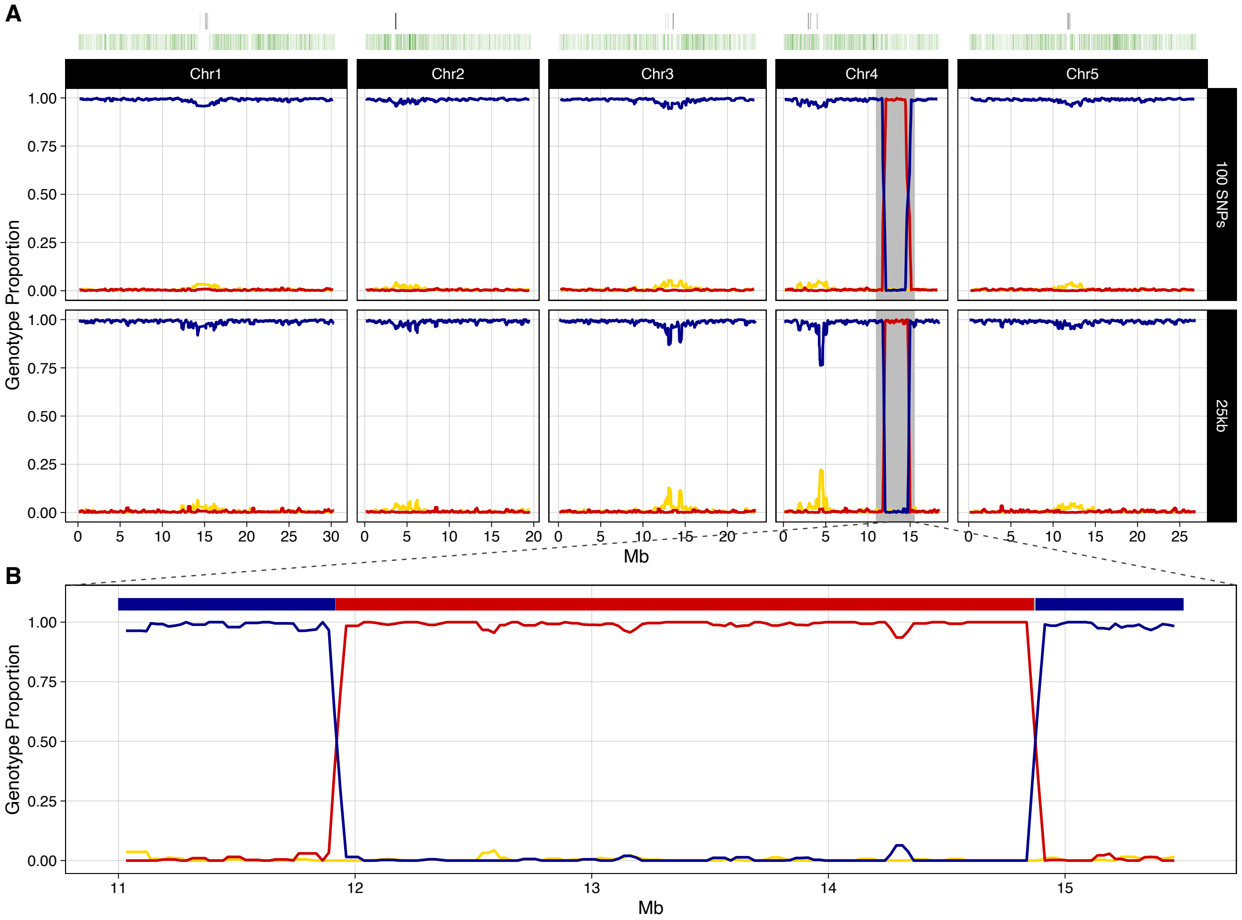
A
Chr1
Chr2
Chr3
Chr4
Chr5
100 SNPs
25kb
Genotype Proportion
1.00
0.75
0.50
0.25
0.00
Mb
B
11
12
13
14
15

**R_147**

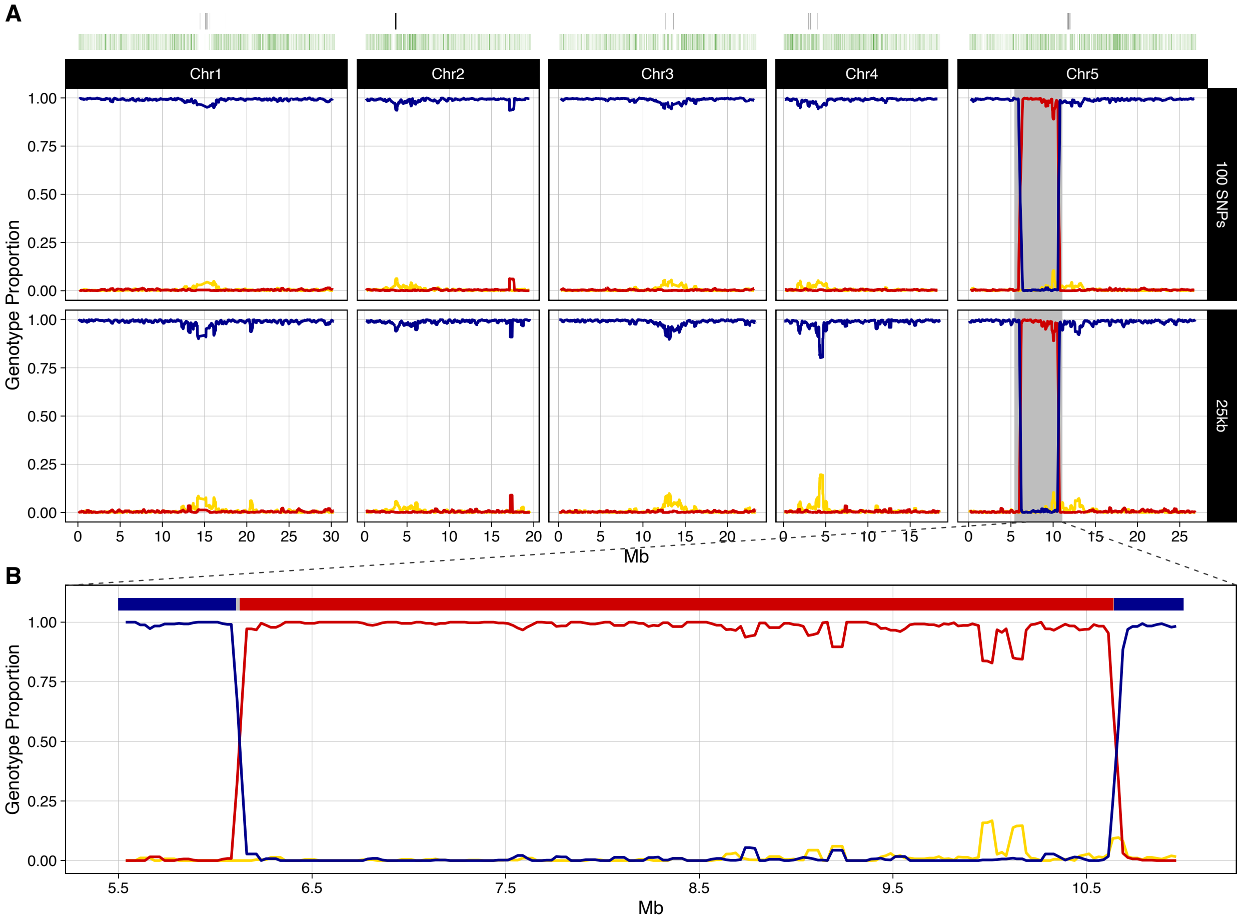

## R_153

**Figure S2. Detailed genotyping of all NILs. (A)** Rolling average of genotype proportion (IT: red, SW: blue, heterozygous: gold) in non-overlapping 100 SNP windows and non-overlapping 25kb windows across the 5 chromosomes of the *A. thaliana* genome. Heatmaps above show the density of centromeric repeats (grey) and density of diagnostic SNP markers (green) across the genome. **(B)** Zoomed in view of the introgressed region of interest, rolling average of genotype proportion (IT: red, SW: blue, heterozygous: gold) in non-overlapping 25kb windows, with a summary haplotype map above. Crossover regions between SNP markers (grey) are also shown.
